# Endosomal escape of delivered mRNA from endosomal recycling tubules visualized at the nanoscale

**DOI:** 10.1101/2020.12.18.423541

**Authors:** Prasath Paramasivam, Christian Franke, Martin Stöter, Andreas Höijer, Stefano Bartesaghi, Alan Sabirsh, Lennart Lindfors, Marianna Yanez Arteta, Anders Dahlén, Annette Bak, Shalini Andersson, Yannis Kalaidzidis, Marc Bickle, Marino Zerial

## Abstract

Delivery of exogenous mRNA using lipid nanoparticles (LNP) is a promising strategy for therapeutics. However, a bottleneck remains the poor understanding of the parameters that correlate with endosomal escape vs. cytotoxicity. To address this problem, we compared the endosomal distribution of six LNP-mRNA formulations of diverse chemical composition and efficacy, similar to those employed in mRNA-based vaccines, in primary human adipocytes, fibroblasts and HeLa cells. Surprisingly, we found that total uptake is not a sufficient predictor of delivery and different LNP vary considerably in endosomal distributions. Prolonged uptake impaired endosomal acidification, a sign of cytotoxicity, and caused mRNA to accumulate in compartments defective in cargo transport and unproductive for delivery. In contrast, early endocytic/recycling compartments have the highest probability for mRNA escape. By super-resolution microscopy we could resolve single LNP-mRNA within sub-endosomal compartments and capture events of mRNA escape from endosomal recycling tubules. Our results change the view of the mechanisms of endosomal escape and define quantitative parameters to guide the development of mRNA formulations towards higher efficacy and lower cytotoxicity.

## Introduction

In recent years, RNAs have emerged as potentially powerful therapeutics (Coutinho, Matos et al. 2019) to inhibit gene expression (splicing modulators, siRNAs and antisense oligonucleotides), express proteins (mRNA) or gene editing (CRISPR/Cas9 system). An increasing number of RNA-based therapeutics have proven effective for clinical treatment (Akinc, Maier et al. 2019, Dammes and Peer 2020, Sahin, Muik et al. 2020). More recently, optimization of chemical and physical properties have focused the attention on mRNA-based therapeutics, including for vaccines (Feldman, Fuhr et al. 2019, Kose, Fox et al. 2019). Major improvements towards clinical application have come from chemical modifications of RNAs that increase stability and reduce immunogenicity. Nevertheless, efficacy remains a crucial challenge due to limited or poor delivery (Kowalski, Rudra et al. 2019).

Lipid nanoparticles (LNP) are currently the non-viral RNA delivery platform of choice (Reichmuth, Oberli et al. 2016, Kowalski, Rudra et al. 2019). LNP have different chemical compositions and show vastly different delivery efficiency, toxicity and immunological liability. The mechanistic basis for such differences is unclear. Since delivery is a multi-step process (Wittrup and Lieberman 2015), attempting structure/activity relation without understanding the underlying mechanisms can yield complex and contradicting results, as the outcome of experiments is influenced by the non-linear combination of the various steps. Besides endocytosis, a major challenge remains the ability of RNA to cross the endosomal membrane(Pei and Buyanova 2019). Ineffective escape from endosomes imposes higher dosage, thus causing toxicity. Toxicity is due to both cell autonomous, i.e. cytotoxicity, as well as non-cell autonomous effects, e.g. inflammation (Reichmuth, Oberli et al. 2016, Sato, Matsui et al. 2017). The reasons for cytotoxicity are diverse, comprising oxidative stress and apoptosis(Ahmad, Wahab et al. 2020). Whether they include alterations of the endosomal system is unknown.

The precise site(s) and mechanisms whereby LNP help mRNA escape from the endosomal lumen are to date mysterious. Escape efficiency arguably depends on the distribution of LNP in various subcellular compartments, of which only a selected few are poised to macromolecule escape. Previous studies have yielded contradictory results on LNP internalization, endosomal distribution and escape of RNA (siRNA). Whereas we and others (Gilleron, Querbes et al. 2013, Wittrup, Ai et al. 2015) reported that escape is restricted to an early endosomal compartment prior to conversion into late endosomes (Rink, Ghigo et al. 2005), another study claimed that escape occurs mainly from late endosomes where LNP accumulate (Sahay, Querbes et al. 2013). However, this conclusion was based on perturbations (drugs and gene downregulation) that induce pleiotropic effects, making the data difficult to interpret. Furthermore, the complexity of the endosomal network cannot be underestimated. It consists of populations of organelles that are sub-compartmentalized (Sönnichsen, De Renzis et al. 2000, Franke, Repnik et al. 2019), dynamically exchange cargo and trafficking machinery, and change in size and position over time, calling for a thorough quantitative and high resolution analysis to interpret their precise identities. Besides the endosomal compartments granting RNA escape, the underlying mechanisms remain mysterious. Live cell imaging has shown that lipoplexes deliver siRNAs by endosome bursting, but this mechanism does not apply to LNP (Gilleron, Querbes et al. 2013, Wittrup, Ai et al. 2015). Despite claims of membrane fusion and imaging mRNA escape into the cytoplasm (Miao, Lin et al. 2020), the sensitivity and resolution of conventional microscopic techniques are insufficient to visualize escape of a small number of mRNAs. For this, single molecule techniques are essential.

To address this critical problem, we aimed to determine whether differences in LNP-mediated mRNA (LNP-mRNA) delivery efficacy may originate from variations in 1) uptake and/or 2) transport to endosomal sub-compartments with higher probability to escape than others. As yet, LNP have predominantly been investigated for intravenous or intramuscular administration. Subcutaneous administration in adipose tissue opens the possibility of patient self-administration and, hence, long-term chronic treatment that could enable mRNA as a novel modality for protein replacement or regenerative therapies. Therefore, we performed a systematic comparison of six mRNA-containing LNP with different cationic lipids including MOD5, an analogue of the SM-102 lipid used in Moderna “mRNA-1273” SARS-CoV-2 vaccine (Sabnis, Kumarasinghe et al. 2018), in primary human adipocytes, fibroblasts and HeLa cells to identify candidate parameters diagnostic for efficacy of delivery.

## Results

### LNP-mRNA internalization is necessary but not sufficient to predict delivery efficacy

We performed a comparative analysis on uptake and endosomal distribution of mRNA encoding eGFP formulated in six distinct LNP with similar size distribution (54-73 nm) and mRNA content (Supplementary Table 1) but chosen on the basis of various chemical structures of the cationic lipid (Extended Data Fig. 1a), efficacy, as evaluated by intensity of eGFP expression (Fig. 1a, Extended Data Fig. 2a) as well as toxicity (Ex. MC3 vs. MOD5, or L319) (Maier, Jayaraman et al. 2013, Sabnis, Kumarasinghe et al. 2018). We used primary human adipocytes, as a relevant cell model for mRNA administered by subcutaneous injection and detected the mRNA with single molecule fluorescence in situ hybridization (smFISH), unless otherwise mentioned. We applied image analysis methods to discriminate between adipocytes and fibroblasts in the same culture. Measurement of uptake kinetics indeed showed that internalization and trafficking vary between LNP-mRNA (Fig. 1b). Interestingly, although in most cases uptake correlated with transfection efficacy (e.g. MC3 vs. L319), this was not a strict rule. L608 demonstrated highest eGFP expression despite moderate LNP-mRNA uptake, compared with e.g. MC3 and ACU5 (Fig. 1b). Besides uptake efficiency, efficacy may depend on qualitative and/or quantitative differences in endosomal distributions.

**Figure 1:**
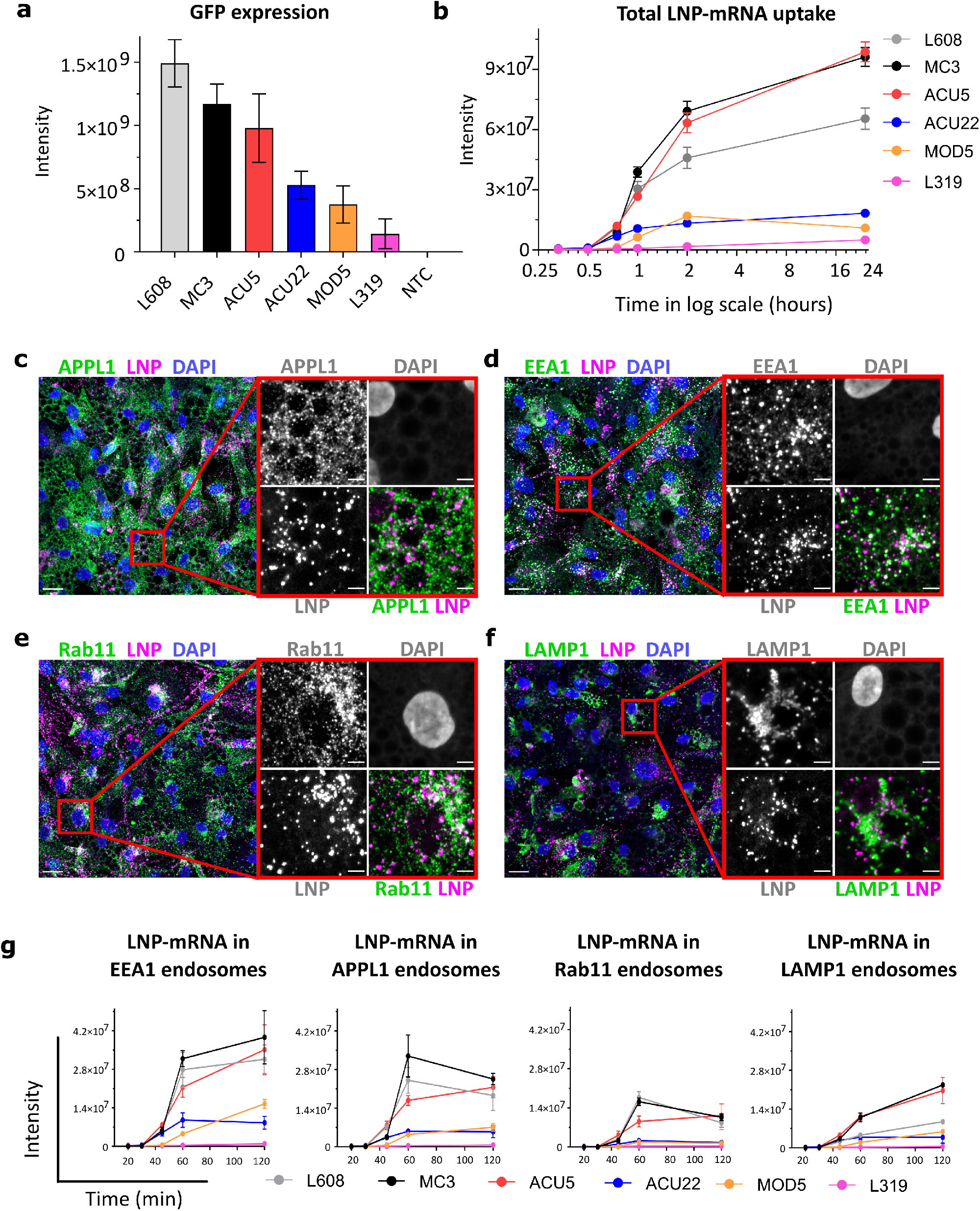
Comparative analysis of activity, endocytic uptake and endosomal distribution of six LNP-mRNA in primary human adipocytes. **a**) Cells were incubated with various LNPs (1.25ng/µL) formulated with eGFP-mRNA, fixed, and imaged after 24h. The graph illustrates GFP expression for the indicated LNP-mRNA. N=3 independent experiments (mean ± s.e.m). (**b**) Cells incubated with LNP-mRNA as described above were fixed at the given time point, processed for smFISH to fluorescently label mRNA and imaged by fluorescence microscopy. The quantification shows that LNP-mRNA uptake generally correlates to the GFP expression efficacy of LNP formulations displayed in panel **a** (Ex. MC3 Vs L319). (**c-f**) Representative images of human primary adipocytes incubated with L608 LNP-mRNA for 2h, immunostained with antibodies against endosomal markers (in green): APPL1 (c), EEA1 (d), Rab11 (e) and LAMP1 (f), exogenous mRNA detected by smFISH (labeled as LNP in the figure panel) and nuclei by DAPI. The magnified representative area is presented with split and merged color images. Bars indicate 20µm in the overview and 5µm in the magnified images. (**g**) Kinetics and endosomal distribution of the different LNP-mRNA incubated with cells as described in (c-f). N=3 replicates. (mean ± s.e.m).

**Figure 2:**
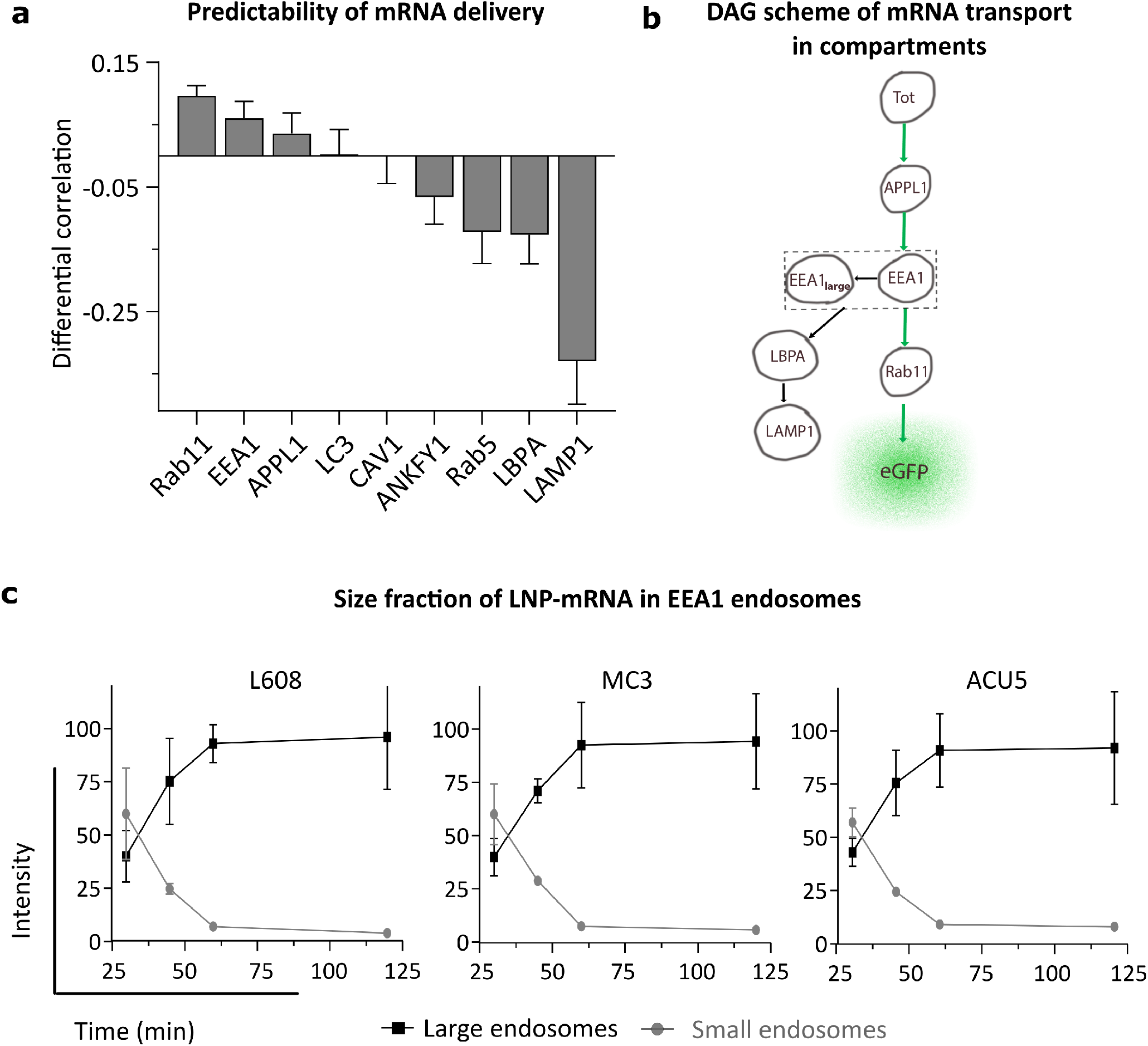
Differential correlation and size distribution kinetics analysis reveal that Rab11, small EEA1-and APPL1-positive endosomes are more favorable than other endosomal compartments for mRNA delivery. **(a)** The graph shows the differential correlation between LNP-mRNA distribution in 9 endosomal compartments to eGFP expression in human primary adipocytes (see **Methods**). Three compartments Rab11, EEA1 and APPL1 have correlation above total uptake and therefore are potential hotspots of mRNA escape, whereas e.g. Lamp1 is not. **(b) S**chematic representation of LNP-mRNA transport through endosomal compartments reconstructed based on the DAG differential correlation (panel **a**) and size fraction analysis (panel **c**). The total LNP-mRNA uptake is represented by node “Tot” which is upstream of the various endosomal compartments (represented by the indicated nodes) where cargo is sequentially distributed to. The Rab11 node is located close to the node eGFP because of its high correlation of LNP-mRNA distribution to eGFP expression (see **Methods**). Whereas the node EEA1 is upstream because it consists of small EEA1 LNP-mRNA endosomes that are productive for delivery, the large “arrested” (EEA1large) endosomes are less/unproductive for LNP-mRNA delivery. **(c)** Quantification of LNP-mRNA accumulation in EEA1 endosomes based on endosome size and intensity. The graph illustrates that the fraction of large LNP-mRNA endosomes (object diameter > 1µm) increases substantially whereas the smaller ones (object diameter = up to 0.75µm) decrease over time. Percentage decrease of small LNP-mRNA endosomes, L608 = 60.0% ± 14 to 5.7 ± 0.4; MC3 = 59.9% ± 21 to 4 ± 0.5; ACU5 = 57% ± 7 to 8% ± 1.0 Percentage increase of large LNP-mRNA endosomes, L608 = 40.0% ± 9 to 94.2 ± 22; MC3 = 40.0% ± 12 to 96.0 ± 24; ACU5 = 42.9% ± 7 to 92.3% ± 26. N=3 replicates, (mean ± s.e.m).

Internalized cargo molecules are transported to early endosomes where they are either sorted to recycling endosomes and returned to the plasma membrane, or to late endosomes/lysosomes where they are degraded. For LNP to be effective and non-toxic, they should carry mRNA to compartments where it can be released into the cytoplasm or at least degraded without interfering with endosomal function. Therefore, the average residence time of LNP-mRNA in compartments that are favourable for escape must be rate-limiting for delivery (Gilleron, Querbes et al. 2013). Such residence time is proportional to the accumulation of LNP-mRNA. To gain insights into these properties, we measured the distribution of endocytosed mRNA in endosomal compartments immuno-stained against EEA1 and APPL1 to label different types of early endosomes (Kalaidzidis, Miaczynska et al. 2015), Rab11 for early/recycling endosomes (Sönnichsen, De Renzis et al. 2000) and LAMP1 for late endosomes/lysosomes (Fig. 1c-f). Interestingly, the proportion of LNP-mRNA differed quantitatively between compartments. First, LNP-mRNA distribution in EEA1-and APPL1-positive early endosomes is significantly higher than other compartments (Fig. 1g). Second, the fraction of LNP-mRNA in early (EEA1, APPL1) and recycling (Rab11) endosomes is higher for the more efficient LNP (L608, MC3 and ACU5) compared to the less efficient LNP (ACU22, MOD5 and L319). This tendency was not apparent in LAMP1 endosomes (compare L608 with MC3 and ACU5), suggesting that this compartment contributes only minimally to mRNA delivery.

### Accumulation of mRNA in Rab11 endosomes is a strong predictor of delivery efficacy

In order to rank the compartment(s) with the highest probability of escape, we used a directed acyclic graph (DAG), a general mathematical tool to infer dependencies between observed variables by differential correlation (Extended Data Fig. 3, **see Methods**). We applied DAG to the sequential transport of LNP-mRNA through the compartments of the endocytic system, from internalization to mRNA escape, to infer the correlation between amount of LNP-mRNA in a compartment and escape. Out of 9 compartments tested, only EEA1, APPL1 and Rab11 compartments had positive differential correlations (Fig. 2a-b, **see Methods**). This suggests that, in the path from uptake to escape, mRNA sequentially traverses APPL1-, EEA1-and Rab11-positive compartments, with Rab11 endosomes having the highest probability for mRNA escape (Fig. 2b, **see Methods** and Extended Data Fig. 3). This sequence is consistent with previous measurements of cargo flux through the endosomal system (Jovic, Sharma et al. 2010, Kalaidzidis, Miaczynska et al. 2015). LBPA-(multi-vesicular bodies, late endosomes), LAMP1-(late endosomes and lysosomes) and LC3-positive (autophagosomes) compartments have a negative or zero differential correlation (Fig. 2a), and, thus, are unlikely to promote endosomal escape.

**Figure 3:**
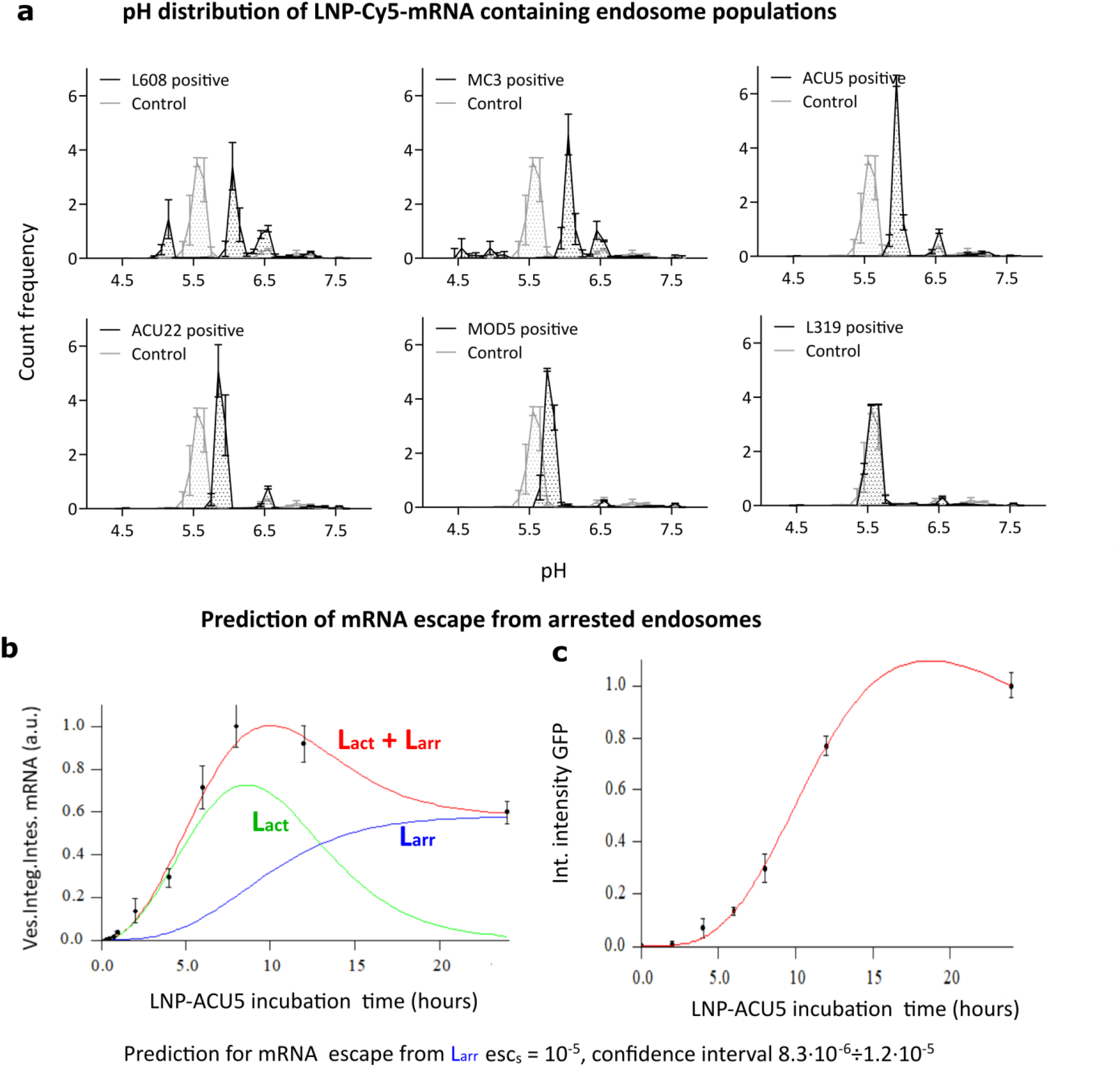
Endosomes accumulating LNP-Cy5-mRNA display acidification defects and negligible endosomal mRNA escape. **(a)** HeLa cells were incubated with pH-sensitive Rodo-Red and pH stable Alexa-488 fluorophore labeled LDLs with and without LNP-Cy5-mRNA, for 2h and imaged live. The pH of LNP-Cy5-mRNA-containing endosomes was calculated by intensity ratiometric analysis from LDL Rodo-Red and LDL-488 (see **Methods**). Whereas in control cells (LDL probes internalized without LNP-Cy5-mRNA, grey line), a significant proportion of endosomes was acidified to pH 5.5, characteristic of late endosomes/lysosomes, most LNP-Cy5-mRNA-containing endosomes (black line) stalled between early (pH 6.5) and late endosomal pH (pH 5.5). **(b-c)** Mathematical model for prediction of mRNA escape from arrested (Larr) endosomes in human primary adipocytes. The theoretical model fit (red lines in panel **b-c**) to experimental ACU5 LNP-Cy5-mRNA uptake kinetics (panel **b**, black dots) and GFP expression (panel **c**, black dots). Green and blue lines denote model predictions for active (Lact) and arrested (Larr) endosomes, respectively (see **Methods**, Extended Data Fig. 5a-b). N=3 independent experiments (mean ± s.e.m).

Rab5-and EEA1-positive early endosomes constitute a heterogeneous population that gradually increase in size as they accumulate degradative cargo until they convert into late endosomes (Rink, Ghigo et al. 2005). The small sized early endosomes actively sort recycling cargo through Rab11 endosomes whereas the large ones assume characteristics of multi-vesicular bodies (MVB) enriched in LBPA(Gruenberg 2020). Given the low correlation of LBPA to delivery (Fig. 2a), we hypothesized that the small EEA1 endosomes are more competent to delivery than the large ones. Therefore, we analysed EEA1 endosomes with respect to mRNA delivery dependent on size. We found that the fraction of endosomes with diameter >1.0 µm (denoted as EEA1_large_ in Fig. 2b) increased over time at the expense of the fraction with diameter <0.75 µm, (EEA1_small_) in L608, MC3, and ACU5 LNP (Fig. 2c). Note that accumulation of LNP-mRNA in large-sized endosomes is not due to saturation of the endosomal system, as only less than 5% of EEA1 endosomes contained LNP-mRNA during the initial two hours of uptake (Extended Data Fig. 4). The finding that LNP-mRNA accumulate (∼90% at 2h) in few (<5%) large early endosomes without converting to late endosomes (Rink, Ghigo et al. 2005) suggests that progression of cargo is inhibited. However, since such accumulation did not correlate with eGFP expression, we conclude that these compartments contribute minimally or not at all to delivery.

**Figure 4.**
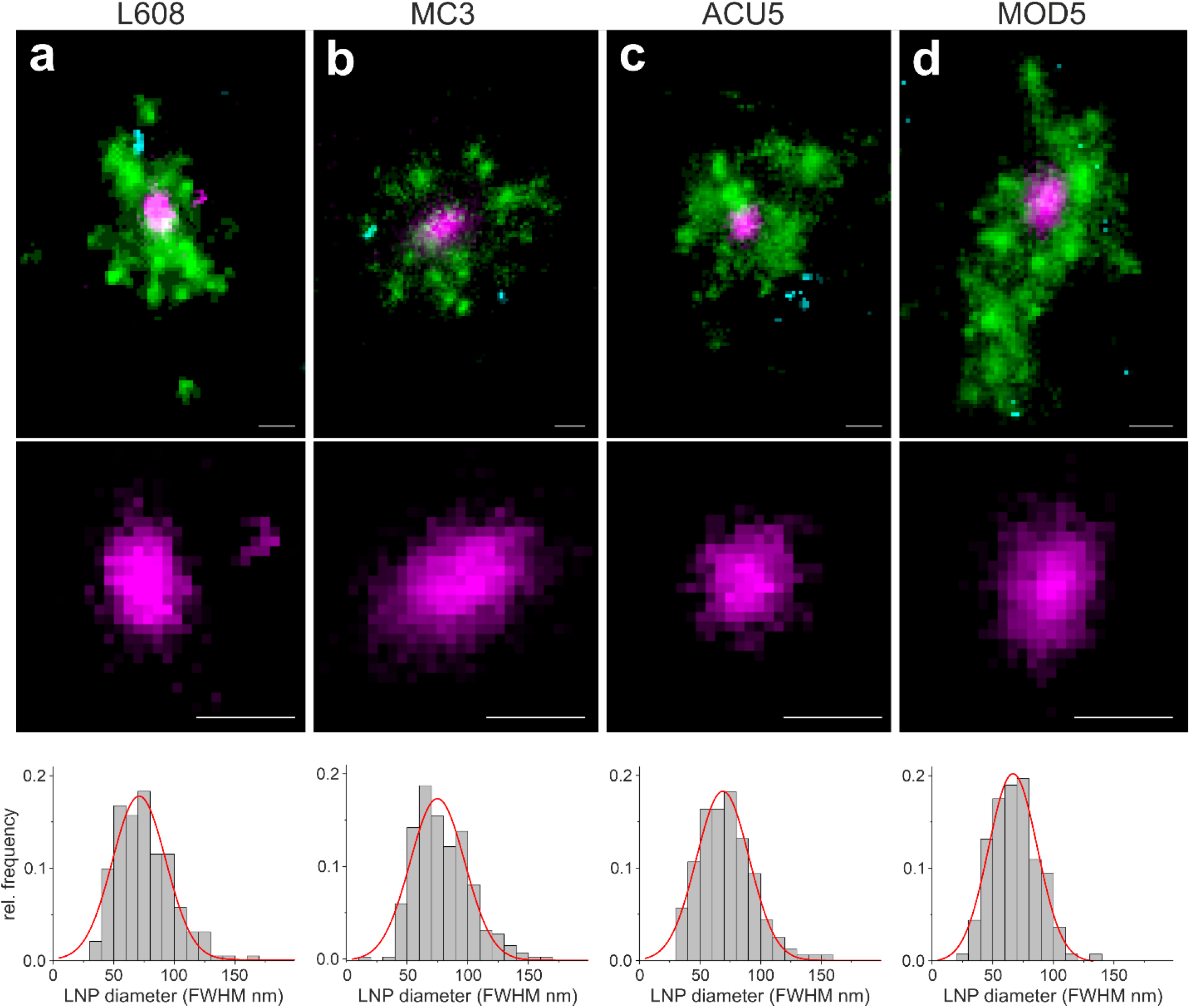
Multi-colour SMLM detects and visualizes singular LNP-mRNA in endosomal compartments with nanometer resolution. **(a-d)** Exemplary images of endosomes containing a single, clearly detectable LNP-mRNA (magenta) together with Transferrin (green) and EGF (cyan) cargo for each imaged LNP formulation (*upper row*). SMLM resolves single LNP as round nano-domains (zoom in, *middle row*) with characteristic size distribution. LNP diameter distributions were built by determining the FWHM of single LNP (see **Methods**). Mean LNP diameters (FWHM) were calculated by Gaussian fitting of the distributions to 74.9 ± 22.6, 71.0 ± 22.0, 66.7 ± 20.2 and 68.9 ± 21.7 nm, respectively (mean ± sd). The cellular context of displayed endosomes is provided in Extended Data Fig. 10, 25, 26-27. Scale bars 100 nm.

### LNP differentially impair endosomal acidification leading to mRNA accumulation in non-productive compartments

A possible mechanism blocking early endosome maturation is suppression of endosomal acidification (Novikoff, Beaufay et al. 1956, Yoshimori, Yamamoto et al. 1991, Mauthe, Orhon et al. 2018). Therefore, to measure the pH of endosomes, we used ratiometric intensity measurements (Tycko and Maxfield 1982) of LDL conjugated to pH-sensitive Rodo-Red and pH-stable Alexa Fluor-488 probes co-internalised with LNP-Cy5-mRNA for 45, 120, and 180 min and imaged in living cells (**see Methods**). The excessive amount of auto-fluorescence precluded the possibility to make such measurements in adipocytes. Therefore, we turned to HeLa cells as a well-established cell system. We first verified that HeLa cells could be transfected with LNP-mRNA (Supplementary Figure 5). In control cells that co-internalized the LDL probes without LNP-mRNA, 5% of LDL-positive structures had a pH of 6.5 and 74% a pH of 5.5, characteristic of early and late endosomes, respectively (Maxfield and Yamashiro 1987) (Fig. 3a, Supplementary Table 2). In contrast, a major fraction of LNP-Cy5-mRNA (except for L319) containing endosomes had higher pH values (between early and late endosomes), i.e. failed to acidify to late endosomal pH, corroborating the hypothesis of endosomal maturation arrest. The percentage of arrested endosomes continued to increase over time (Supplementary Table 3). Interestingly, LNP formulations differed profoundly in their effects on endosomal acidification (Fig. 3a). The poorly endocytosed L319 LNP-Cy5-mRNA did not significantly affect endosomal acidification.

**Figure 5.**
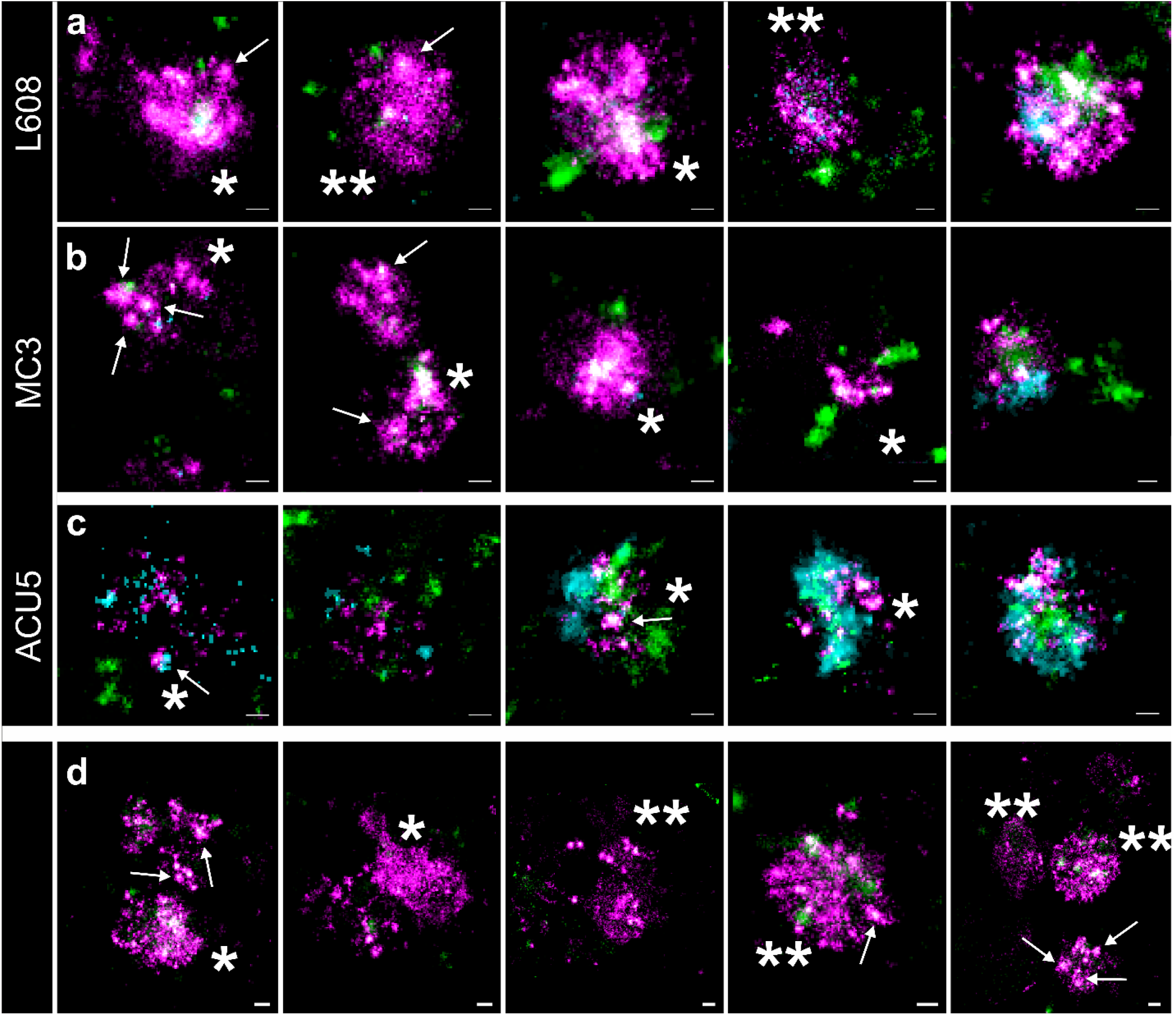
Arrested endosomes are large structures, filled with dense and disperse mRNA and often devoid of endocytosed cargo. **(a-c)** SMLM visualizes large, LNP-mRNA-rich (magenta) structures in HeLa cells, incubated with L608 (**a**) MC3 (**b**) and ACU5 (**c**) LNP. These endosomes can display an accumulation of LNP-mRNA like puncta (likely to be single LNP indicated with arrows, single asterisk), but more often a disperse signal over large areas (double asterisk). Most endosomes that exhibit these characteristics show a substantial lack of Transferrin (green) and EGF (cyan) cargo compared to endosomes, containing singular LNP (compare Figure 3), although also endosomes with varying degree of cargo-content together with strong mRNA signal can be found (**a**,**b**: increasing cargo content from left to right). (**c**) In the case of ACU5, large endosomal structures, also containing more than one LNP, can be found. Contrary to **a** and **b**, cells incubated with ACU5 often exhibit endosomes with cargo, especially EGF. (**d)** Arrested endosomes with similar features were consistently found in primary human adipocytes. ROIs, indicating the cellular context of displayed endosomes, are provided in Extended Data Fig. 10-19. Scale bars 100nm.

If endosomes are impaired in acidification, they may also fail to recycle LDL receptor to the plasma membrane, thus reducing LDL uptake. Consistent with this hypothesis, we found that the cells incubated with LNP-Cy5-mRNA that impairs endosomal acidification also show poor LDL uptake compare to the control and L319 LNP-Cy5-mRNA incubated cells (Extended Data Fig. 6-7).

**Figure 6.**
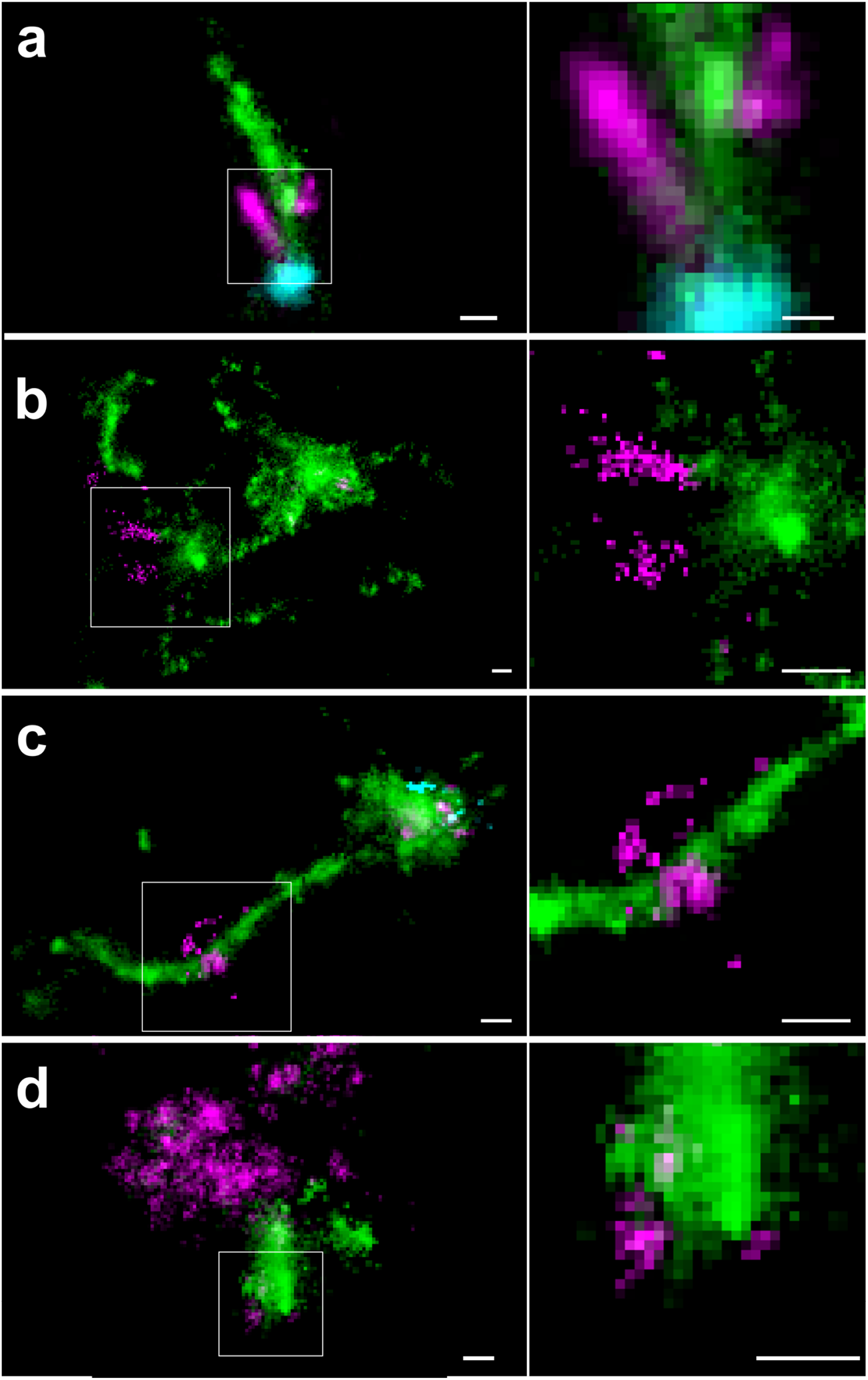
Multi-colour SMLM reveals endosomal mRNA escape at the nanoscale from Transferrin-containing tubules in primary human adipocytes and HeLa cells. Exemplary images of the different observable types of mRNA (magenta) escape in primary human adipocytes and HeLa cells, related to Transferrin positive tubules (green). **(a)** Concentrated mRNA signal is located at the very tip of a Transferrin positive tubule, connected to an elongated mRNA signal colocalising along the tubule. Very sparse mRNA signal can be detected outside the tubule. **(b)** Disperse mRNA is seemingly emanating from the Transferrin positive tubule, from which it is already segregated. These patterns clearly do not constitute intact LNP and are likely to represent partly stretched mRNA molecules escaping from endosomal structures. **(c)** An LNP is located on a long Transferrin positive tubule, together with a perpendicular disperse mRNA signal, likely representing an instance of mRNA escape. **(d)** In very rare cases, also in arrested endosomes (compare Figure 4), disperse mRNA signal, attached to a Transferrin positive structure, can be detected. ROIs, indicating the cellular context of displayed endosomes, are provided in 15, 17-26. Zoom-in of the indicated regions are presented on the right panels. Scale bars are 100 nm.

**Figure 7:**
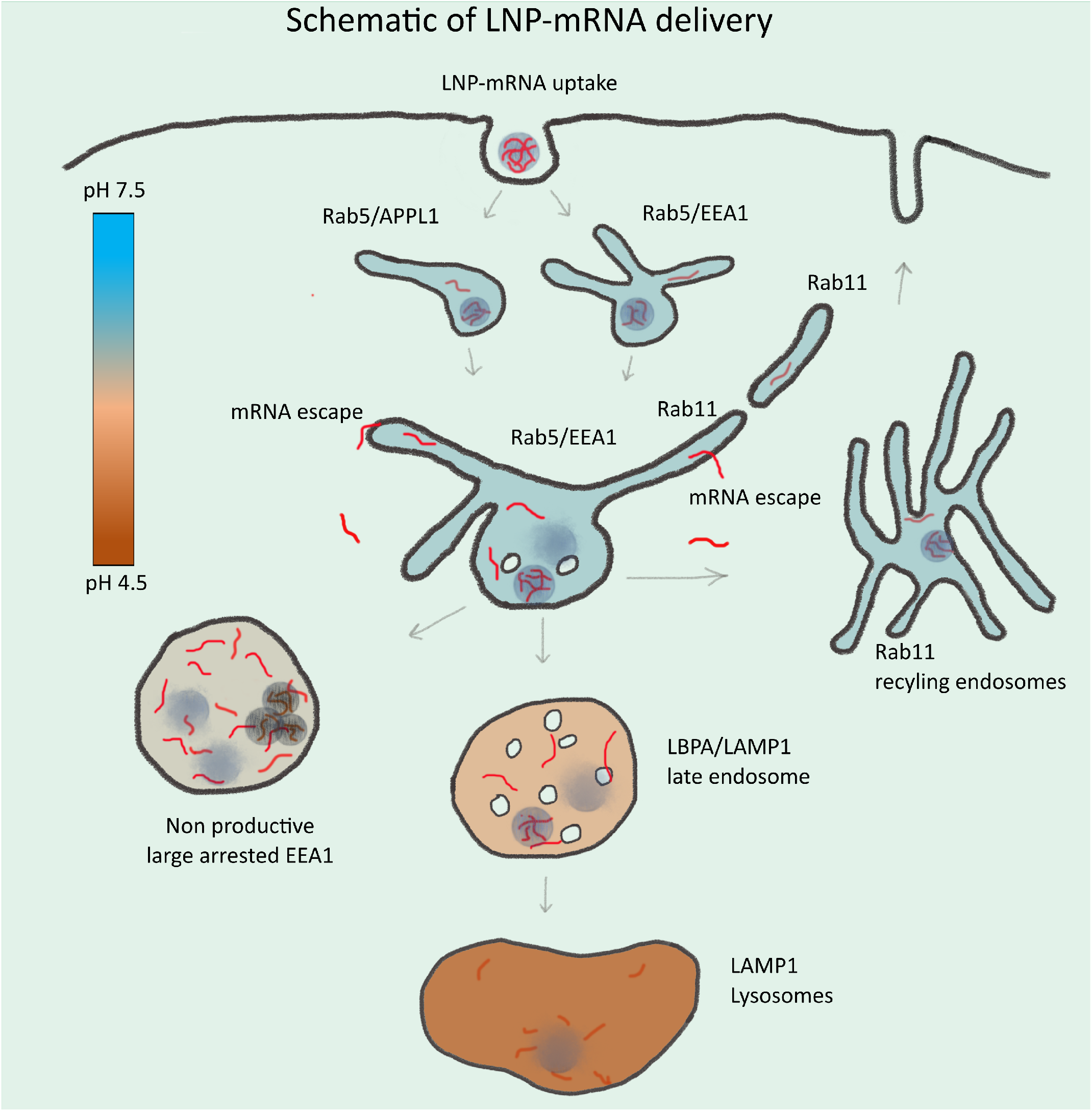
Schematic illustration of endosomal-mediated LNP-mRNA delivery: LNP-mRNA are taken up in cells via endocytosis and sequentially transported to various endosomal compartments. Under normal conditions, these endosomal compartments maintain a characteristic pH (see color heat map) for their functionality. Following uptake, mRNA in endosomes can be detected as individual LNP-mRNA similar to the starting material by SMLM. Acidification of endosomal lumen leads to release of mRNA from LNP and escape from the endosomal lumen into the cytoplasm. Escape occurs mainly from small APPL1-, EEA1-and/or Rab11-positive tubular endosomes. Over time, the majority of LNP-mRNA accumulates in large, EEA1 endosomes where individual compact LNP are disrupted and mRNA signal becomes amorphous. Large endosomes become deficient in acidification, maturation-arrested, and non-productive for delivery. This may account for cytotoxicity of LNP. Late endosomes and lysosomes are not favorable for mRNA escape.

If endosomes are impaired in maturation (Fig. 2c, Extended Data Fig. 4), acidification and cargo uptake, they are probably also unproductive for mRNA escape. A simple model and experimental data of LNP propagation through endocytic compartments provided support to this idea (Extended Data Fig. 8a, **see Methods**). In this model, we considered two groups of endosomes, “active” (L_act_) and “arrested” endosomes (L_arr_). The eGFP degradation rate was taken from Balleza Kim *et al*., (Balleza, Kim et al. 2018) and other parameters were deduced by fitting model predictions (Fig. 3b, lines) to experimentally measured ACU5 LNP-Cy5-mRNA intensity values in endosomes (Fig. 3b, black dots) and intensity of expressed eGFP (Fig. 3c, black dots) (**see Methods**). The best fit to experimental data predicted that escape from the arrested endosomes is negligible (Extended Data Fig. 8b; esc_S_ = 10^−5^, 95% confidence interval 8.3·10^− 6^÷1.2·10^−5^), indeed suggesting that they do not contribute significantly to mRNA delivery.

### Multi-colour single-molecule localization microscopy resolves singular LNP in endosomes with nanometer resolution

Our data point towards Rab11-, APPL1-and (non-arrested) EEA1-positive endosomes as potential sites for mRNA escape into the cytoplasm. All these compartments share tubular structures transporting recycling cargo, such as Transferrin, to the plasma membrane (Ullrich, Reinsch et al. 1996, Sönnichsen, De Renzis et al. 2000, Kalaidzidis, Miaczynska et al. 2015, Franke, Repnik et al. 2019). We wondered whether recycling tubules may be preferential sites of mRNA escape into the cytosol. Conventional light microscopy does not provide sufficient resolution to visualize mRNAs within endosomes or in the cytoplasm. Therefore, we deployed multi-colour single-molecule localization microscopy (SMLM) (Franke, Repnik et al. 2019). The number of mRNA molecules per LNP range between 5 and 25 (Sabnis, Kumarasinghe et al. 2018, Yanez Arteta, Kjellman et al. 2018) and each mRNA contains multiple Cy5 (average ∼25 per mRNA, Trilink, L-7701), permitting to resolve the geometry of single LNP and even single Cy5-mRNA, i.e. stretched (max. length of 996 nucleotides ≈ 300 nm, Trilink L-7701) vs. condensed. We first benchmarked our SMLM approach by visualizing isolated LNP rested on glass surface at the nanoscale (**see Methods**). Individual LNP were resolved as isolated clusters of single molecule spots with median diameter ∼60 nm (Extended Data Fig. 9). Smaller elongated clusters (ca. 25nm diameter) may correspond to damaged LNP or even dissociated Cy5-mRNAs (Extended Data Fig. 9).

Next, we performed triple-colour SMLM on cells having endocytosed both LNP and fluorescently-labelled cargo, Transferrin (Alexa Fluor 568) and EGF (Alexa Fluor 488) to label early-recycling and early-late endosomes, respectively (Franke, Repnik et al. 2019). In adipocytes, the large lipid droplets function as micro-lenses which distort the single molecule emission patterns. Therefore, we first performed SMLM in HeLa cells on the entire cell and then validated the observations in human primary adipocytes, by focusing on the endosomes underneath the plasma membrane by full TIRF. As we aimed at resolving mRNA localization within endosomes vs. cytoplasm, we refrained from using immunofluorescence labelling, which requires membrane permeabilization and may cause membrane leakage artefacts. We compared four LNP formulations with high, middle and low (L608, MC3, ACU5 and MOD5) eGFP expression. Strikingly, we could resolve individual LNP within endosomes labelled by EGF and/or Transferrin both in HeLa cells and primary human adipocytes (Fig. 4). Interestingly, singular LNP were most often associated with Transferrin-positive structures, i.e. within early-recycling compartments. To our knowledge, this is the first time that LNP are resolved by SMLM in sub-endosomal compartments. Based on our previous correlative ultrastructural analysis of early endosomes(Franke, Repnik et al. 2019), closely located clusters of single-molecule signals from labelled cargo belong to a single endosome (**see Methods**). Therefore, we quantified the size distribution of single LNP within endosomes. The size was determined as a full width at half maxima (FWHM) of intensity (**see Methods**). The calculated LNP diameters (Fig. 4) are in good agreement with measurements of LNP size by dynamic light scattering (Supplementary Table 1) and SMLM on glass surface (Extended Data Fig. 9). Therefore, SMLM allows resolving individual LNP with sub-endosomal precision.

### LNP accumulate in arrested endosomes, often lacking internalized EGF and Transferrin

For L608 and MC3 we consistently found a significant number of larger structures with an abundance of Cy5-mRNA signal in HeLa cells, primary human adipocytes as well as fibroblasts in the same culture (Fig. 5, Extended Data Fig. 10-19, 29). Within these structures, in some cases individual LNP could be resolved (arrows in Fig. 5). We also detected large Cy5-mRNA-positive structures which either correspond to LNP too closely located to be resolved (* in Fig. 5) or to dispersed Cy5-mRNA (** in Fig. 5). This suggests that some LNP are disassembled and the mRNAs released into the lumen of the endosomes. Interestingly, most of these large endosomes filled with LNP-mRNA exhibited little EGF/Transferrin signal, in contrast to endosomes having no LNP-mRNA, or small endosomes with a single LNP-mRNA which contained internalized EGF/Transferrin (Fig. 5, Extended Data Fig. 10-19). These observations corroborate the pH measurement data of Fig. 3a showing that a significant fraction of endosomal structures with accumulated LNP are in a maturation-arrested state, and insulated from normal cargo uptake (Extended Data Fig. 6-7).

### Multi-colour SMLM reveals mRNA escape from Transferrin-containing endosomal tubules

In many cases, we found Cy5-mRNA distinctly localized to Transferrin-positive tubules for all four LNP formulations (Fig. 6, Extended Data Fig. 15, 17-26). In the majority of such cases, the Cy5-mRNA signal could be observed alongside or at the very tip of tubules (Fig. 6a). Interestingly, we could also detect Cy5-mRNA signal which was not as intense and condensed as the single LNP shown in Fig. 4, but formed a pattern ranging from clustered to dispersed single molecule flashes (Fig. 6b). Such a signal was adjacent to, or outside of, Transferrin-positive structures, suggesting that this may correspond to mRNA escaped or in the process of escaping (Fig. 6b). Strikingly, in very rare cases, we could resolve Cy5 flashes ordered along a smooth line spanning a tubule and projecting into the cytoplasm (Fig. 6c). We interpret these events as capturing the escape of the mRNA from the endosomal tubule. This interpretation is supported by several lines of evidence. First, most mRNA signal has the nanoscopic appearance of single LNP and is associated with endosomal structures easily recognizable by the presence of EGF/Transferrin. Cy5-mRNA which does not colocalise with, or is in close proximity of, endosomal markers is most likely escaped into the cytoplasm. Second, as a control for false-positive Cy5-mRNA, only single LNP were visible predominantly in endosomes at the cell periphery after 30 min uptake of both LNP and EGF/Transferrin (Extended Data Fig. 28), but neither endosomal dispersed nor cytoplasmic mRNA signal was detectable. The combination of the HiLo illumination and the restriction of the fitted FWHM results in a reasonably narrow axial distribution of mRNA localizations in our SMLM images (Extended Data Fig. 30). Therefore, a random axial colocalisation of distinct Cy5-mRNA signal to Transferrin tubules can be excluded with great certainty. Third, the frequency of signal outside of endosomes was 6.9% ± 1.5 (L608), 5.9% ± 1.3 (MC3), 4.0% ± 0.7 (MOD5) and 3.0% ± 0.9 (ACU5) (mean ± SD), consistent with a very low probability of endosomal escape (**see Methods**). Fourth, the observed patterns did not change during the time of acquisition, ruling out sample drift or diffusion artefacts (Extended Data Fig. 31). Fifth, escape events could also be detected at endosomes containing multiple LNP and Transferrin, i.e. competent for cargo transport. The Cy5 signal resembling stretched mRNA molecules, was exclusively associated with the Transferrin tubules and nowhere else at the object perimeter (Fig. 6d). Sixth, we consistently found similar events in primary human adipocytes and HeLa cells (Fig. 6, Extended Data Fig. 15, 17-26), substantiating the notion that mRNA escape from Transferrin-positive tubules is a general mechanism contributing to effective transfection and shared by different cell types.

Altogether, SMLM allows distinguishing intracellular Cy5-mRNA into three categories, condensed LNP, dispersed within endosomes and a small fraction of non-endosomal de-condensed signal which is consistent with escaped cytoplasmic mRNA.

## Discussion

The biggest challenge for the delivery of mRNA, and macromolecular therapeutics in general, is to target them to the correct cells and, once endocytosed, let them cross the endosomal membrane. Only a small fraction of exogenous macromolecules can escape from endosomes via yet unknown mechanisms. To gain insights into this outstanding problem, we performed a comparative analysis of six LNP with distinct chemical composition and delivery efficiency in primary human adipocytes, fibroblasts and HeLa cells, to identify endosomal compartments which are most favourable to mRNA escape. Contrary to what is generally assumed (Novakowski, Jiang et al. 2019, Pei and Buyanova 2019), our analysis revealed that delivery efficacy cannot be predicted by total cellular uptake alone. LNP had different uptake efficiencies and, most importantly, different endosomal distributions. We found that a sub-population of EEA1-, APPL1 and Rab11-positive recycling endosomes have a higher probability for mRNA escape than the total population of Rab5 endosomes (Kalaidzidis, Miaczynska et al. 2015) and late endosomes (i.e. LBPA and LAMP1). Therefore, our data contrast both claims that RNA escape does not occur from EEA1 endosomes but from Rab5 endosomes (Wittrup, Ai et al. 2015) and that LNP recycling is counterproductive for delivery (Sahay, Querbes et al. 2013).

The endosomal compartments that correlate best with delivery efficacy, EEA1, APPL1 and Rab11, share a high proportion of recycling tubules (Sönnichsen, De Renzis et al. 2000, Kalaidzidis, Miaczynska et al. 2015). Consistent with this interpretation, by SMLM we could capture rare events of single mRNA molecules in the process of escaping from Transferrin-containing tubules in primary human adipocytes and HeLa cells. What makes endocytic/recycling endosomes particularly favourable to macromolecule escape? Multiple potential mechanisms have been proposed, such as proton sponge and subsequent endosomal bursting, cationic lipid based membrane destabilization and pore formation-mediated membrane fusion (Pei and Buyanova 2019). However, none of these mechanisms have received compelling experimental support and endosome bursting appears restricted to lipoplexes but not to LNP (Gilleron, Querbes et al. 2013, Wittrup, Ai et al. 2015). It is possible that multiple mechanisms are in play depending on differences in LNP composition and surface properties, macromolecules and their transport. Our data argue for a new mechanism. Endosomal recycling tubules are characterized by high positive curvature along the tubules and sharp transition to negative curvature at the neck of the tubules. Exogenous cationic lipids may interfere with the packing of lipids in the membrane bilayer, resulting in local instability and, thus, membrane leakage. In addition, recycling tubule fission could create spontaneous breakage of the membrane favouring macromolecular escape. Therefore, we propose that endosomes with recycling tubules are hotspots for mRNA escape events (Fig. 7).

For some LNP-mRNA formulations, a large fraction accumulates in a small population of early endosomes which are defective in new cargo uptake and arrested in maturation. We found these structures consistently in adipocytes, fibroblasts and HeLa cells, suggesting that this is a common defect. By SMLM, we could resolve mRNA packed in single LNP and in an unpacked form within these endosomes. The unpacking probably depends on lipid re-organization at low pH that facilitates release of mRNA from LNP into the endosomal lumen. Despite the abundance of LNP-mRNA in the enlarged endosomes, we never detected mRNA escape events associated with them. We interpret these endosomes as incompetent for mRNA delivery, in agreement with the correlation analysis. In view of these considerations, the claim that enhanced retention of LNP can result in productive RNA escape (Sahay, Querbes et al. 2013) is untenable. The accumulation of undeliverable LNP-mRNA in endosomes does not support the hypothesis of endosome disruption by the proton sponge mechanism and argues that retarding cargo degradation along the endosomal pathway may not increase the chance for cargo escape but rather contribute to toxic effects. An important finding was that such LNP accumulation is accompanied by endosome acidification defects. Interestingly, MOD5 (analogue of the lipid used in “mRNA-1273” SARS-CoV-2 Vaccine) also had an impact on acidification, albeit less than MC3. One potential reason could be protonation of ionisable cationic lipids in the formulation that may produce buffering effects similar to the proton sponge mechanism (Pei and Buyanova 2019). Such defect cannot persist over time if the V-ATPase maintaining the pH of endosomes remains functional. However, ionisable LNP lipids may interfere with the V-ATPase activity (Jones, Harrison et al. 1995) or increase the leakiness of endosomal membrane to protons. If the endosomal pH is above the pH required for LNP lipid reorganization, the unpacking of LNP and release of mRNA within the endosomal lumen will fail. In addition, acidification is critical to various endosomal activities, such as protein sorting, endosomal progression, lysosomal degradation and cellular homeostasis, that if compromised will have a series of cytotoxic consequences. As endo-lysosomal hydrolytic enzymes require acidic pH (Dubland and Francis 2015), suppressed acidification will prevent the biodegradation of the LNP lipids (per se bio-degradable), thus exacerbating LNP accumulation further. LNP uptake is typically mediated by LDL receptor (LDLR) (Akinc, Querbes et al. 2010). Lack of acidification will impede LDL-LDLR dissociation preventing LDLR from recycling to the surface (Davis, Goldstein et al. 1987, Bai, Zhang et al. 2018), thus blocking further uptake. This is consistent with the decreased LNP uptake at later times of internalization (Fig. 1b). Endosomal maturation arrest and accumulation of undegraded cargo are reminiscent of lysosomal storage disorders (Ballabio and Gieselmann 2009, Platt, Boland et al. 2012). In addition to cytotoxicity, these alterations may cause an inflammatory response, similar to the immune system defects characterizing lysosomal storage disorders (Castaneda, Lim et al. 2008). Therefore, defective endosomal acidification may account for a great deal of the cytotoxic effects of LNP (Garber 2017, Kulkarni, Cullis et al. 2018).

Our results define quantitative endosomal parameters that can guide the development of new mRNA formulations towards high efficacy and low cytotoxicity, by breaking down the process of delivery into at least three distinct stages. The first step to optimize is clearly the uptake of LNP, which is mediated by ApoE (Akinc, Querbes et al. 2010) and, therefore, depends on the binding of ApoE to LNP. The second stage is the dissociation of LNP and mRNA within the early endosomes, a process that can be monitored at the nanoscale (Fig. 4, 5). A parameter to monitor is the potential source of cytotoxicity by suppression of endosome acidification. The third stage is the escape of mRNA, which depends on the fraction that accumulates in recycling tubules. A possible strategy would be to develop LNP that can distribute more evenly between endosomes or even preferentially sorted to recycling tubules. For this, extended binding to LDLR may prolong the resident time in recycling endosomes, increasing efficacy and decreasing toxicity. We suggest the importance to carry out a structure/function relationship analysis of different LNP components, e.g. structure of lipid heads and tails, with respect to the different delivery stages independently of each other, but also with respect to the cytotoxic effects derived from the block of acidification. The assays and parameters to measure them described in this study can therefore complement the chemical optimization of delivery systems.

## Methods

### Cell culture

Human adipose stem cells (hASCs) were obtained from patients undergoing elective surgery at Sahlgrenska University Hospital in Gothenburg, Sweden, all of which received written and oral information before giving written informed consent for the use of the tissue. The studies were approved by The Regional Ethical Review Board in Gothenburg, Sweden. All procedures performed in studies involving human participants were in accordance with the ethical standards of the institutional and national research committee and with the 1964 Helsinki declaration and its later amendments or comparable ethical standards. All subjects complied with ethical regulations. hASCs were tested negative for mycoplasma. We adapted previously published protocol (Bartesaghi, Hallen et al. 2015) from AstraZeneca (AZ) to differentiate hASCs to mature white-like adipocytes in 384 well format. Briefly, EGM-2 proliferation medium was prepared according to the manufacture’s protocol with EBM-2 medium supplemented with 5% FBS, all provided supplements, except hydrocortisone and GA-1000 (Lonza, Cat No. 3202, EGMTM-2 MV BulletKitTM (CC-3156& CC-41472). Cryopreserved human adipose stem cells were then resuspended in EGM-2 medium and centrifuged at 200xg for 5min. Cells were counted with a CASY cell counter (Schärfe System) and 4000 cells per well were seeded in 50 µl EGM-2 medium containing 50 U/ml penicillin and 50μg/ml streptomycin (P/S) (Gibco, 15140-122) into 384 well plates (Greiner Bio-One, 781092) using the drop dispenser Multidrop. The cells were cultured at 37^°^C and 5% CO_2_ for 3 to 4 days. For adipocyte differentiation, 90% confluent cells were incubated for 1 week with Basal Medium (Zenbio, BM-1) supplemented with 3% FBS Superior, 1μM dexamethasone (Sigma Aldrich), 500μM 3-isobutyl-1-methyxanthine (Sigma Aldrich), 1μM pioglitazone (provided by AZ), P/S and 100 nM insulin (Actrapid Novonordisk, provided by AZ). Medium was replaced with BM-1 medium supplemented with 3% FBS Superior, 1μM dexamethasone, P/S and 100nM insulin and cells were incubated for another 5 days.

HeLa cells were cultured in DMEM media supplemented with 10% FBS Superior (Merck, S0615) and 50µg/mL Gentamycin (Gibco, G1397) at 37°C with 5% CO_2_. For LNP trafection for eGFP expression and endosomal pH measurement studies, 3000 HeLa cells/well were seeded in 384 well plates using the drop dispenser (Multidrop, Thermo Fischer Scientific) one day prior to experiment.

### Chemicals and reagents

FBS Superior (Merck, S0615), dexamethasone (Sigma Aldrich, D2915), 3-isobutyl-1-methyxanthine (Sigma Aldrich, I5879). Insulin (Actrapid Novonordisk) and Pioglitazone (AZ10080838) are provided by Astra Zeneca. Penicillin and Streptomycin (Gibco, 15140-122), Transferrin Alexa-568 (Invitrogen™, T23365), EGF Alexa-488 (Invitrogen™, E13345), LDL-pHRodo (Invitrogen ™, L34356), LDL-Alexa 488 was homemade, as previously reported (Franke, Repnik et al. 2019). pH calibration buffers (Invitrogen™, P35379). LNP were prepared using the cationic ionizable lipids, O-(Z,Z,Z,Z-heptatriaconta-6,9,26,29-tetraem-19-yl)-4-(N,N-dimethylamino)butanoate (MC3), L608, ACU5, ACU22, MOD5 and L319 (AstraZeneca), and the helper lipids cholesterol (Sigma Aldrich), 1,2-distearoyl-sn-glycero-3-phosphocholine (DSPC, CordenPharma), 1,2-dimyristoyl-sn-glycero-3-phosphoethanolamine-N [methoxy (polyethyleneglycol)-2000] (DMPE-PEG2000, NOF Corporation) and contained CleanCap® Enhanced Green Fluorescent Protein (eGFP) mRNA (5-methoxyuridine)and/or Clean Cap®Cyanine5 (Cy5) Enhanced Green Fluorescent Protein mRNA (5-methoxyuridine) (TriLink Biotechnologies). Formaldehyde (Merck), Digitonin (Sigma Aldrich).

### LNP mRNA formulation and characterization

All LNP were formulated by a bottom-up approach(Zhigaltsev, Belliveau et al. 2012) using a NanoAssemblr microfluidic apparatus (Precision NanoSystems Inc.). Lipids were characterized by NMR for quality control (Extended Data Fig. 32). Prior to mixing, the lipids were dissolved in ethanol and mixed in the appropriate molar ratios (cationic ionizable lipid:DSPC:cholesterol:DMPE-PEG ratios of 50:10:38.5:1.5) while mRNA was diluted in RNase free 50 mM citrate buffer pH 3.0 (Teknova). The aqueous and ethanol solutions were mixed in a 3:1 volume ratio at a mixing rate of 12 mL/min to obtain LNP with a mRNA:lipid weight ratio of 10:1. Finally, they were dialyzed overnight using Slide-A-Lyzer G2 dialysis cassettes with a molecular weight cutoff of 10 K (Thermo Scientific). The size was determined by DLS measurements using a Zetasizer Nano ZS (Malvern Instruments Ltd.) The encapsulation and concentration of mRNA were determined using the Ribo-Green assay (Supplementary Table 1).

### LNP mRNA uptake

For both mature human white-like adipocytes and HeLa cell experiments, LNP at a final mRNA concentration of 1.25ng/µl was incubated at indicated time points in the figures. Mature human white-like adipocytes were transfected in the presence of fresh BM-1 medium supplemented with 1% human serum (Sigma, H4522) to mimic the subcutaneous tissue environment. HeLa cells were transfected in the presence of 10% FBS Superior. LNP addition on 384well plates was done either using automated liquid handling robot Fluent or manual multichannel pipettes.

### Combined immuno-fluorescence and single molecule fluorescence in situ hybridization staining

Endosomes and unlabelled mRNA formulated LNP were labelled by immuno-fluorescence (IFS) and smFISH, respectively. We used an optimized protocol compatible with quantitative retention of the delivered mRNA in the cell cytoplasm [manuscript under review]. Briefly, cells incubated with LNP-mRNA were washed with PBS, fixed with freshly prepared 7.4% formaldehyde for 2h, washed 3 times with PBS and permeabilized with 0.004% Digitonin for 2min (Note: Digitonin was first purified using manufacturer’s protocol (product number D5628). Due to batch variations of this natural product, we recommend to optimize the concentration for each new purchase). After permeabilization, cells were washed and incubated with 3% BSA-PBS blocking solution for 30 to 45min. Primary antibodies prepared in 3% BSA-PBS solution were incubated for 2h and then washed 3 times with PBS. Secondary antibodies (final concentration of 3% BSA PBS solution) were incubated for 1h and washed 3 times with PBS. After this step, cells were fixed second time with 3.7% FA for 10min to preserve antibody staining and before proceeding for smFISH to fluorescently label mRNAs. The cells were washed 3 times with PBS, the supernatant was removed manually with an 8-needle aspirator and 70% Ethanol was added for 1h. After washing the cells with PBS using the Power Washer 384 (PW384, Tecan), Wash Buffer A (40 µl/well) was added for 2-5min. The supernatant was removed and eGFP -CAL Fluor® Red 590 Dye probe, from Stellaris® was diluted 1:100 in Hybridization Buffer (12.5µl/well) and incubated with cells for 16h at 37°C. The supernatant was removed, washed 2 times by incubating 40µL Wash Buffer A for 30min at 37°C. Cells were then washed again with Wash Buffer B for 2-5min. Finally, cells were incubated with DAPI (1µg/mL) to stain the nuclei and or CMB (0.25µg/mL) to stain the cytoplasm. All solutions were prepared in nuclease free PBS and all blocking and antibody solutions were mixed with 1X final concentration of nuclease inhibitors. Most washing steps and addition of solutions were performed via automated liquid handling robotic systems. All antibodies used in this study and their dilutions are given in Supplementary Table 4. We avoided double antibody staining in order to keep the green fluorescent channel free to subtract auto-fluorescence from adipocytes by image analysis methods.

### Fluorescence imaging and quantification

All confocal imaging was performed on an automated spinning disc confocal microscope (Yokogawa CV7000) using a 60x 1.2NA objective. In all cases, at least 6 images were acquired per well and each condition was in triplicate wells.

Image analysis was performed using a custom-designed software, MotionTracking (Collinet, Stöter et al. 2010) (http://motiontracking.mpi-cbg.de). Images were first corrected for illumination, chromatic aberration and physical shift using multicolour beads. All fluorescent objects in corrected images were then segmented, their number and intensity per image mask area were calculated. Cells differentiated into fibroblasts in the same culture were excluded by mask image generated by lipid droplets of adipocytes using a Fiji script. For kinetics and co-localization correlation analysis, the auto-fluorescence objects were eliminated using green channel as a base. First, the images taken by exciting green laser was used to segment auto-fluorescence objects. mRNA objects (CAL Fluor® Red 590 Dye) co-localizing to auto-fluorescence objects were then excluded from the analysis. The quality of the quantification was verified by internal non treated control cells.

Co-localization and statistical analysis were performed using the MotionTracking software. Co-localization of mRNA to endosomal compartments was performed with a threshold of 35% and corrected for random co-localization as described previously (Kalaidzidis, Kalaidzidis et al. 2015). Image analysis was also done using Fiji (Schindelin, Arganda-Carreras et al. 2012) and CellProfiler (Carpenter, Jones et al. 2006) software. Briefly, corrected images were pre-processed for segmentation in Fiji, LNP spots and nuclei were segmented and quantified in CellProfiler. Some data from MotionTracking were loaded in KNIME (Berthold, Cebron et al. 2008) for visualization with customized R scripts.

### Differential correlation analysis

For correlation analysis of mRNA distribution in endosomal compartments and eGFP expression, we chose a specific time point in LNP kinetics based on two criteria to exclude false predictions: 1) LNP mRNA signal in the endosomal compartments is above the noise level. We used LNP L319 (both uptake and eGFP expression) as an internal control due to its low uptake. 2) A time point prior to mRNA saturation in endosomal compartments.

The LNP-mRNA traffic through endocytic compartments could be drawn as a directed acyclic graph (DAG), where the nodes of the graph denote the compartments and directed edges (arrows) denote the direction of cargo flow. We expanded such presentation by adding a master node A (Extended Data Fig. 3a) to denote total uptake of LNP-mRNA by the endocytic system and node Ex to denote release of mRNA to cytoplasm (specifically, expression of eGFP as a proxy for mRNA release). The edges of such an extended graph represent cause-consequence dependencies between amounts of cargo in the endocytic compartments. Indeed, total uptake A is the cause of the amount of LNP-mRNA in compartment C, and the amount of LNP-mRNA in D depends on C, etc. The correlation between the amount of LNP-mRNA in any endocytic compartment and the amount of released mRNA (Ex) decreases proportionally to the stochasticity of the edges connecting these two nodes (regardless of the direction of connections). Indeed, total mRNA escape (node Ex) correlates with all compartments which receive LNP-mRNA from node A (nodes B-F). However, since the escape occurs from compartment D, the correlation between amount of LNP-mRNA in D and eGFP expression is the largest (r_6_). The correlation of the upstream compartment C to eGFP expression Ex is equal to r_C- >Ex_ = r_3_ * r_6_, where r_3_ is correlation between the amounts of LNP-mRNA in compartment C and D, and thus lower than r_6_. Therefore, we chose r_A->Ex_ of total LNP-mRNA uptake (A) to eGFP expression (Ex) as a baseline and subtracted it from the correlations of amount LNP-mRNA in each endocytic compartment to eGFP expression in order to get the differential correlation. It is obvious, that compartments on the path from uptake A to escape Ex (but not only them) have a differential correlation above zero. On the other hand, compartments with differential correlations below zero are located on side branches of the graph (e.g. E, F, and B). Therefore, by ranking compartments according to the differential correlation, such analysis allows reconstructing the DAG of causal dependencies for mRNA delivery (see Fig 2a& b).

### Uncertainty estimation for correlation

Let’s assume that we have two zero-mean experimental vectors {*a*_*i*_} and {*b*_*i*_} with per-element uncertainty estimation {δ *a*_*i*_} and {δ *b*_*i*_}. If elements *a*_*i*_ and *b*_*i*_ are means of experimental measurements, then δ *a*_*i*_ and δ *b*_*i*_ are SEMs.

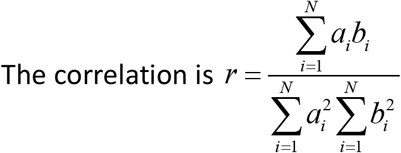

Let us assume that uncertainties of vector elements are independent and drawn from normal distribution. For example, if the elements are mean of experimental measurements, then this assumption is correct as a result of central limit theorem.

Given such assumption, the variance of correlation can be estimated as

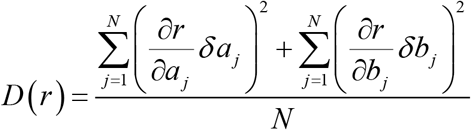

After substitution the final expression for variance of correlation is

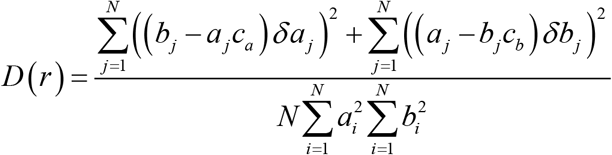

Where

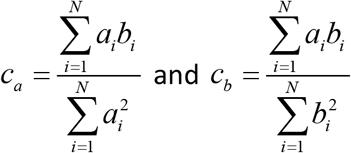

One can see, that if *r* → 1 and *a*_*i*_. *b*_*i*_ < 0, then *D (r)* → 0 independent on uncertainty δ *a*_*I*_ *and* δ*b*_*i*_.

DAG graphical schemes were prepared in Adobe Photoshop 2020 and Inkscape software.

### Ratiometric pH measurement by LDL probes

HeLa cells were seeded one day prior to transfection either in 96 or 384 well plates at a density of 12,000cells/well or 3000cells/well, respectively. pH measurements of LNP-mRNA-positive endosomes were determined by ratiometric analysis of intensities of pH sensitive LDL pHRodo-Red and pH stable LDL-Alexa 488 probes, similar to previously reported (Tycko and Maxfield 1982). Briefly, LNP-mRNA (1.25ng/µl) with or without LDL pHRodo-Red (20µg/mL) and LDL Alexa-488 (1:100 from homemade stock, **see Chemicals and reagents**) were co-internalized in cells for 45min, 2h, 3h imaged live and fixed at the end of the experiment. The pH calibration measurements were performed on each plate. LDL-pHRodo-Red/LDL-Alexa-488 were co-internalized in HeLa cells for 3h. Cells were fixed (3.7% FA, 10min), and incubated with pH = 4.5, 5.5, 6.5 and 7.5 calibration buffers (ThermoFisher, P35379) with DAPI/CMB and equilibrated for at least 2h. cells were imaged using an automated spinning disc confocal microscope (Yokogawa CV7000) 60X, 1.2NA (Extended Data Fig. 33). The intensity of pHRodo-Red increased ∼30 fold when pH decreased from 7.5 to 4.5.

Images were segmented using the MotionTracking software and individual endosomes identified. Integral intensities were calculated for both LDL probes and the ratio of integral intensities was determined. The calibration of ratios of integral intensity was performed for each individual experiment (Extended Data Fig. 34). Surprisingly, the distributions of ratios significantly varied between experiments and were multimodal for all pH values. Such wide multimodal distribution does not allow to use the mean value of ratios of calibration measurements as a coefficient to convert ratios into pH for the live cell measurements (Extended Data Fig. 35)

We hypothesized that the ratio of distributions could be described as the sum of log-normal distribution and used Bayesian analysis to find how many components are required to describe them (Sivia and Carlile 1992). From this, we found that the most probable number of components was 3. Therefore, we globally (simultaneously) fit the calibration of distributions of 3 independent experiments by the sum of log-normal components, keeping the parameters of components equal for the experiments, but allowing different amplitudes:

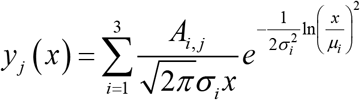

Where *y* _*j*_ *(x)* is the model distribution of ratios in the repeat *j*; *x* is ratio of intensities; μ*I*, σ *I* are parameters of component *i, A*_*i, j*_ is the contribution of component *i* in the repeat *j*. The resulting parameters μ_*i*_, σ_*i*_ as function of pH are presented on Extended Data Fig. 36. Given, that the fit was performed independently for each pH, the smooth changes of parameters values with pH circumstantially support our 3-component model.

The deduced parameters allowed us to interpolate intensities ratio distributions for intermediate values of pH (Extended Data Fig. 37).

In order to find the pH distribution in endocytic compartments, we divided the pH interval from 4.5 to 7.5 on bins by steps of 0.05 units. We introduced amplitudes of all 3 components for each bin (*A*_*i*_*(pH), i* = 1,2,3) and used them as free variables to fit theoretical distribution to experimental data (Extended Data Fig. 38a). The fit was performed by library Climb of MotionTracking software. Finally, the pH distribution was calculated as a sum of amplitudes of all 3 components in each pH bin: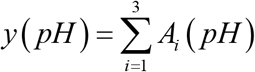 The results of such fit for 3 independent experiments are presented on (Extended Data Fig. 38b).

Despite differences in calibration distributions of experimental repeats (Extended Data Fig. 34), the predicted pH distributions of independent experiments are very much in agreement with each other (Extended Data Fig. 38b) and, as such, provide additional support to our model.

### LNP uptake and eGFP expression kinetics measurements

For determining the arrested endosomes contribution to mRNA escape, we used ACU5 LNP formulated with either unlabelled (eGFP expression) or Cy5 fluorophore labelled mRNA (uptake). Adipocytes were incubated with ACU5 mRNA formulation for 20min, 30min, 45min, 60min, 2h, 4h, 6h, 8h, 12h, 24h and fixed with optimized protocol [manuscript under review]. The cells were then stained for nuclei (DAPI 1µg/ml) and cytoplasm (CMB 0.5µg/mL), imaged using an automated confocal microscope (Yokogawa CV7000), the total eGFP intensity and the segmented mRNA intensities were analysed using the MotionTracking software.

### Model for prediction of mRNA escape from arrested endosomes

Experimental data and theoretical model were combined to determine the contribution of the maturation arrested endosomes to mRNA escape. In our model, we considered two groups of endosomes – “active” and “arrested”. The LNP propagate through “active” early endosomes and either route towards late endosomes and lysosomes for degradation or long-living “arrested” endosomes. The amount of LNP in “active” and “arrested” endosomes are respectively denoted as Le and Ls (Extended Data Fig. 8a. eq. (1) and (2)). The lag in LNP uptake is modelled by the transition function F(t). The amount of LDL receptors (LDLR) on the plasma membrane is denoted by R (eq.(1) and (3)). Surprisingly, we found that we could explain the declining LNP uptake (between 8h-24h, Figure 3b, Black dots) without assuming that a massive uptake of LNP leads to the downregulation of LDLR from the plasma membrane (eq. (3)). This led us to hypothesize that the “arrested” endosomes are not sufficiently acidic to release LNP from LDLR, thus preventing its recycling to the plasma membrane. This process was modelled by eq. (3). The fraction of mRNA that leaks from early “active” and “arrested” endosomes are described by eq. (4), where esc_s_ denotes the ratio of mRNA leakage from “arrested” to “active” endosomes. The eq. (5) describes the synthesis and degradation of eGFP (Balleza, Kim et al. 2018).

### Single molecule localization microscopy

Cover-glasses of 24mm diameter were first sonicated in 70% ethanol for 5min and washed 3 times using autoclaved water and PBS. The clean cover glasses were distributed on 6 well plate and seeded HeLa cells at a density of either 200,000 cells prior to one day of the experiment. Human primary adipose stem cells were seeded at the density of 500,000 cells and differentiated to adipocytes as mentioned in cell culture methods. The cells were incubated with LNP-mRNA (at a final mRNA concentration of 1.25ng/µl) for total of 120min. EGF-Alexa 488 (100ng/mL) and Transferrin-Alexa 568 (10µg/mL) or homemade LDL-Alexa 488 (1:100 dilutions, **see Chemicals and reagents**) was incubated for last 30min of LNP uptake. LNP incubation in HeLa cells were supplemented with 10% FBS and primary human adipocytes supplemented with 1% Human serum. The cells were then fixed with 7.4% FA for 2h, washed 3 times with PBS.

Multi-color SMLM experiments were performed on a Nikon Eclipse Ti microscope, which is specified elsewhere in detail (Franke, Repnik et al. 2019). Prior to acquisition, cells were irradiated in epifluorescence illumination mode to turn emitters, which were out-of-focus in the HILO illumination scheme, into the dark state. In all experiments, the length of the acquisition was set to capture the majority of emitters, i.e. imaging was concluded when only a very minor number of active emitters was detectable. Typical acquisition lengths were 30000-60000 frames for 641 and 561 nm and 15000-30000 frames for 488 nm excitation, respectively. Hereby, mEOS2 was excited at 561 nm and activated with 405 nm. Raw image stacks were analysed with rapidSTORM 3.2 (Wolter, Loschberger et al. 2012). The FWHM was set as a free fit parameter, but in the limits of 275–475 nm for 561 nm and 640 nm and 275-450 nm for 488 nm excitation, respectively. This way, we allow only a narrow axial range (ca. 500 nm) (Franke, Sauer et al. 2017) to contribute to the final SMLM data set, in order to minimize the likelihood of random axial colocalisation. The fit window radius to 1200 nm, while all other fit parameters were kept from the default settings in rapidSTORM 3.2. Linear lateral drift correction was applied manually by spatio-temporally aligning distinct structures to themselves. This was facilitated by color-coding of the temporal coordinate with the built-in tool. Same tool was used to create spatio-temporal images of mRNA escape events (Extended Data Fig. 31).

Prior to imaging of samples, a glass surface with Tetraspeck beads (Thermo Scientific) was imaged with alternating 488 nm, 561 nm and 641 nm excitation to create a nanometer precise map to allow the correction of chromatic shift.

Candidates for single LNP were selected in the range of 25 – 250 nm on the basis of previously reported LNP diameters. FWHMs were determined from the super-resolved images (10 nm pixelsize) by Gaussian fitting. Mean diameters for LNP in cells were determined to 74.9nm ± 22.6 (mean ± SD) for L608, 71.0nm ± 22.0 for MC3, 66.7nm ± 20.2 for ACU5 and 68.9nm ± 21.7 for MOD5.

In adipocytes, large lipid droplets function as micro-lenses, making quantitative SMLM-based studies very challenging, since the three-dimensional distortions to the single molecule emission patterns negate the theoretical advantage of nm resolution. Therefore, we imaged only the endosomes underneath the plasma membrane at the bottom of adipocytes by full TIRF.

### Estimation of cytosolic mRNA signal

LNP/mRNA localizations were classified as non-cytosolic when satisfying one of the following criteria:

- Object is colocalised with endosomal cargo
- Object is localized within an arrested endosome
- Object is localized on cellular edge (cell border)

All other mRNA localizations not fulfilling at least one of the above criteria were considered likely to be cytosolic and the ratio regarding the total number of localizations was given.

However, it is important to consider that the photo-physical properties of the multiple Cy5 molecules per mRNA may vary between the different settings, i.e. mRNA cramped in a LNP, mRNA escaped or fully elongated in the cytosol. The actual fraction might therefore be even lower, since it is to be expected, that possible quenching effects between the dyes will occur predominantly in the densely packed environment of the intact LNP, thus reducing the localization count.

## Acknowledgements

We thank the following Services and Facilities of the Max Planck Institute of Molecular Cell Biology and Genetics for their support: Light Microscopy Facility (LMF) and the High-Throughput Technology Development Studio (HT-TDS) Facility. We thank the Center for Information Services and High Performance Computing (ZIH) of the TU Dresden for the generous provision of computing power. This work was financially supported by the Max Planck Society (MPG) and AstraZeneca.

## Author contributions

M.Z., S.A., P.P., C.F conceived the project, P.P., C.F., M.S. and M.B. designed the experiments. M.Y.A formulated and characterized LNP. P.P. and M.S. performed experiments in adipocytes and HeLa cells, analysed and interpreted the results. C.F. performed single molecule localization microscopy experiments, analysed data and interpreted the results. Y.K conceived the mathematical models for pH and endosomal escape from arrested endosomes, programmed and interpreted the results. M.Z., P.P., C.F., and Y.K. wrote the manuscript with edits by M.Y.A. M.S. M.B., A.H., S.B., A.S., L.L., A.D., A.B., and S.A. commented on results and interpretations. P.P. and C.F. contributed equally to this study.

