## Supplementary Data for "Endosomal escape of delivered mRNA from endosomal recycling tubules visualized at the nanoscale"

### Extended Data Figures: Overview

Extended Data Figure 1: Structure of lipids used for different LNP formulation

Extended Data Figure 2: Representative images for eGFP expression

Extended Data Figure 3: DAG scheme for differential correlation analysis of LNP-mRNA delivery

Extended Data Figure 4: Percentage of EEA1 endosomes co-localized to LNP-mRNA

Extended Data Figure 5. eGFP expression in LNP-mRNA transfected HeLa cells

Extended Data Figure 6. LDL-488 uptake in HeLa cells

Extended data Figure 7. pH distribution LNP-containing endosomes in HeLa cells after 45min uptake

Extended Data Figure 8. Model for prediction of mRNA escape from arrested endosomes

Extended Data Figure 9: LNPs on glass surfaces visualized by SMLM

Extended Data Figure 10: Partial cellular overview of SMLM data in a HeLa cell

Extended Data Figure 11: Partial cellular overview of SMLM data in a HeLa cell

Extended Data Figure 12: Partial cellular overview of SMLM data in a HeLa cell

Extended Data Figure 13: Partial cellular overview of SMLM data in a HeLa cell

Extended Data Figure 14: Partial cellular overview of SMLM data in an adipocyte with ROIs indicating endosomes

Extended Data Figure 15: Partial cellular overview of SMLM data in an adipocyte with ROIs indicating additional examples of arrested endosomes for the L608 LNP formulation

Extended Data Figure 16: Partial cellular overview of SMLM data in an adipocyte with ROIs indicating additional examples – L608

Extended Data Figure 17: Partial cellular overview of SMLM data in an adipocyte with ROIs indicating additional examples – L608

Extended Data Figure 18: Partial cellular overview of SMLM data in an adipocyte with ROIs indicating additional examples – MC3

Extended Data Figure 19: Partial cellular overview of SMLM data in an adipocyte with ROIs indicating additional examples – MC3

Extended Data Figure 20: Partial cellular overview of SMLM data in an adipocyte with ROIs indicating additional examples – ACU5

Extended Data Figure 21: Partial cellular overview of SMLM data in an adipocyte with ROIs indicating additional examples – ACU5

Extended Data Figure 22: Partial cellular overview of SMLM data in an adipocyte with ROIs indicating additional examples – MOD5

Extended Data Figure 23: Partial cellular overview of SMLM data in an adipocyte with ROIs indicating additional examples of possible mRNA escape events –L608

Extended Data Figure 24: Partial cellular overview of SMLM data in a HeLa cell with ROIs indicating the endosome presented in Figure 6b

Extended Data Figure 25: Partial cellular overview of SMLM data in a HeLa cell with ROIs indicating the endosome presented in Figure 4c and Figure 6c

Extended Data Figure 26: Partial cellular overview of SMLM data in a HeLa cell with ROIs indicating additional examples of possible mRNA escape

Extended Data Figure 27: Partial cellular overview of SMLM data in HeLa cells with ROIs indicating the endosome presented in Figure 4d

Extended Data Figure 28: Exemplary field of views of cells incubated with LNPs and cargo molecules simultaneously for 30 minutes

Extended Data Figure 29: Arrested endosomes in primary fibroblasts

Extended Data Figure 30: Distribution of fitted FWHM of mRNA-Cy5 localizations

Extended Data Figure 31: The temporally color-coded images of mRNA escape events

Extended Data Figure 32: NMR characterization data of cationic lipids (Page 33 – 35)

Extended Data Figure 33: Representative images of co-internalized mixture of LDL-pHRodo-Red/LDL-Alexa-488

Extended Data Figure 34: Distribution ratios of integral intensities of pHRodo-Red/Alexa-488 in calibration measurements

Extended Data Figure 35: Distribution of objects ratios of integral intensities of pHRodo-Red/Alexa-488 in live HeLa cells

Extended Data Figure 36: pH dependency of parameters $\mu$ and $\sigma$ of log-normal components

Extended Data Figure 37: Predicted distribution of ratios with equal contributions components at pH in range from 4.5 to 7.5

Extended Data Figure 38: Experimental distribution of intensities ratios and Fitted distribution of pH.

Supplementary Table 1: LNPs size and encapsulation efficiency calculated by DLS and Ribo green assays

Supplementary Table 2: Percentage of LNP-Cy5 mRNA containing endosomes with indicated pH-2h

Supplementary Table 3: Percentage of LNP-Cy5 mRNA containing endosomes with indicated pH-3h

Supplementary Table 4: Details of antibodies and their dilutions used in this study

Supplementary References

**L608**

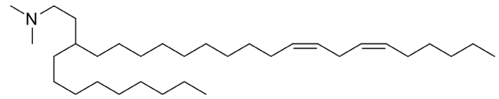

WO 2017/201350 A1, L608

**MC3**

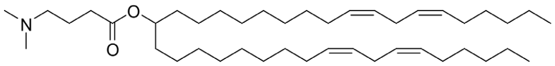

Jayaraman, Ansell et al. 2012

**ACU5**

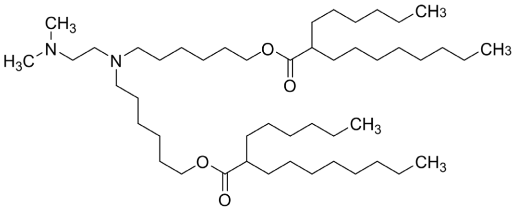

WO 2015/199952 A1, compound 5

**ACU22**

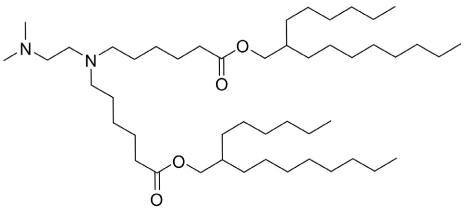

WO 2015/199952 A1, compound 22

**MOD5**

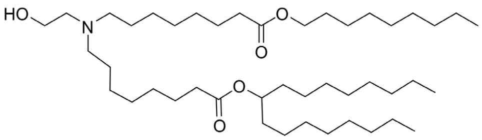

Sabnis, Kumarasinghe et al. 2018

**L319**

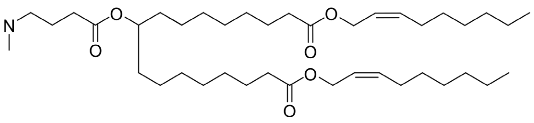

Maier, Jayaraman et al. 2013

**Extended Data Figure 1**: **(a)** Structure of cationic lipids used in different LNP-mRNA formulations.[1-5]

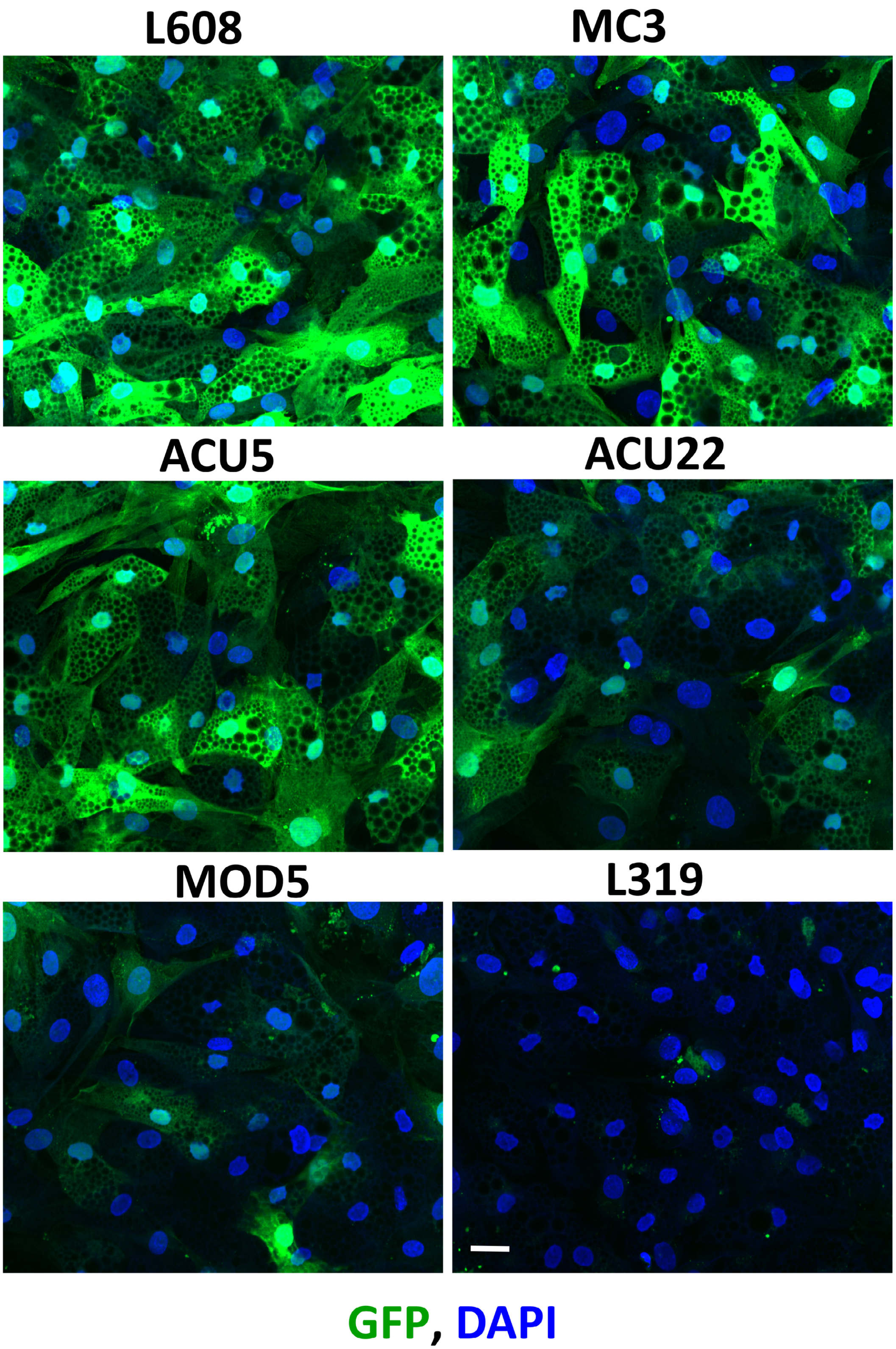


**Extended Data Figure 2**: **(a)** Representative images of human primary adipocytes expressing eGFP after 24h of LNP-mRNA uptake. The cells were fixed, stained for DAPI and imaged by fluorescence microscope. Bar indicates 20μm for all images.

**DAG analysis schematic**

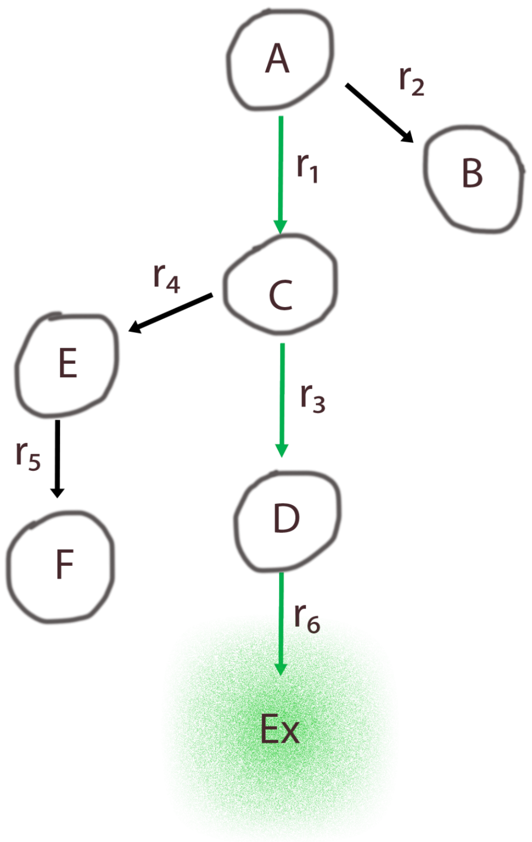


**Extended Data Figure 3:** DAG scheme for differential correlation analysis of LNP-mRNA delivery. The first (root) node A represents the total uptake of LNP-mRNA and the last node Ex represents eGFP expression as proxy for mRNA escape. All other nodes of the graph represent the amount of LNP-mRNA in different endocytic compartments received from node A. Compartments on the path from mRNA uptake to endosomal mRNA escape are represented by green edges. Compartments that contribute minimally to mRNA escape are on the side branches (black edges, E, F and B). (**see Methods**).

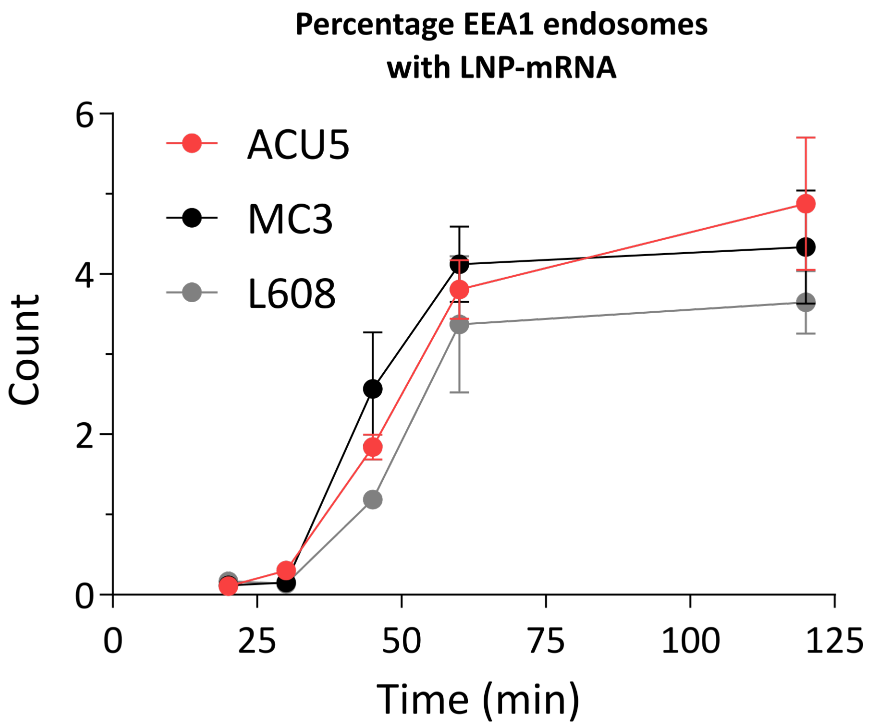


**Extended Data Figure 4.** Percentage of EEA1 endosomes co-localized to LNP-mRNA. The graph illustrates that only low percentages of EEA1 endosomes contain LNP-mRNA after 2h of uptake.

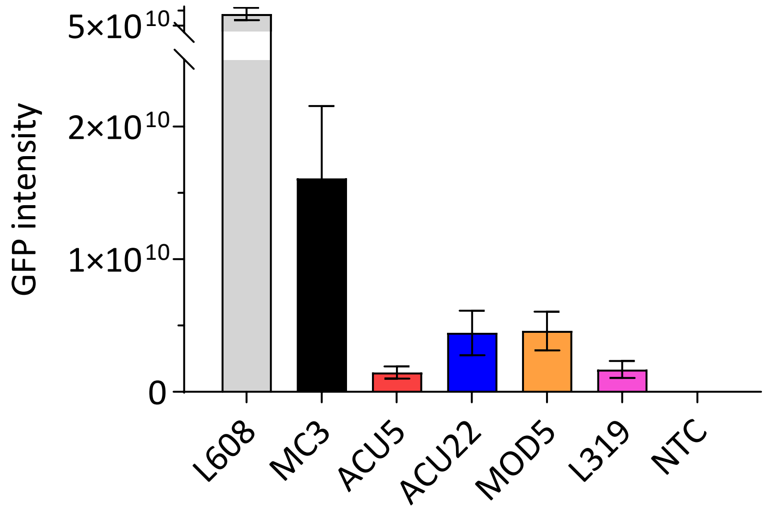


**Extended Data Figure 5.** eGFP expression in LNP-mRNA transfected HeLa cells. HeLa cells were incubated with LNP-mRNA (1.25ng/µL) for 24h, fixed and imaged. eGFP expression was quantified by MotionTracking software.

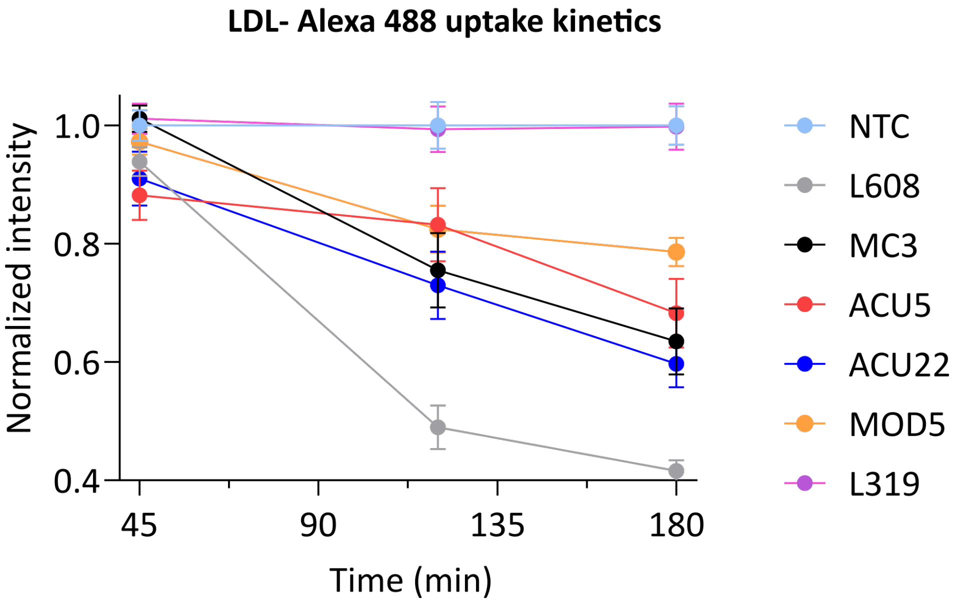


**Extended Data Figure 6. LDL-488 uptake in HeLa cells.** LDL-Alexa Fluor 488 total uptake kinetics in HeLa cells that were co-incubated with LNP-Cy5-mRNA (1.25ng/µL). The total LDL uptake was quantified by integrated intensity of LDL positive objects and normalized to non-LNP-treated control NTC (i.e LDL uptake without LNP-Cy5-mRNA). The graph shows that when cells treated with LNP-Cy5-mRNA, the uptake of LDL is mildly affected at 45min, at which time point the pH of endosomes is not severely impaired (see Extended Data Figure 7a). However, the LDL uptake was significantly reduced over time especially when cells were treated with LNP-Cy5-mRNA that blocked endosomal acidification at the given time points (Supplementary Table 2 & 3). L319 LNP-Cy5-mRNA does not block endosomal acidification and show LDL uptake similar to that of non-LNP-treated control (NTC).

**pH distribution of LNP-Cy5-mRNA containing endosome populations - 45min**

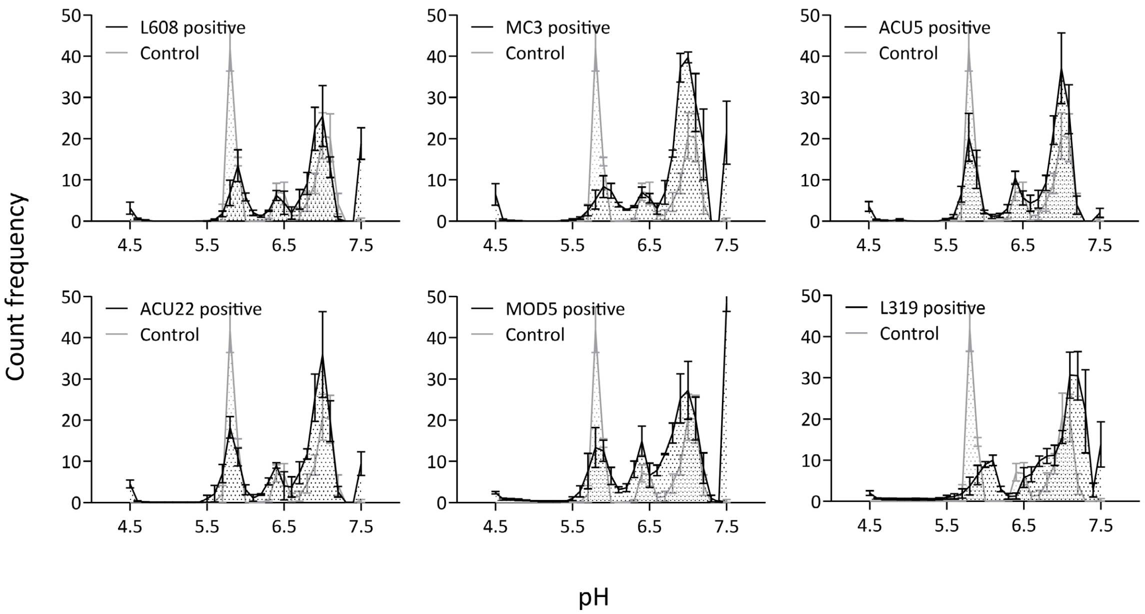


**Extended Data 7: pH distribution LNP-Cy5-mRNA containing endosomes in HeLa cells.** LDL-probes co-incubated with LNP-Cy5-mRNA (1.25ng/µL) for 45min and imaged live. The graph shows that the pH of LNP-Cy5 mRNA containing endosomes are only mildly affected in contrast to 120min and 180min time points presented in Figure 3a, Supplementary Table 2, and 3.

**a**

$$F(t) = 1 - e^{-\left(\frac{t}{t_0}\right)^2}$$

$$\begin{cases} \dfrac{dL_e}{dt} = F(t) \cdot k_{in} \cdot R - k_{\deg_0} \cdot L_e - k_s \cdot L_e & (1) \\ \dfrac{dL_s}{dt} = k_s \cdot L_e - k_{\deg} \cdot L_s & (2) \\ \dfrac{dR}{dt} = k_{ldl\_syn} - k_{ldl\_\deg} \cdot R - F(t) \cdot k_{in} \cdot R & (3) \\ \dfrac{dmGFP}{dt} = \dfrac{1}{1+esc_s} L_e + \dfrac{esc_s}{1+esc_s} \cdot L_s - k_{m\_\deg} \cdot mGFP & (4) \\ \dfrac{dGFP}{dt} = k_{syn} \cdot mGFP - k_{g\_\deg} \cdot GFP & (5) \end{cases}$$

**b**

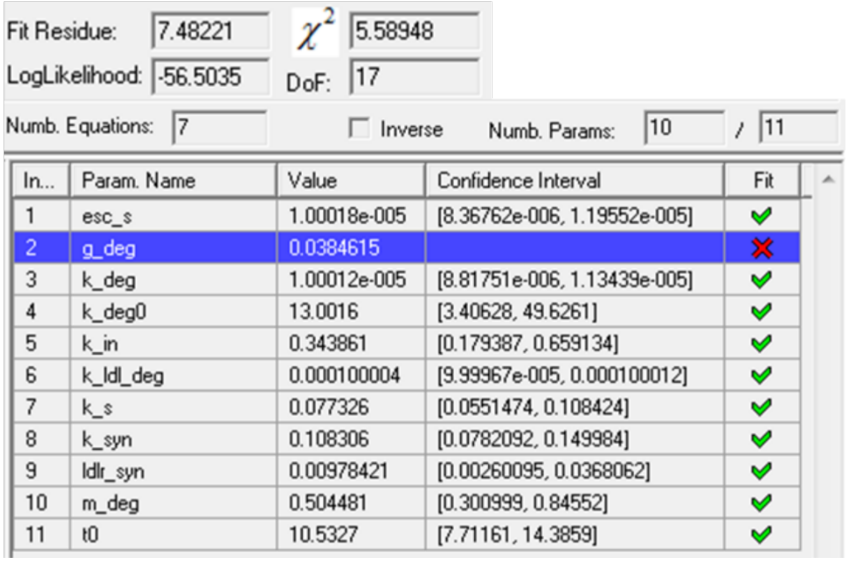

Fit Residue: 7.48221 | $\chi^2$ 5.58948
LogLikelihood: -56.5035 | DoF: 17
Numb. Equations: 7 | Inverse | Numb. Params: 10 / 11

| In... | Param. Name | Value | Confidence Interval | Fit |
|---|---|---|---|---|
| 1 | esc_s | 1.00018e-005 | [8.36762e-006, 1.19552e-005] | ✓ |
| 2 | g_deg | 0.0384615 | | ✗ |
| 3 | k_deg | 1.00012e-005 | [8.81751e-006, 1.13439e-005] | ✓ |
| 4 | k_deg0 | 13.0016 | [3.40628, 49.6261] | ✓ |
| 5 | k_in | 0.343861 | [0.179387, 0.659134] | ✓ |
| 6 | k_ldl_deg | 0.000100004 | [9.99967e-005, 0.000100012] | ✓ |
| 7 | k_s | 0.077326 | [0.0551474, 0.108424] | ✓ |
| 8 | k_syn | 0.108306 | [0.0782092, 0.149984] | ✓ |
| 9 | ldlr_syn | 0.00978421 | [0.00260095, 0.0368062] | ✓ |
| 10 | m_deg | 0.504481 | [0.300999, 0.84552] | ✓ |
| 11 | t0 | 10.5327 | [7.71161, 14.3859] | ✓ |

**Extended Data Figure 8. (a-b)** Model for prediction of mRNA escape from arrested endosomes**.** The parameters fitted with 95% confidence intervals on experimental data presented in Figure 3 b & c (black dots). The theoretical fit brings the fraction mRNA escape from arrested endosomes to zero (i.e esc_s = $10^{-5}$, confidence interval $8.3 \cdot 10^{-6} \div 1.2 \cdot 10^{-5}$). (**see Methods).**

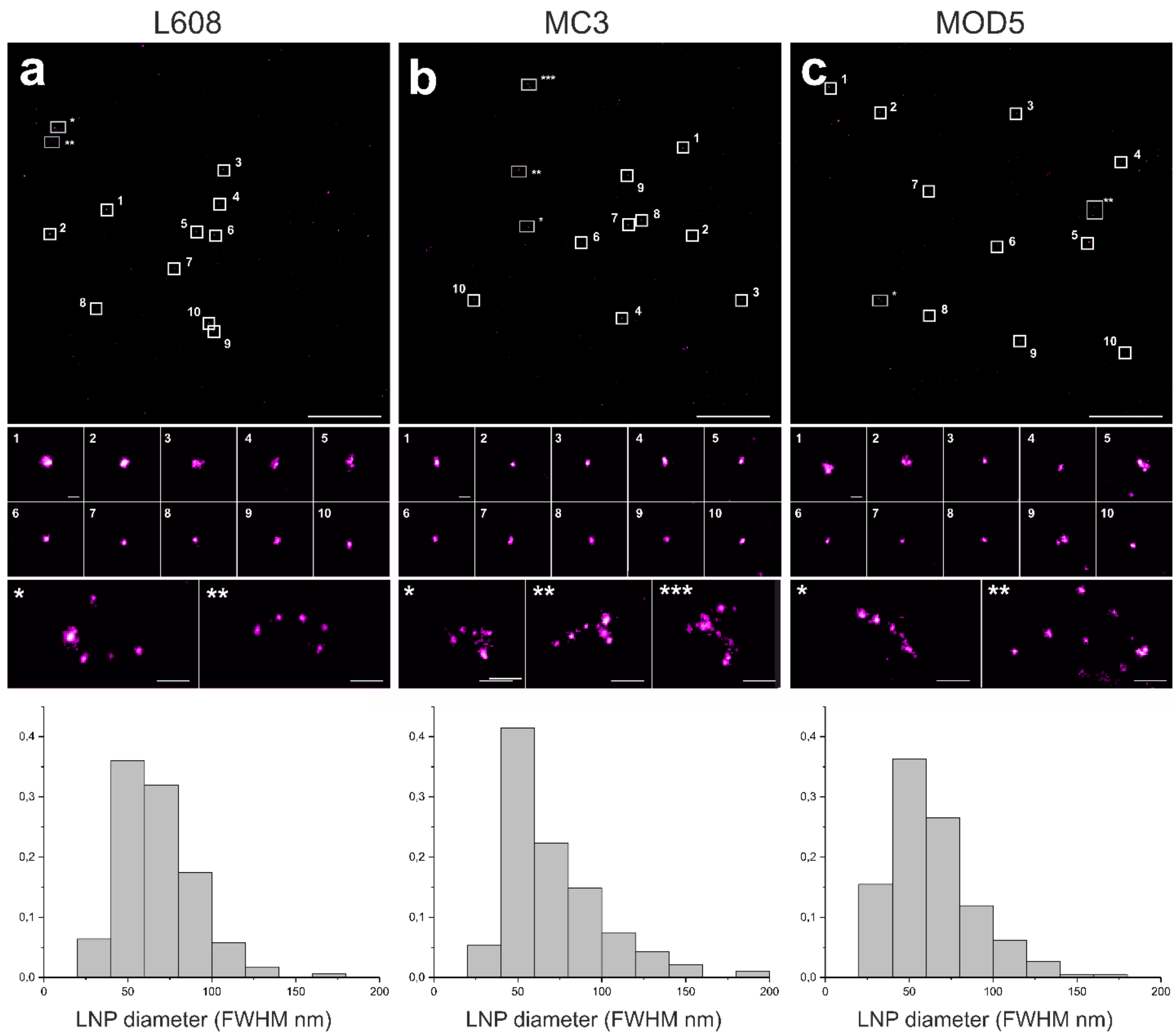


**Extended Data Figure 9**. Exemplary field of views of LNP-Cy5-mRNA (magenta) deposited on glass surfaces visualized by SMLM. All imaged LNP-mRNA formulations were detected as concentrated small spots with consistent sizes over the entire field of view. Diameters (FWHM) were determined to 67.4 ± 22.2 nm, 72.9 ± 32.3 nm and 63.9 ± 25.0 nm for LNPs L608, MC3 and MOD5 respectively (mean ± sd). Scale bars 5 µm.

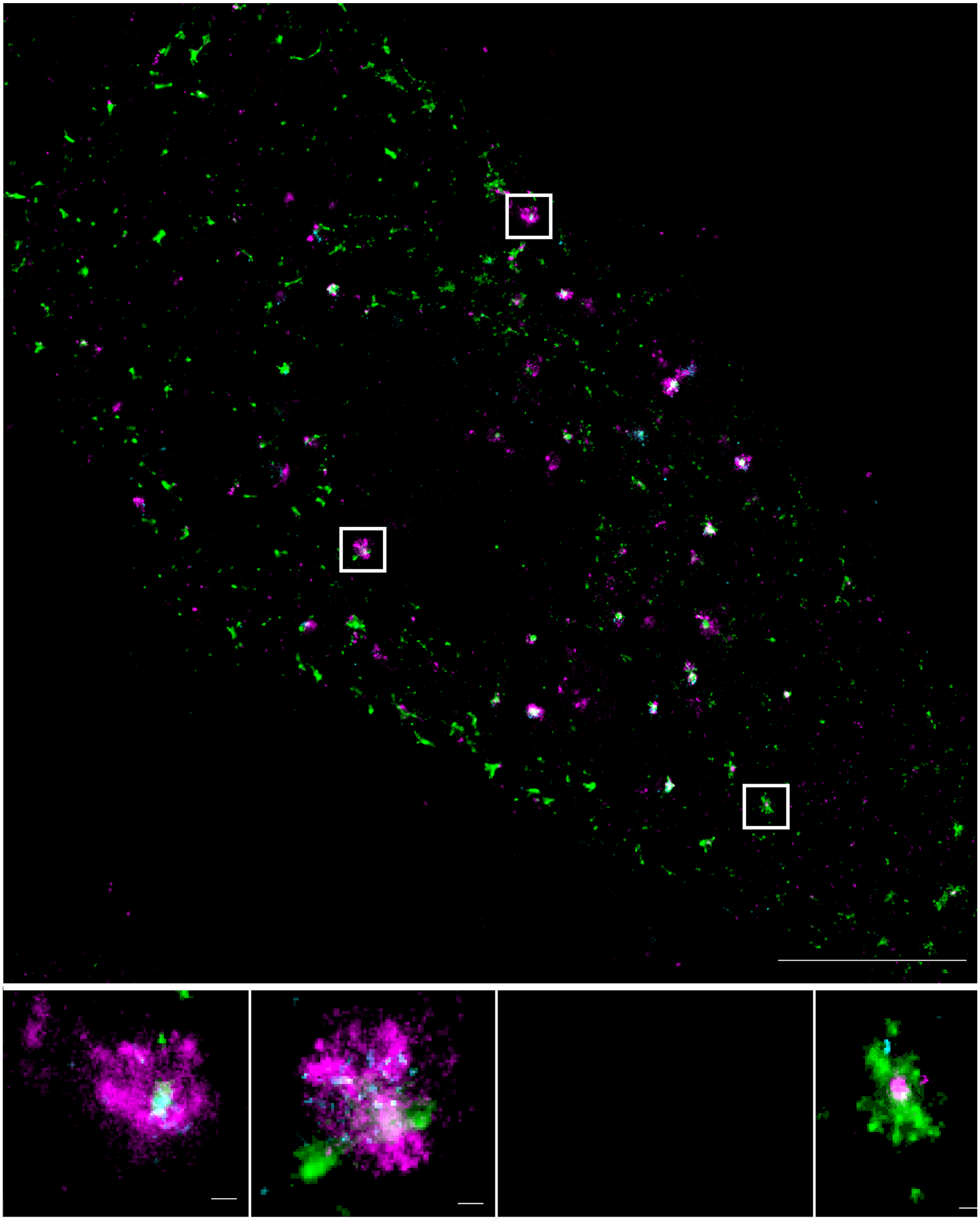

**Extended Data Figure 10.** Partial cellular overview of SMLM data in a HeLa cell with ROIs indicating endosomes presented in Figure 4a and Figure 5a. LNP-Cy5-mRNA (magenta), Transferrin (green), EGF (cyan). Scale bars 5 µm (top), 100nm (bottom).

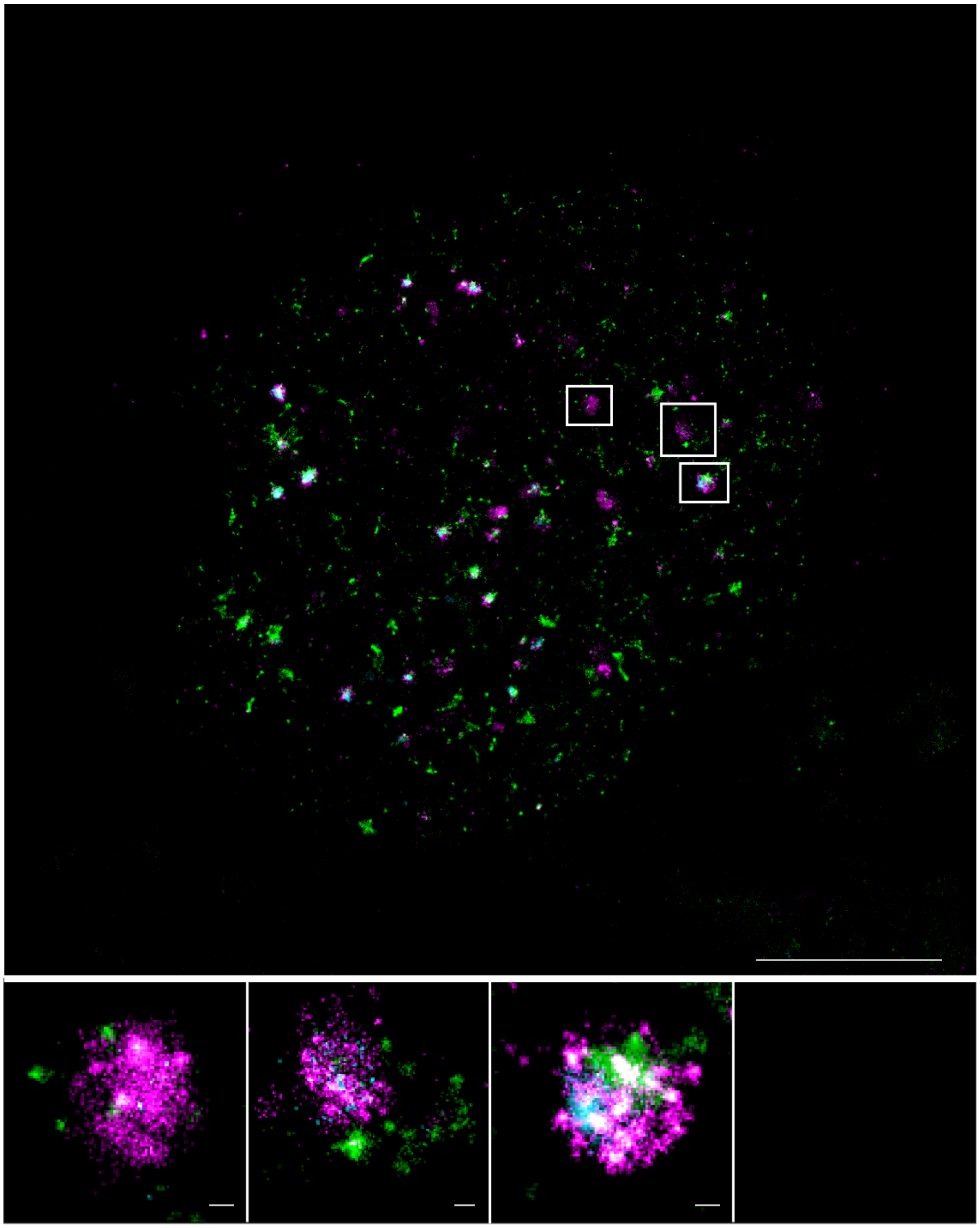

**Extended Data Figure 11.** Partial cellular overview of SMLM data in a HeLa cell with ROIs indicating endosomes presented in Figure 5a. LNP-Cy5-mRNA (magenta), Transferrin (green), EGF (cyan). Scale bars 5 µm (top), 100nm (bottom).

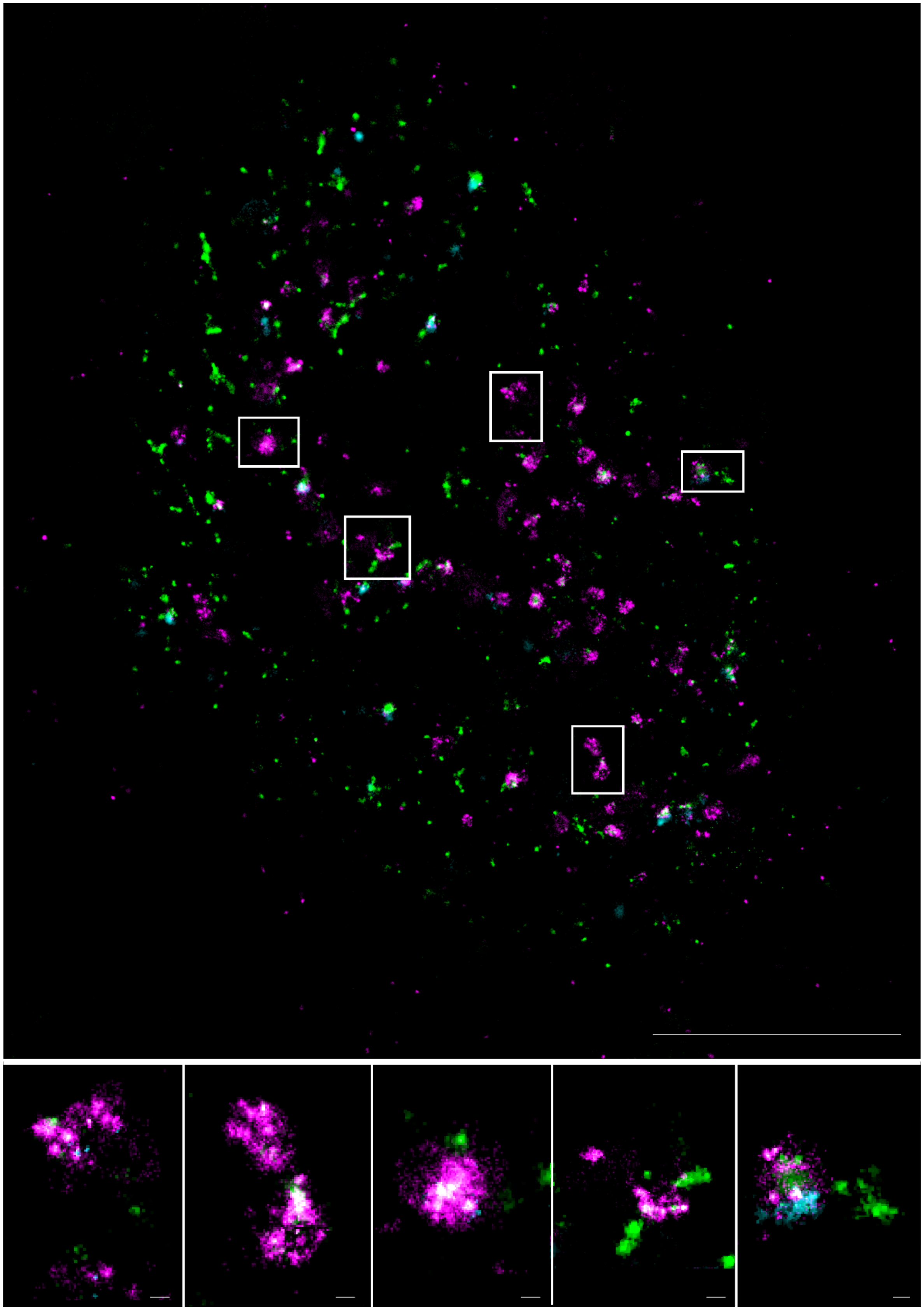

**Extended Data Figure 12.** Partial cellular overview of SMLM data in a HeLa cell with ROIs indicating endosomes presented in Figure 5b. LNP-Cy5-mRNA (magenta), Transferrin (green), EGF (cyan). Scale bars 5 µm (top), 100nm (bottom).

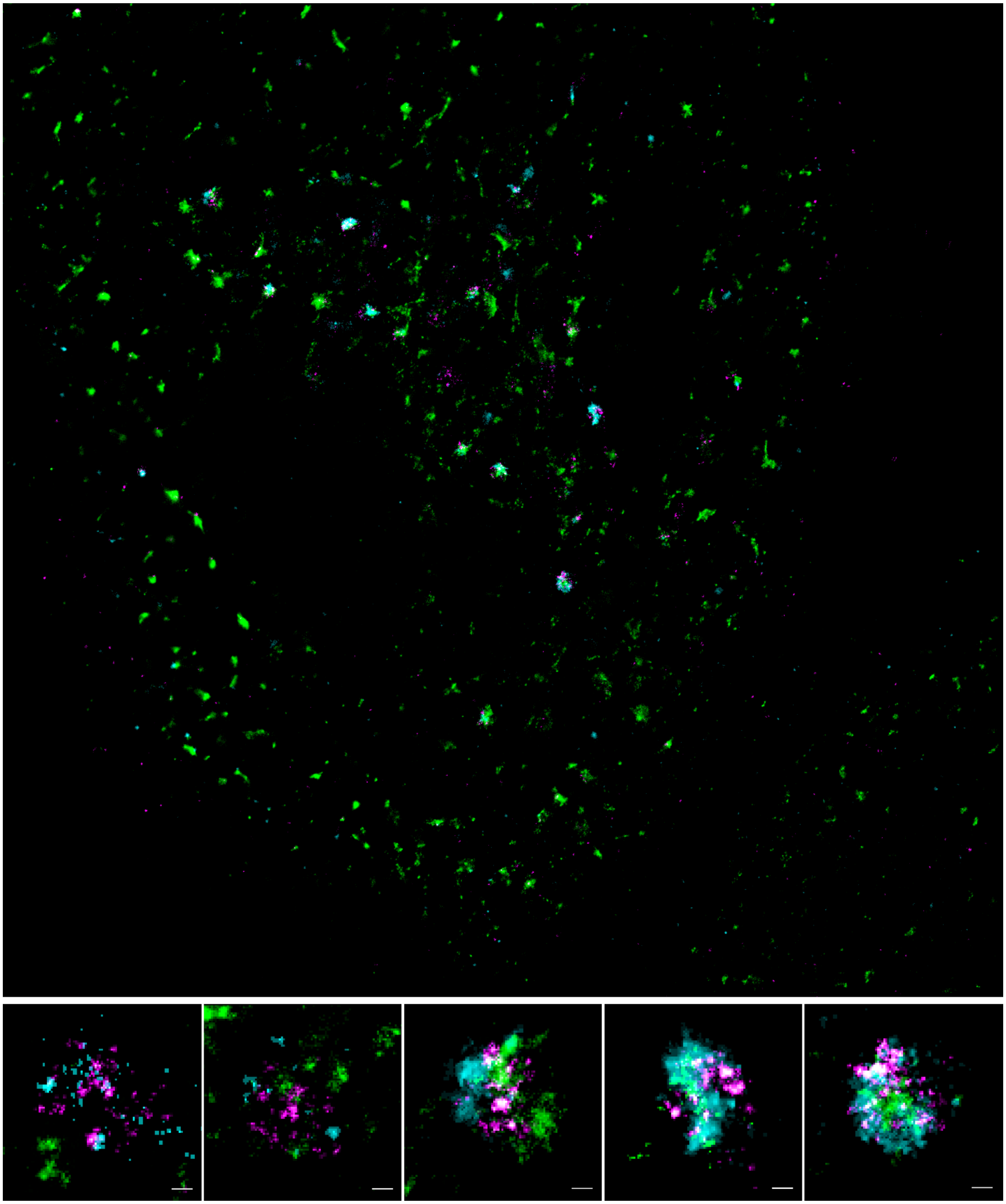

**Extended Data Figure 13.** Partial cellular overview of SMLM data in a HeLa cell with ROIs indicating endosomes presented in Figure 5c. LNP-Cy5-mRNA (magenta), Transferrin (green), EGF (cyan). Scale bars 5µm (top), 100nm (bottom).

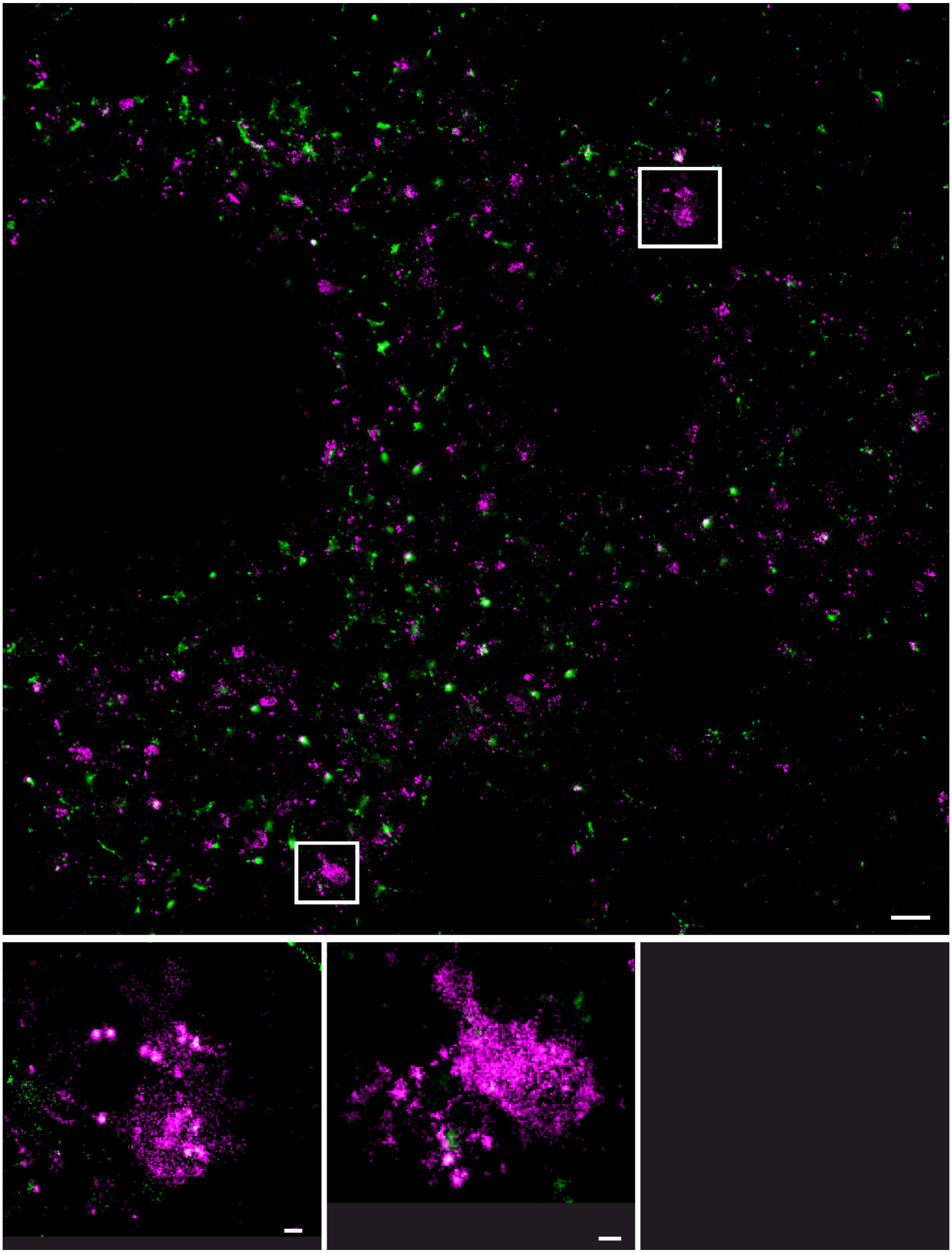

**Extended Data Figure 14.** Partial cellular overview of SMLM data in an adipocyte with ROIs indicating endosomes presented in Figure 5d. LNP-Cy5-mRNA (magenta), Transferrin (green), EGF (cyan). Scale bars 5µm (top), 100nm (bottom).

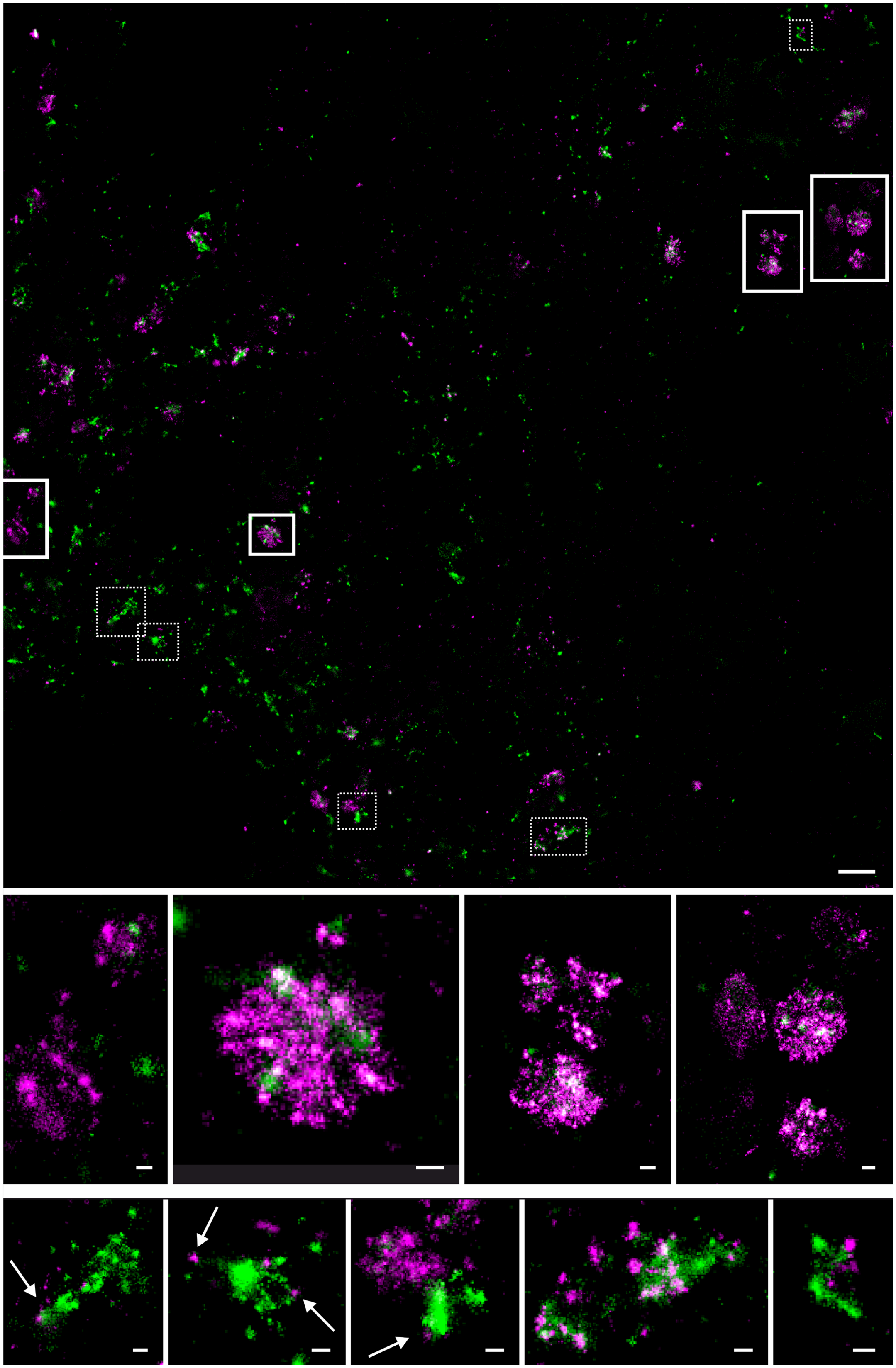

**Extended Data Figure 15.** Partial cellular overview of SMLM data in an adipocyte with ROIs indicating endosomes presented in Figure 5d and 6d. LNP-Cy5-mRNA (magenta), Transferrin (green), EGF (cyan). Scale bars 5µm (top), 100nm (bottom)

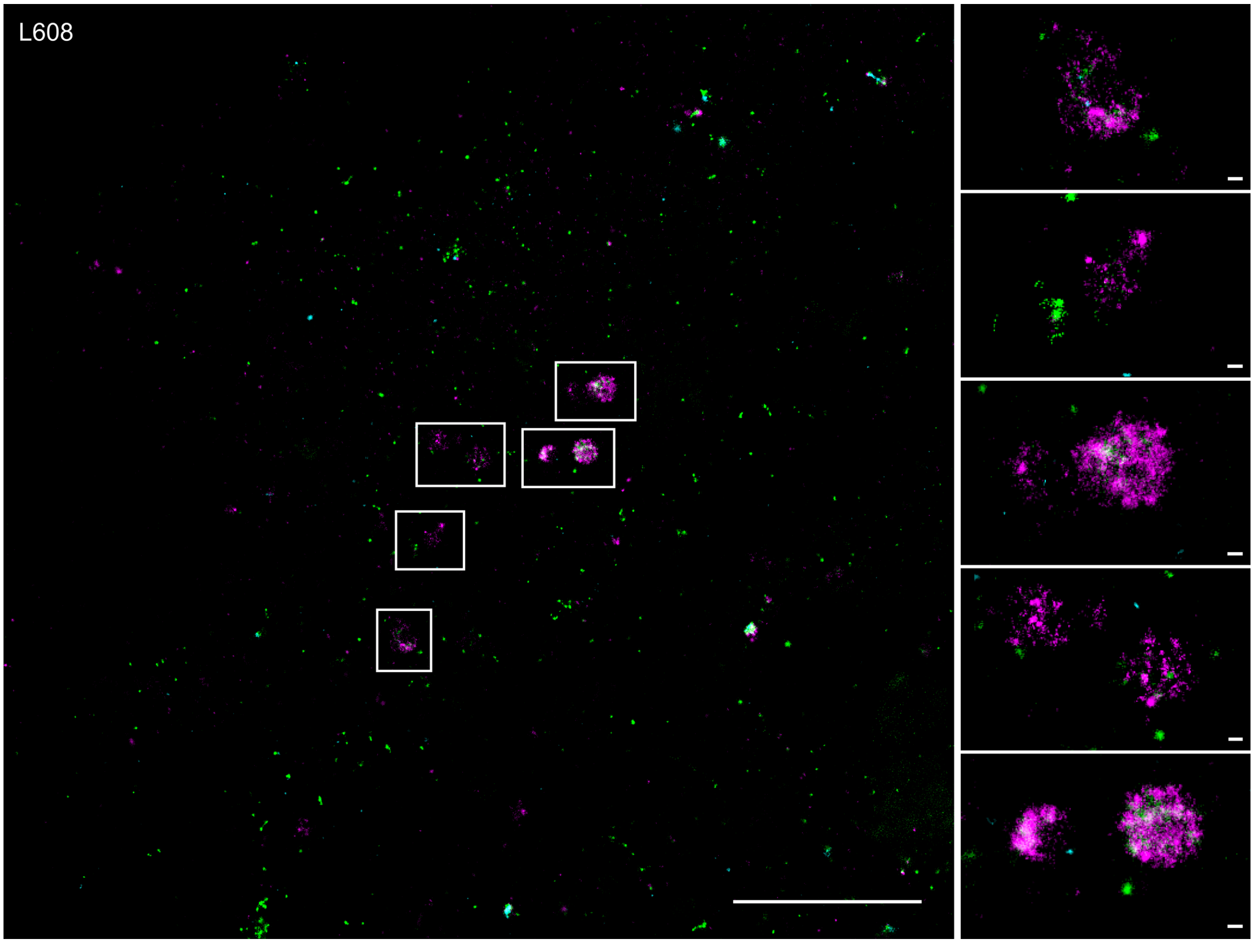


**Extended Data Figure 16.** Partial cellular overview of SMLM data in an adipocyte with ROIs indicating additional examples of ar1rested endosomes for the L608 LNP formulation. LNP-Cy5-mRNA, (magenta), Transferrin (green), EGF (cyan). Scale bars 5 µm (left), 100nm (right).

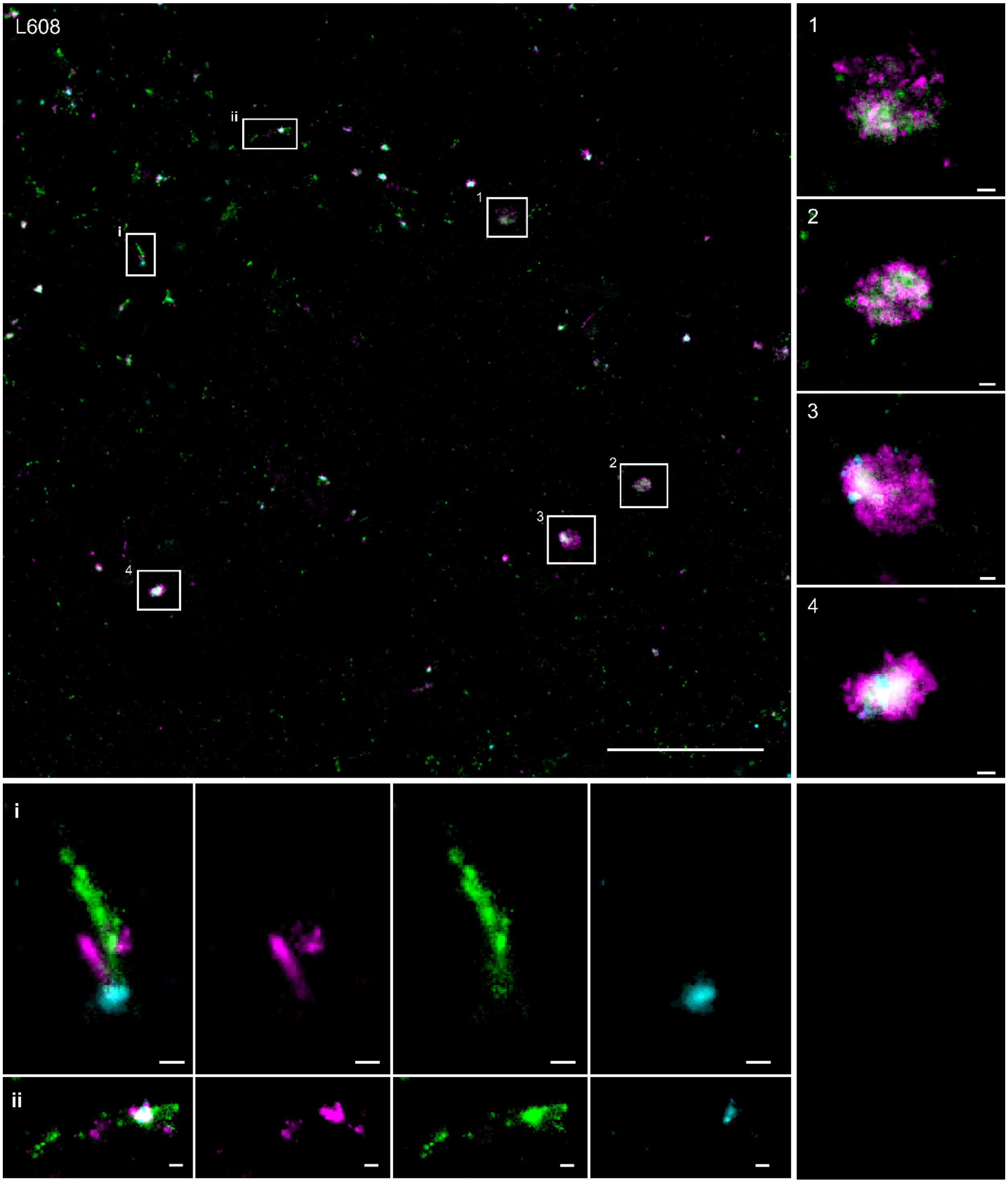


**Extended Data Figure 17.** Partial cellular overview of SMLM data in an adipocyte with ROIs indicating additional examples of arrested endosomes (right panel) and possible mRNA escape events (bottom panel) for the L608 LNP formulation. LNP-Cy5-mRNA (magenta), Transferrin (green), EGF (cyan). Scale bars 5 µm (top left), 100nm (bottom, right).

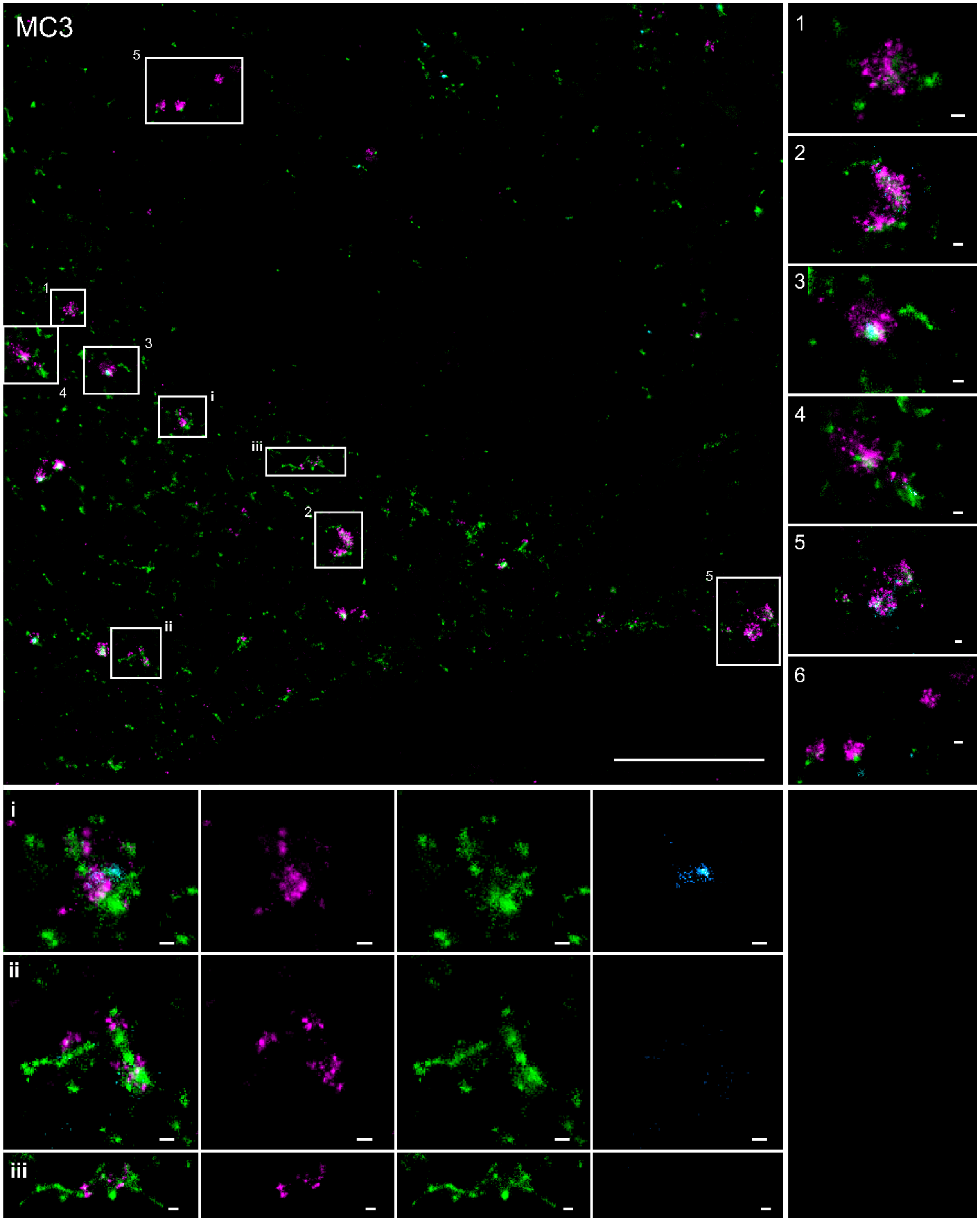


**Extended Data Figure 18.** Partial cellular overview of SMLM data in an adipocyte with ROIs indicating additional examples of arrested endosomes (right panel) and possible mRNA escape events (bottom panel) for the MC3 LNP formulation. LNP-Cy5-mRNA (magenta), Transferrin (green), EGF (cyan). Scale bars 5 µm (top left), 100nm (bottom, right).

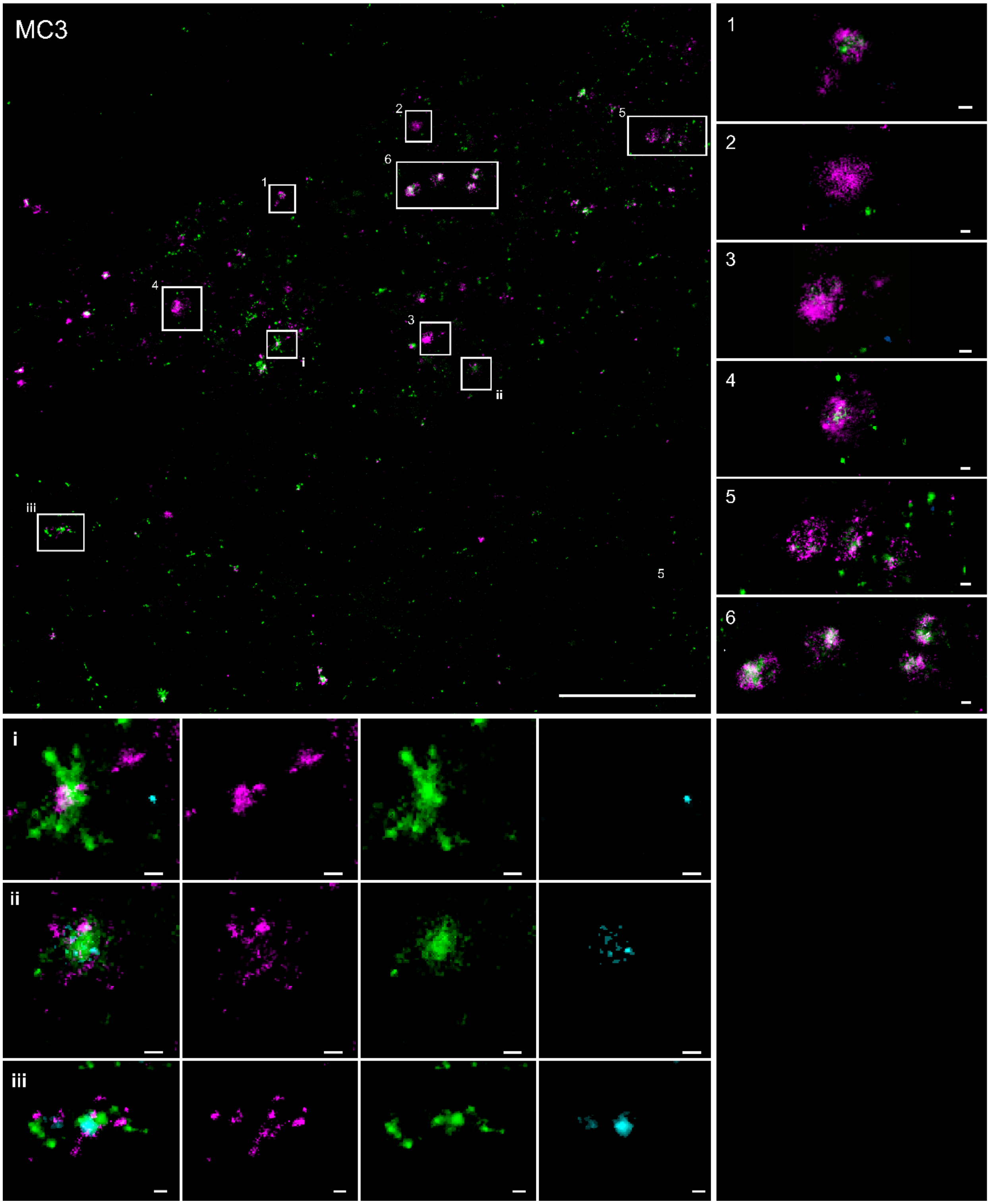


**Extended Data Figure 19.** Partial cellular overview of SMLM data in an adipocyte with ROIs indicating additional examples of arrested endosomes (right panel) and possible mRNA escape events (bottom panel) for the MC3 LNP formulation. LNP-Cy5-mRNA (magenta), Transferrin (green), EGF (cyan). Scale bars 5 µm (top left), 100nm (bottom, right).

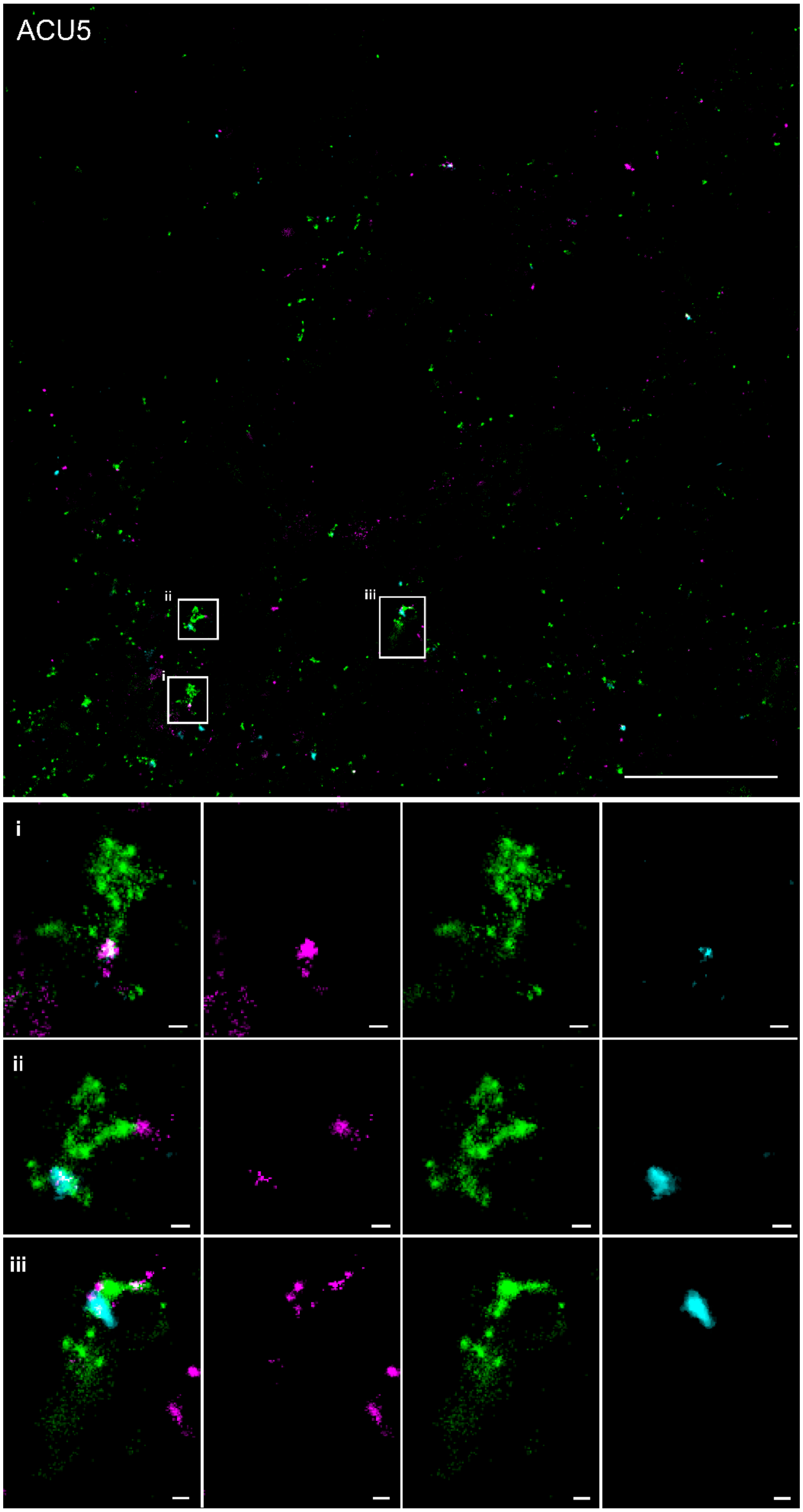


**Extended Data Figure 20.** Partial cellular overview of SMLM data in an adipocyte with ROIs indicating additional examples of possible mRNA escape events (bottom panel) for the ACU5 LNP formulation. LNP-Cy5-mRNA (magenta), Transferrin (green), EGF (cyan). Scale bars 5 µm (top), 100nm (bottom).

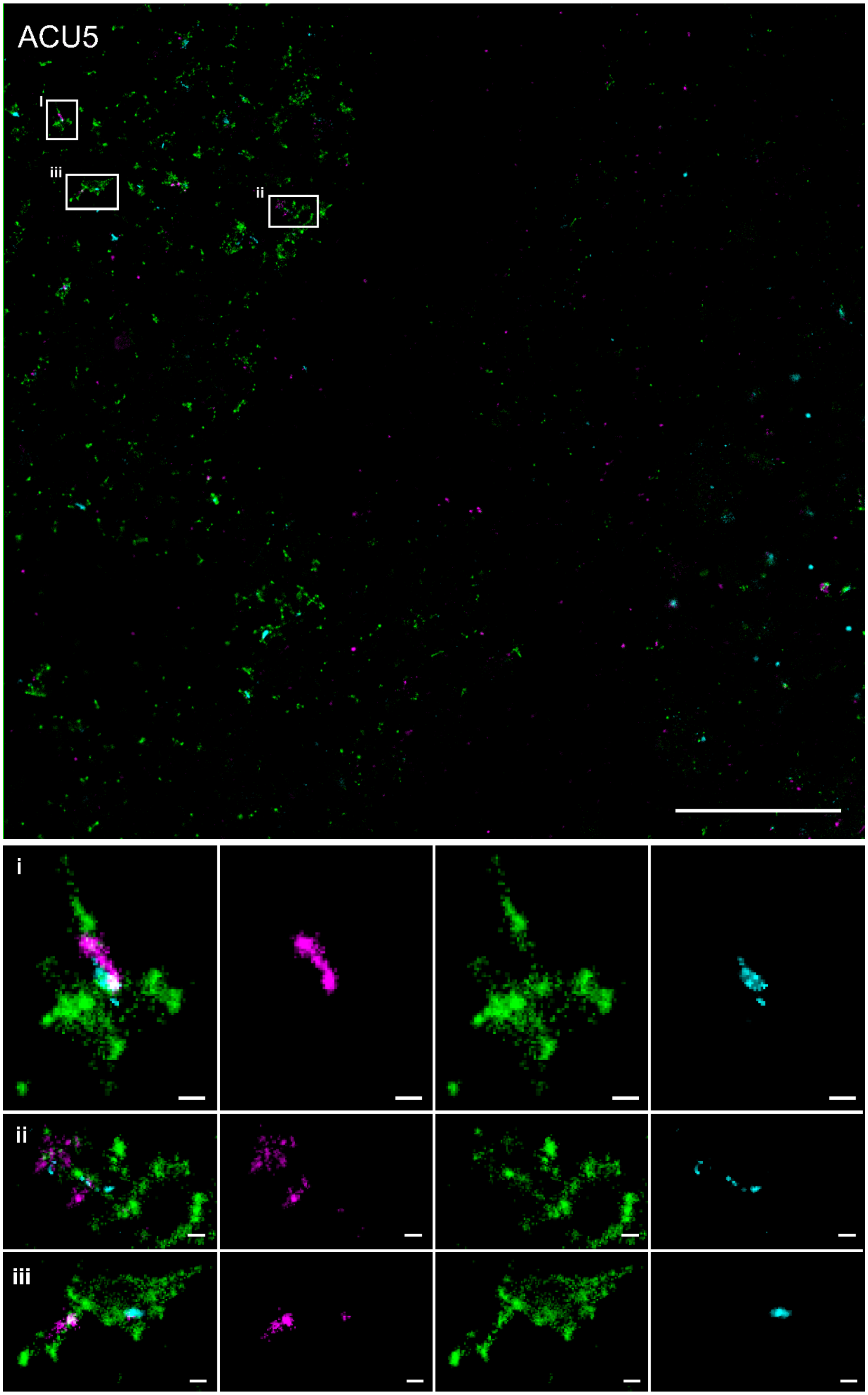


**Extended Data Figure 21.** Partial cellular overview of SMLM data in an adipocyte with ROIs indicating additional examples of possible mRNA escape events (bottom panel) for the ACU5 LNP formulation. LNP-Cy5-mRNA (magenta), Transferrin (green), EGF (cyan). Scale bars 5 µm (top), 100nm (bottom).

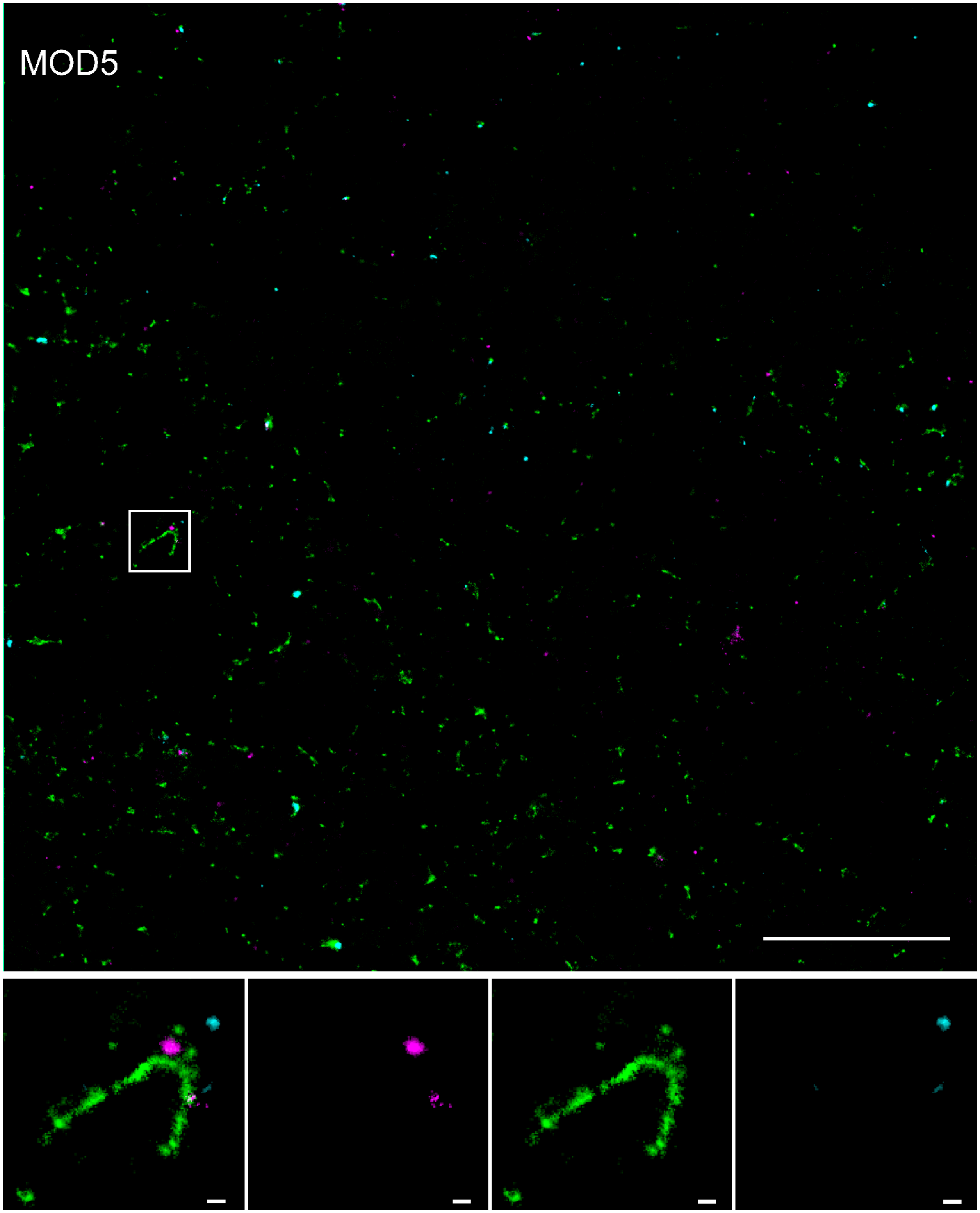


**Extended Data Figure 22.** Partial cellular overview of SMLM data in an adipocyte with ROIs indicating additional examples of possible mRNA escape events (bottom panel) for the MOD5 LNP formulation. LNP-Cy5-mRNA (magenta), Transferrin (green), EGF (cyan). Scale bars 5 µm (top), 100nm (bottom).

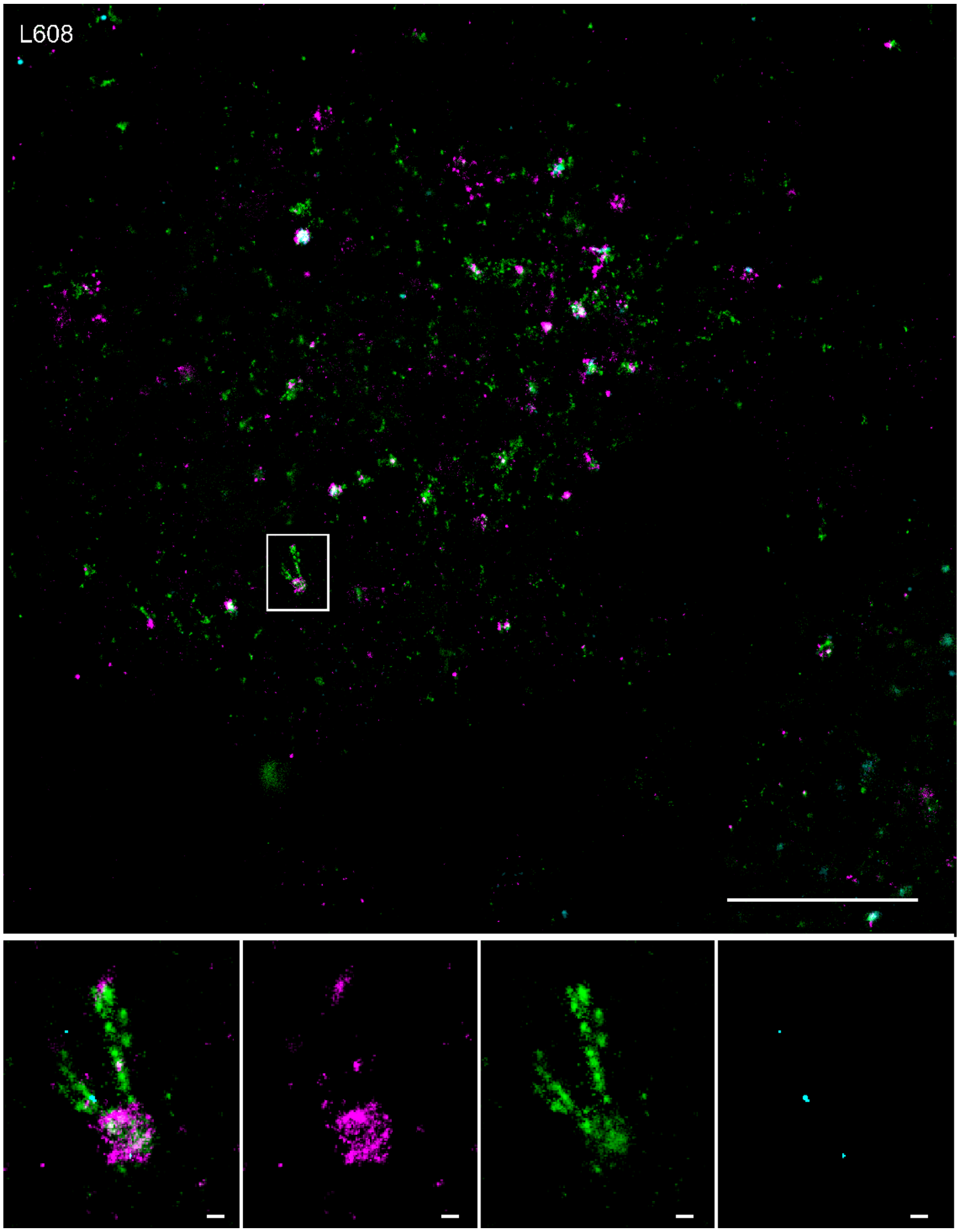


**Extended Data Figure 23.** Partial cellular overview of SMLM data in an adipocyte with ROIs indicating additional examples of possible mRNA escape events (bottom panel) for the L608 LNP formulation. LNP-Cy5-mRNA (magenta), Transferrin (green), EGF (cyan). Scale bars 5 µm (top), 100nm (bottom).

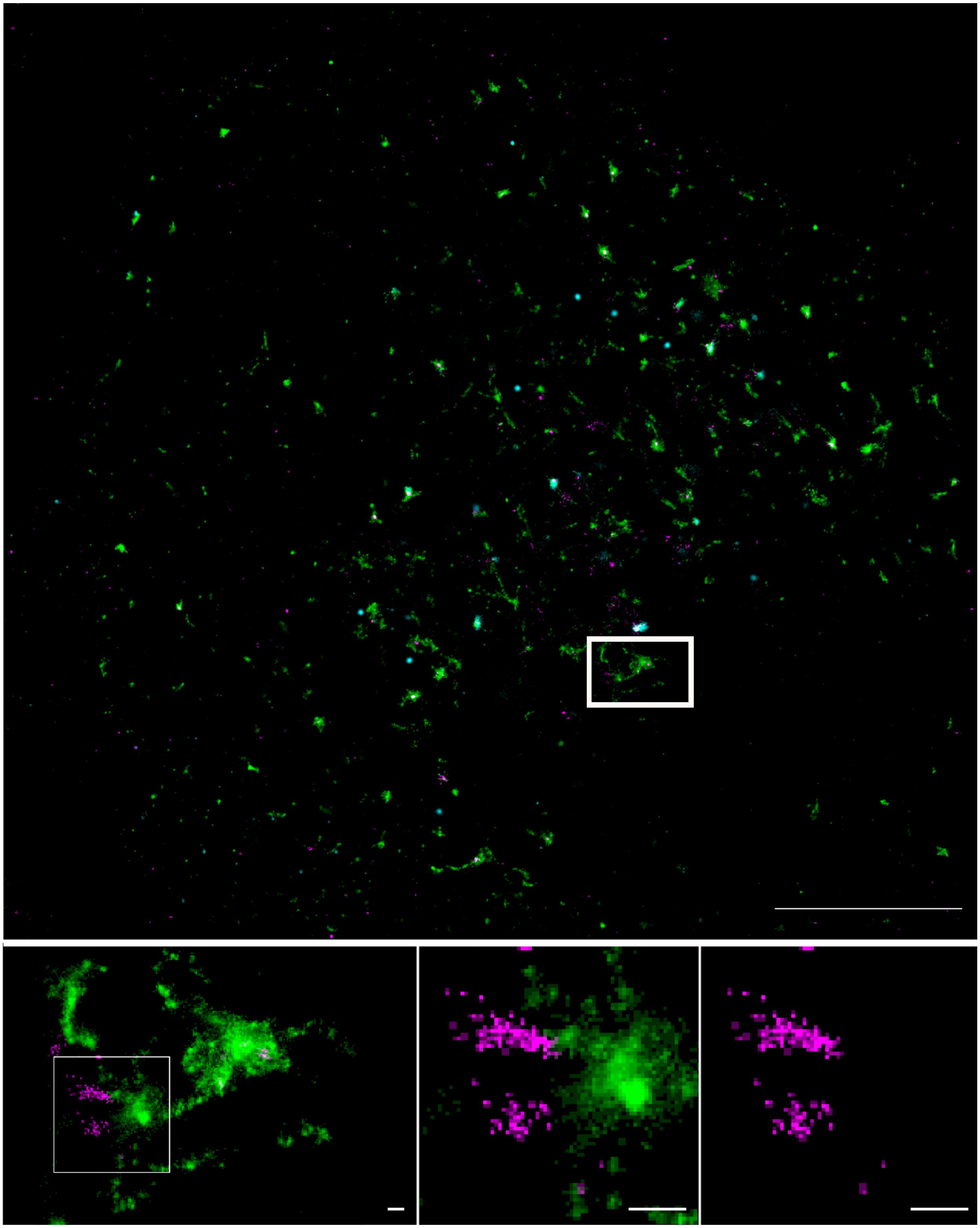

**Extended Data Figure 24.** Partial cellular overview of SMLM data in a HeLa cell with ROIs indicating the endosome presented in Figure 6b. LNP-Cy5-mRNA (magenta), Transferrin (green), EGF (cyan). Scale bars 5 µm (top), 100nm (bottom).

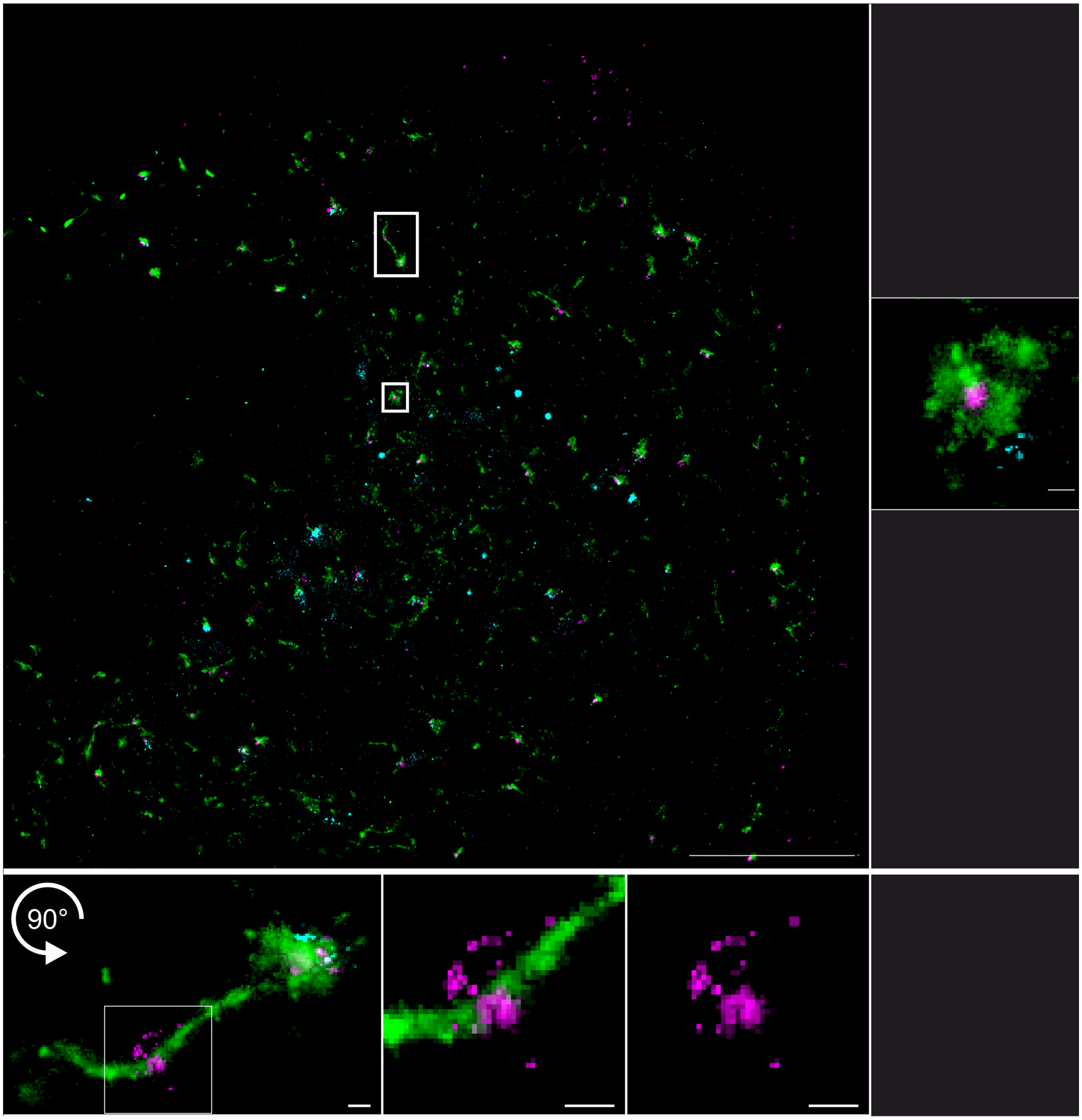


**Extended Data Figure 25.** Partial cellular overview of SMLM data in a HeLa cell with ROIs indicating the endosome presented in Figure 4c and Figure 6c. LNP-Cy5-mRNA (magenta), Transferrin (green), EGF (cyan). Scale bars 5 µm (top), 100nm (bottom).

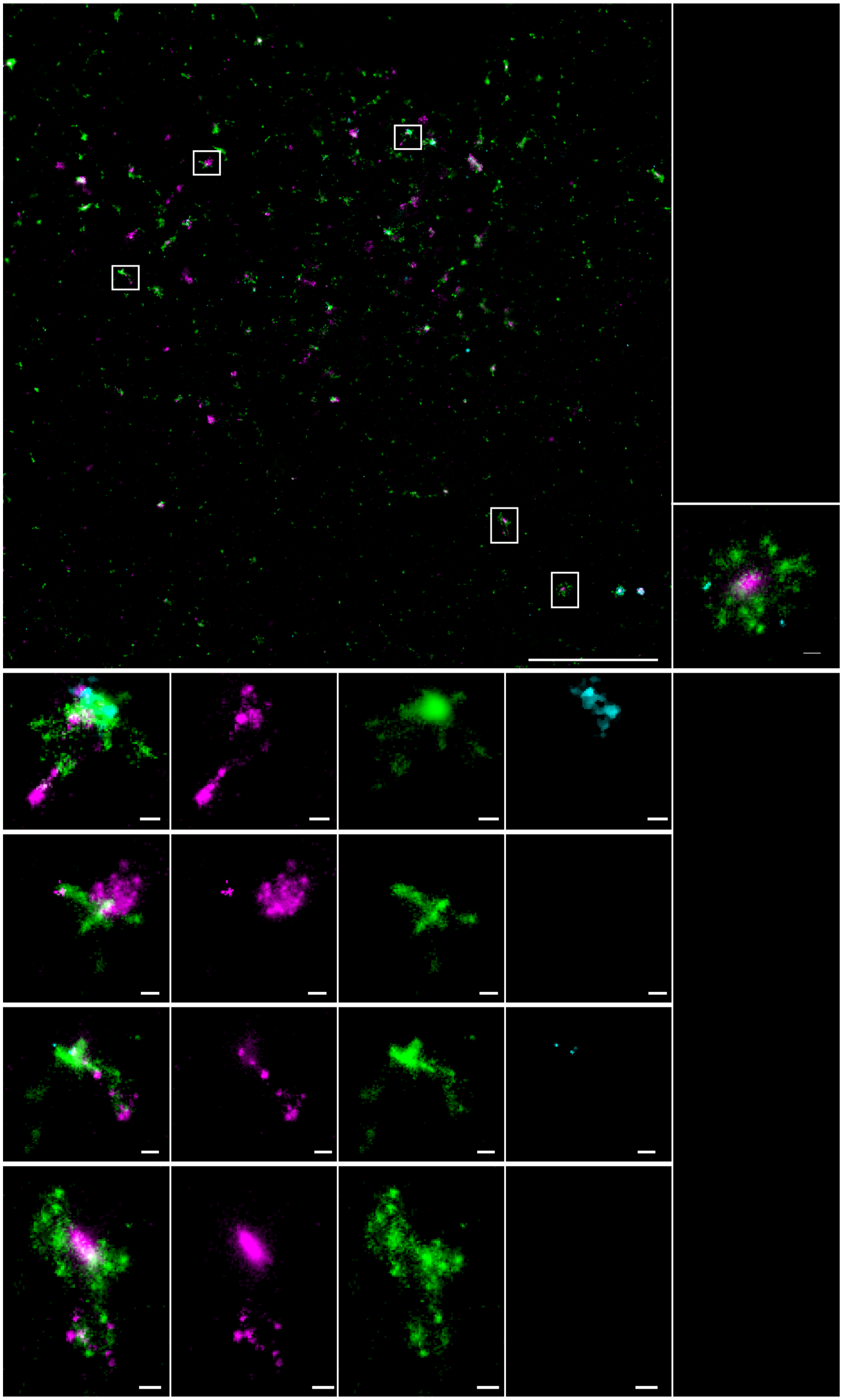

**Extended Data Figure 26.** Partial cellular overview of SMLM data in a HeLa cell with ROIs indicating additional examples of possible mRNA escape events (bottom panel) for the MC3 LNP formulation and ROI for image presented in Figure 4b. LNP-Cy5-mRNA (magenta), Transferrin (green), EGF (cyan). Scale bars 5µm (top), 100nm (bottom).

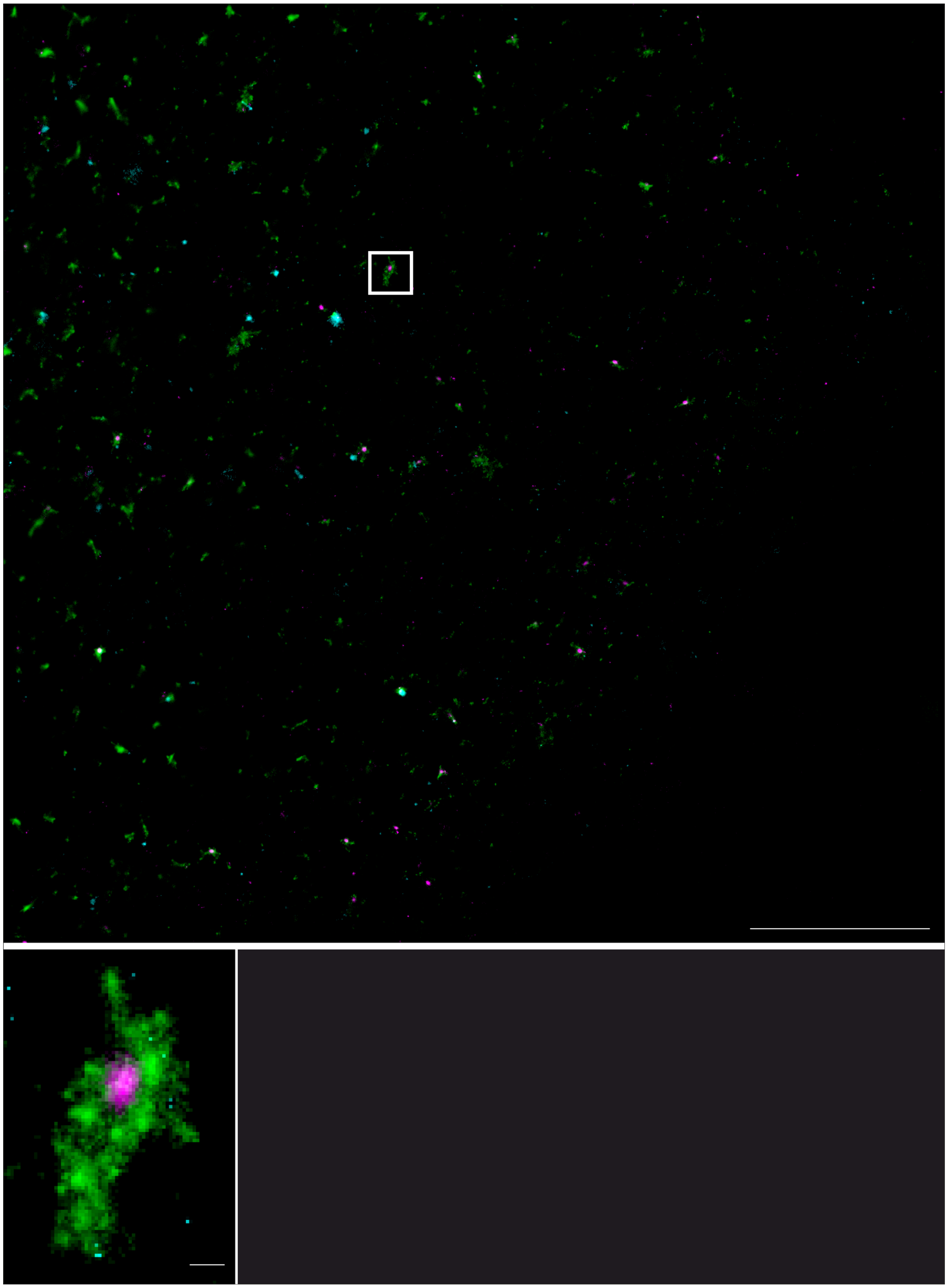

**Extended Data Figure 27:** Partial cellular overview of SMLM data in HeLa cells with ROIs indicating the endosome presented in Figure 4d. LNP-Cy5-mRNA (magenta), Transferrin (green), EGF (cyan). Scale bars 5µm (top), 100nm (bottom).

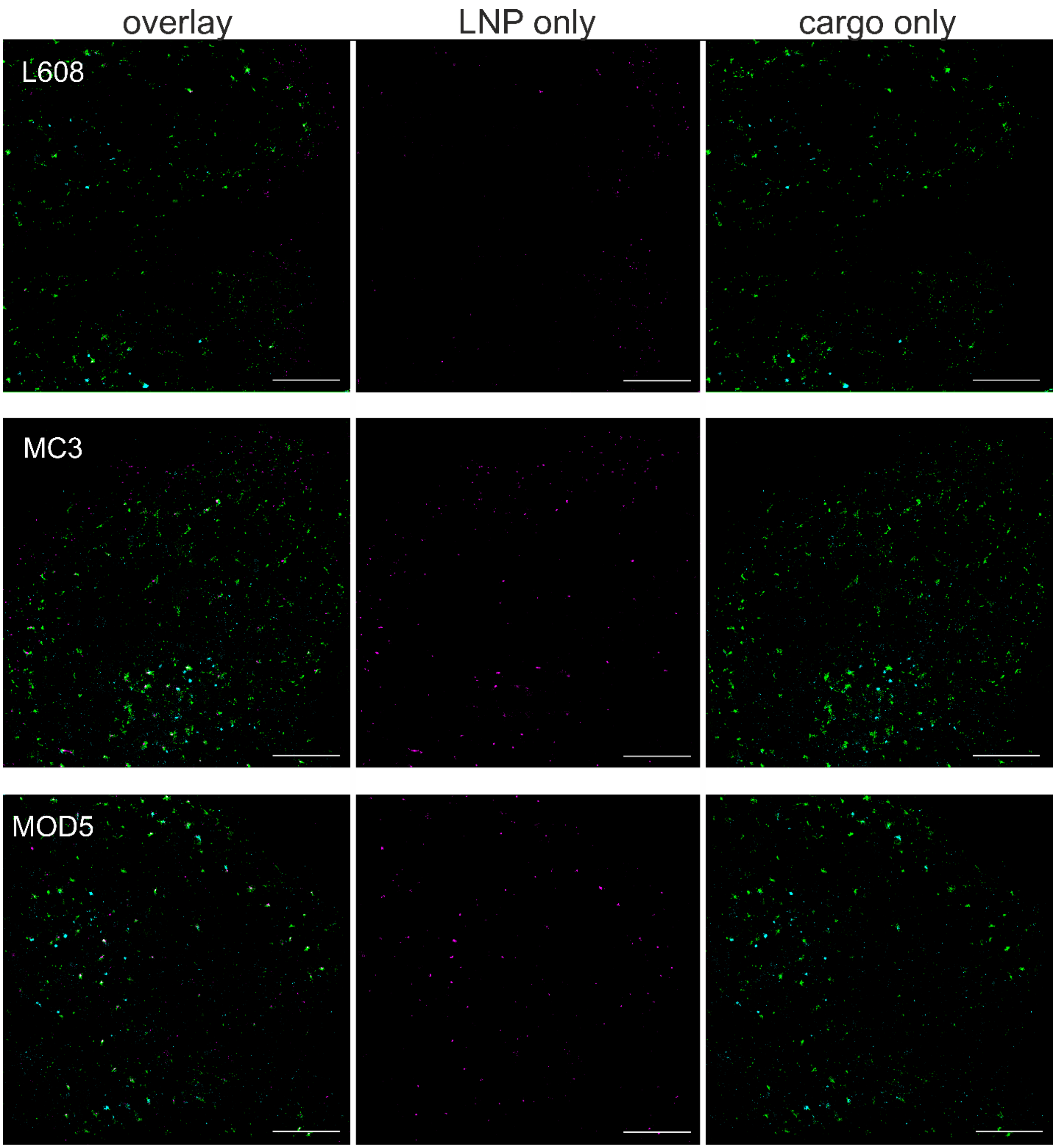


**Extended Data Figure 28**. Exemplary field of views of cells incubated with LNPs and cargo molecules simultaneously for 30 minutes. LNPs (magenta) are distributed almost exclusively at the cellular periphery, while Transferrin (green) and EGF (cyan) cargo is apparent within the entire cell. Scale bars 5µm

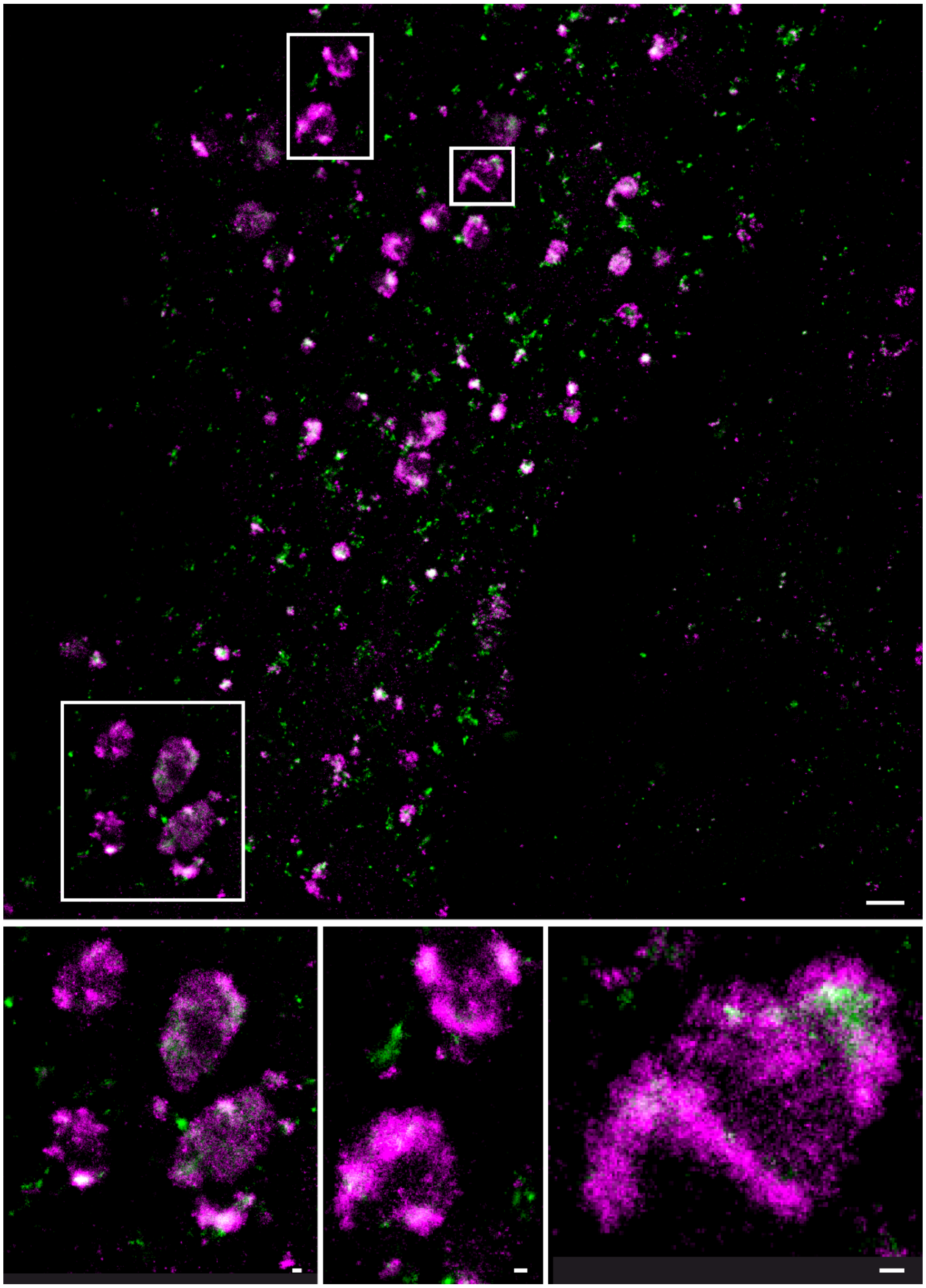

**Extended Data Figure 29:** Arrested endosomes in primary fibroblasts. *Top:* Paradigmatic overview of a fibroblast, imaged by SMLM (Tfn (green) and LNP-Cy5-mRNA (red), Scale bars 1µm). White boxes indicate exemplary arrested endosomes. *Bottom:* Zoom-ins of regions of interest for arrested endosomes. Scale bars 100 nm.

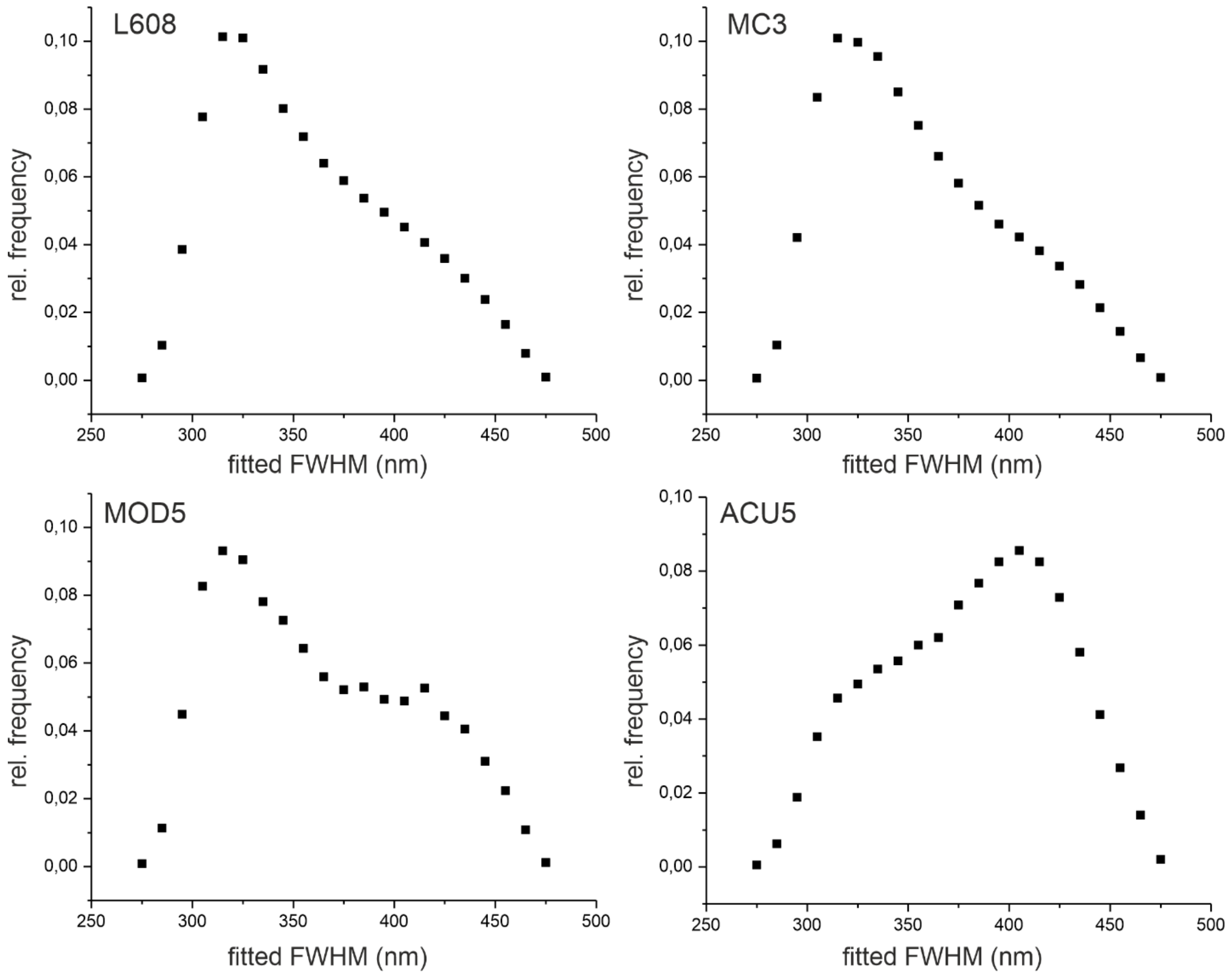


**Extended Data Figure 30.** Distribution of fitted FWHM of mRNA-Cy5 localizations. The main portion of all localizations reside in a window between 325 nm and 425 nm FWHM, which indicates an axial distribution in an approximately 500 nm.

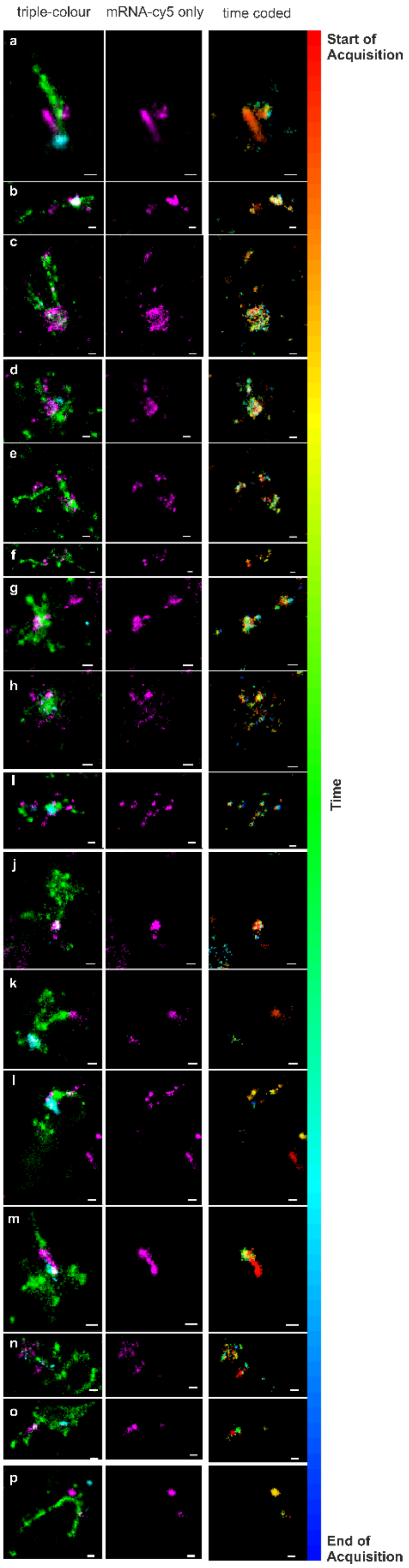


**Extended Data Figure 31**. The temporally color-coded images of mRNA escape events (right coloumn) show no sign of directed spatio-temporal displacements (drift or diffusion artefacts). Therefore, the potential escape events are genuine mRNA distributions in the sample. Scale bars 100 nm.

**L608 AZ14245376 EN11819-93-001**

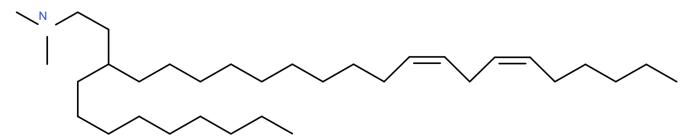


$^1$H NMR (500 MHz, $CDCl_3$) δ 5.36 (tdd, 4H), 2.77 (t, 2H), 2.39 (d, 8H), 2.05 (q, 4H), 1.50 (q, 2H), 1.19 – 1.38 (m, 39H), 0.89 (td, 6H).

LC-MS (ESI) : 476.6 $[M + H]^+$

**MC3 AZ13759693 EN08699-26-001**

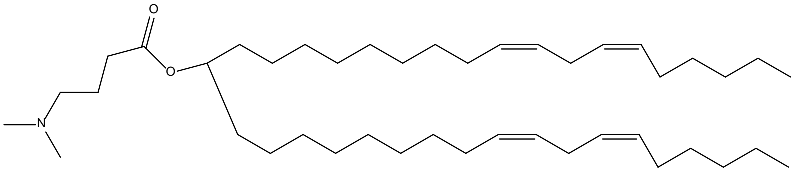


$^1$H NMR (500 MHz, $CDCl_3$) δ 5.28 – 5.43 (m, 8H), 4.74 – 4.9 (m, 1H), 2.77 (t, 4H), 2.3 – 2.37 (m, 4H), 2.26 (s, 6H), 2.04 (qd, 8H), 1.81 (p, 2H), 1.50 (d, 4H), 1.24 – 1.4 (m, 36H), 0.89 (t, 6H).

LC-MS (ESI) : 642.8 $[M + H]^+$

**ACU5 AZ13851985 EN09575-30-001**

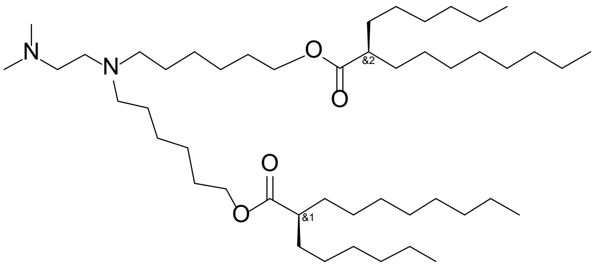


$^1$H NMR (500 MHz, $CDCl_3$) δ 4.05 (t, 4H), 2.52 – 2.58 (m, 2H), 2.40 (dt, 5H), 2.30 (tt, 2H), 2.25 (s, 6H), 1.52 – 1.66 (m, 8H), 1.22 – 1.47 (m, 57H), 0.87 (td, 12H).

LC-MS (ESI) : 765.5 $[M + H]^+$

**ACU22            AZ14028367                EN08257-22-001**

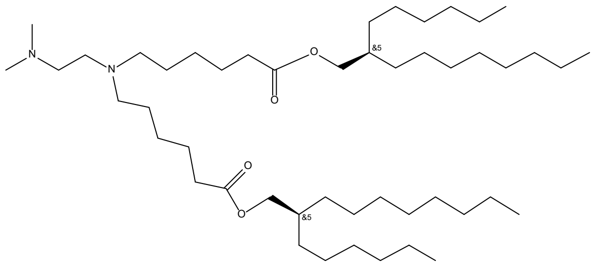


[1]H NMR (500 MHz, $CDCl_3$) δ 3.96 (d, 4H), 2.49 – 2.58 (m, 2H), 2.26 – 2.44 (m, 10H), 2.23 (s, 6H), 1.6 – 1.68 (m, 6H), 1.4 – 1.49 (m, 4H), 1.24 – 1.35 (m, 52H), 0.88 (t, 12H).

LC-MS (ESI) : 765.8  $[M + H]^+$

**Mod5 (SM86)  AZ14118365                EN08699-25-001**

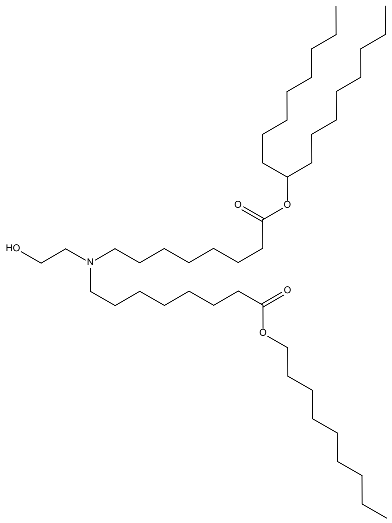


[1]H NMR (500 MHz, $CDCl_3$) δ 4.86 (p, 1H), 4.05 (t, 2H), 3.57 (t, 2H), 2.63 (s, 2H), 2.50 (s, 4H), 2.28 (td, 4H), 1.62 (q, 6H), 1.49 (dq, 8H), 1.18 – 1.34 (m, 49H), 0.88 (td, 9H).

LC-MS (ESI) : 710.8  $[M + H]^+$

**L319 AZ13897989 CRO-TI-CVO-370-1-012-01**

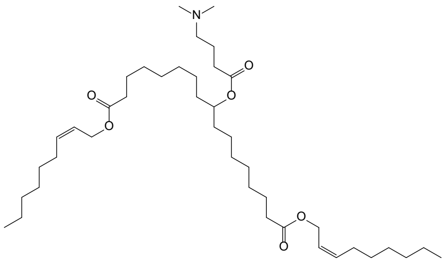


[1]H NMR (400 MHz, DMSO-d6) δ 0.86 (t, 6H), 1.05 – 1.38 (m, 31H), 1.40 – 1.58 (m, 7H), 1.66 (p, 2H), 2.07 (q, 4H), 2.16 (s, 5H), 2.28 (q, 7H), 4.56 (d, 4H), 4.73 – 4.83 (m, 1H), 5.44 – 5.57 (m, 2H), 5.57 – 5.70 (m, 2H), 8.18 (s, 1H)

LC-MS (ESI) : 678.6 $[M + H]^+$

**Extended Data Figure 32:** NMR characterization data of cationic lipids. (Page 33-35).

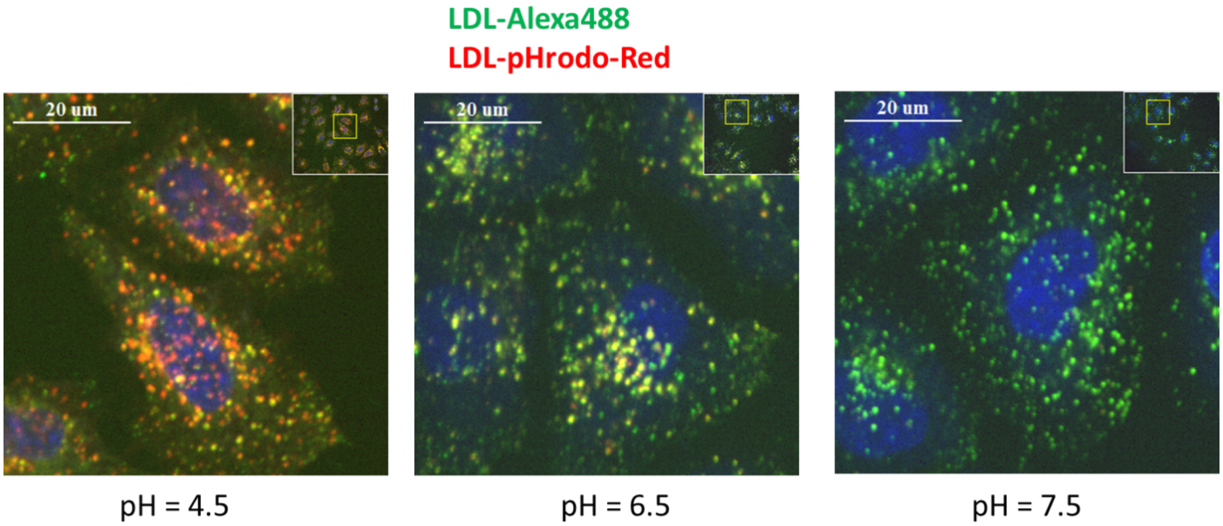


**Extended Data Figure 33.** Representative images of co-internalized mixture of LDL-pHRodo-Red/LDL-Alexa-488. Cells were fixed and imaged in calibrated buffers with pH 4.5, 6.5 and 7.5.

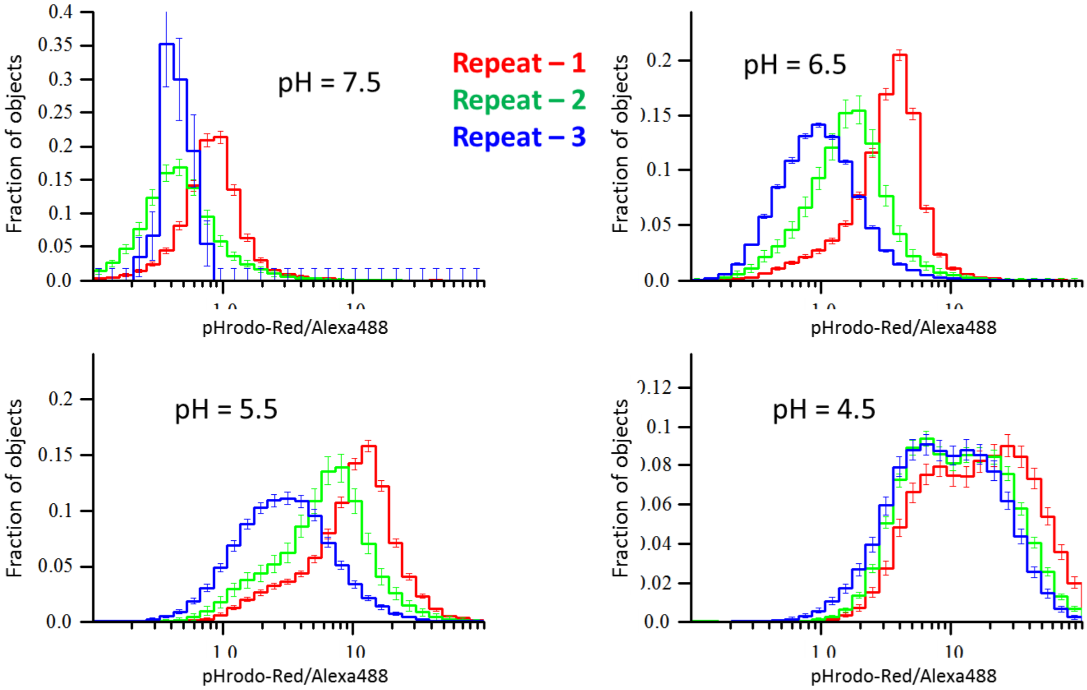


**Extended Data Figure 34.** Distribution ratios of integral intensities of pHRodo-Red/Alexa-488 in calibration measurements in three independent repeats of experiment. The pH of calibration buffer is denoted on respective panels. Colors of curves denote experiment repeats.

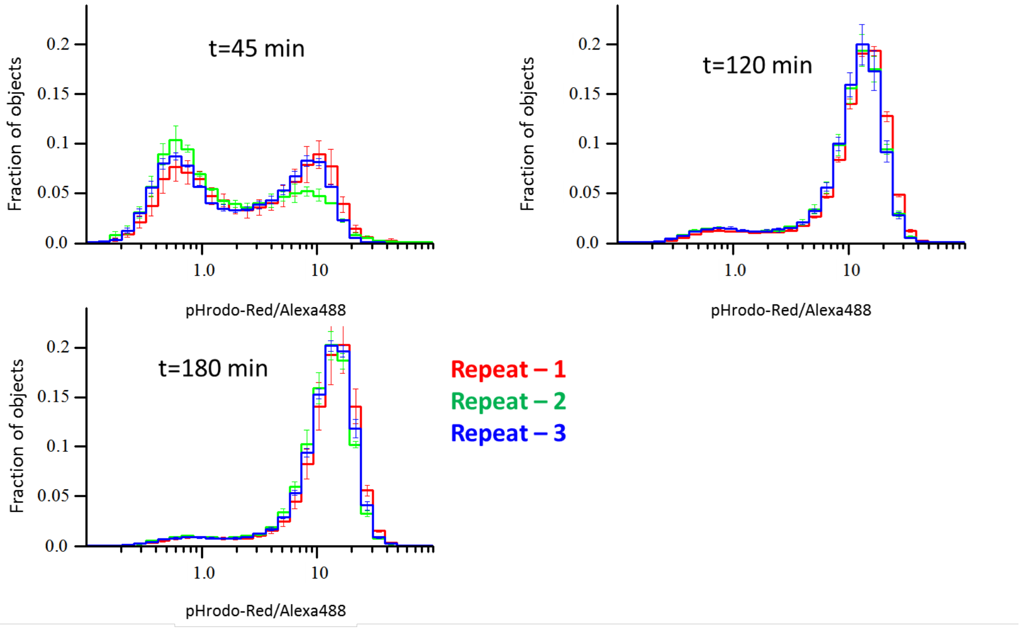


**Extended Data Figure 35.** Distribution of objects ratios of integral intensities of pHRodo-Red/Alexa-488 in live HeLa cells internalized with LDL-pHRodo-Red and LDL-Alexa-488 for the time period denoted on respective panels. Colors of curves denote repeat number of experiments.

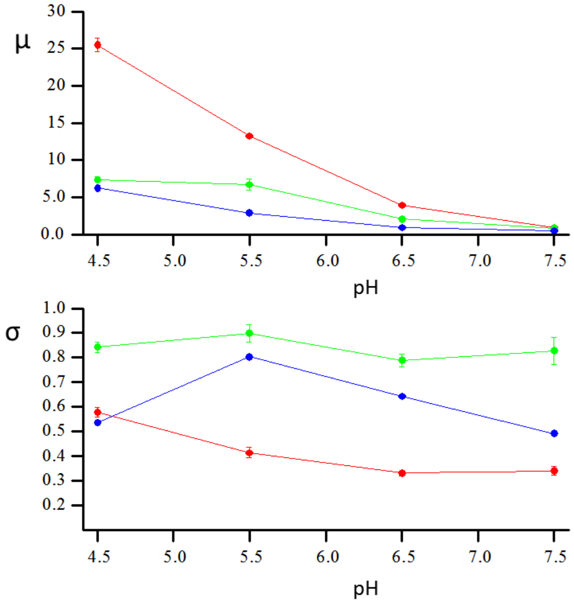


**Extended Data Figure 36:** pH dependency of parameters $\mu$ and $\sigma$ of log-normal components. Red, green and blue colors denote components with largest, middle and smallest ratios $\mu$. The global fit was performed by global histogram fit procedure in GraphView application of MotionTracking software.

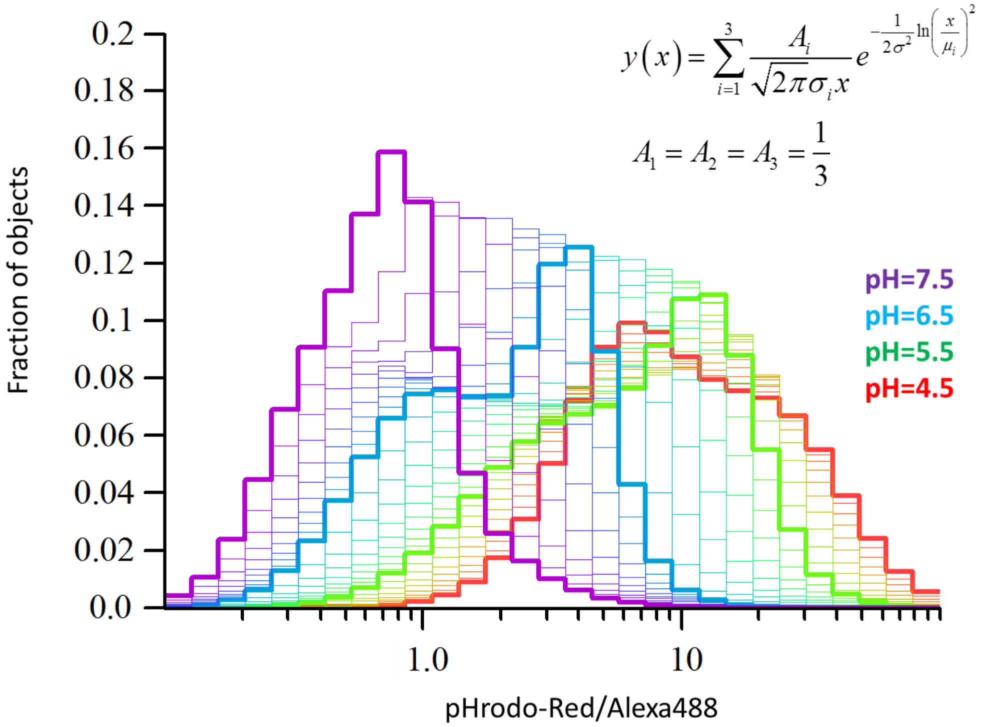


**Extended Data Figure 37:** Predicted distribution of ratios with equal contributions components at pH in range from 4.5 to 7.5.

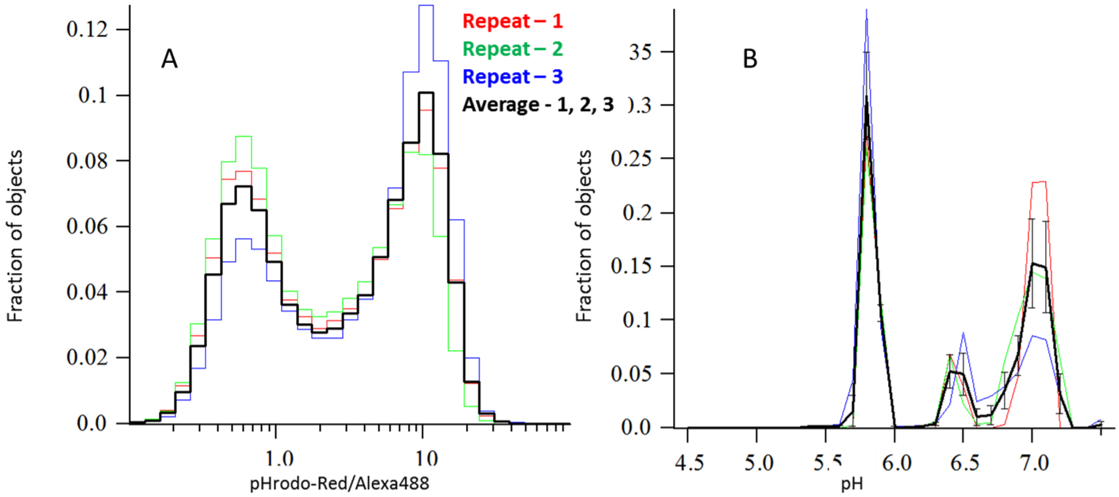


**Extended Data Figure. 38**. **A**. Experimental distribution of intensities ratios (t=45min). **B.** Fitted distribution of pH. Colors encode experimental repeats. Black line is average of repeats. Error bars on average curve are standard error of means (SEM).

| CIL | <Z> (nm) | PDI | <N> (nm) | Encapsulation (%) |
|---|---|---|---|---|
| **L608** | 82 ± 3 | 0.03 ± 0.01 | 68 ± 4 | 97 ± 1 |
| **MC3** | 81 ± 4 | 0.03 ± 0.01 | 67 ± 4 | 98 ± 1 |
| **ACU5** | 82 ± 4 | 0.03 ± 0.01 | 67 ± 5 | 97 ± 1 |
| **ACU22** | 72 ± 6 | 0.08 ± 0.02 | 54 ± 7 | 96 ± 2 |
| **MOD5** | 75 ± 4 | 0.03 ± 0.01 | 62 ± 7 | 96 ± 1 |
| **L319** | 90 ± 2 | 0.06 ± 0.02 | 73 ± 6 | 88 ± 5 |

<Z>: Intensity-averaged size, PDI: polydispersity index, <N>: Number-averaged size

**Supplementary Table 1: LNPs size and encapsulation efficiency calculated by DLS and Ribo green assays**

| Endosomes with | Average pH | % of endosomes |
|---|---|---|
| **LDL probes alone** | 5.56 | 78 |
| | 6.5 | 5 |
| | 7 | 12 |
| | Other pH values close to background | 5 |
| **L608** | 5.12 | 16 |
| | 6.06 | 46 |
| | 6.49 | 20 |
| | Other pH values close to background | 18 |
| **MC3** | 4.99 | 6 |
| | 6.05 | 58 |
| | 6.5 | 13 |
| | Other pH values close to background | 23 |
| **ACU5** | 5.95 | 73 |
| | 6.52 | 8 |
| | Other pH values close to background | 19 |
| **ACU22** | 5.88 | 77 |
| | 6.5 | 7 |
| | Other pH values close to background | 16 |
| **MOD5** | 5.78 | 83 |
| | 6.56 | 2 |
| | Other pH values close to background | 15 |
| **L319** | 5.6 | 78 |
| | 6.02 | 3 |
| | 6.87 | 2 |
| | Other pH values close to background | 17 |

**Supplementary Table 2: Percentage of LNP-Cy5-mRNA containing endosomes with indicated pH.** HeLa cells were incubated with LNP-Cy5-mRNA and LDL pH probes for 2h and pH of the LNP-mRNA containing endosomes were calculated **(see Methods)**. The percentage of endosomes with an average pH of characteristic late endosomes in control (LDL alone) is shown in green shading. The control cells (LDL alone) have 5% of LDL-positive objects with pH of 6.5 and 74% a pH of 5.5, characteristic of early and late endosomes, respectively The rest (21%) of endosomes displayed pH values ranging between neutral (endocytic vesicles, e.g. Clathrin-coated vesicles) and intermediate acidic values. The percentage of arrested endosomes with a pH values between late and early endosomes are shown in red color shade.

| Endosomes with | Average pH | % of endosomes |
|---|---|---|
| **LDL alone** | 5.52 | 74 |
| | 6.48 | 5 |
| | 7 | 12 |
| | Other pH values close to background | 5 |
| **L608** | 5.8 | 13 |
| | 6.16 | 49 |
| | 6.52 | 15 |
| | 6.92 | 19 |
| | Other pH values close to background | 18 |
| **MC3** | 6.1 | 17 |
| | 6.3 | 48 |
| | 6.54 | 10 |
| | 6.9 | 12 |
| | Other pH values close to background | 13 |
| **ACU5** | 6.1 | 89 |
| | 6.5 | 5 |
| | Other pH values close to background | 6 |
| **ACU22** | 5.95 | 76 |
| | 6.5 | 13 |
| | Other pH values close to background | 11 |
| **MOD5** | 5.9 | 92 |
| | 6.49 | 5 |
| | Other pH values close to background | 3 |
| **L319** | 5.5 | 91 |
| | 6.02 | 2 |
| | Other pH values close to background | 7 |

**Supplementary Table 3: Percentage of LNP-Cy5-mRNA containing endosomes with indicated pH**. LNP-Cy5-mRNA and LDL pH probes where incubated for 3h in HeLa cells and pH of LNP-Cy5-mRNA containing endosomes were calculated **(see Methods)**. The percentage of endosomes with an average pH of characteristic late endosomes in control (LDL alone) is shown in green shading. The percentage of arrested endosomes with a pH values between late and early endosomes are shown in red color shade.

| Antibody name | Company | Product number | Antibody dilution used |
|---|---|---|---|
| **LAMP1** | BD Bioscience | 555798 | 1 to 200 |
| **APPL1** | Produced at Eurogentec, Belgium [6]. Purified at MPI-CBG | α APPL1 2624-3 | 1 to 250 |
| **EEA1** | Produced at EMBL [7]. Purified at MPI-CBG | α EEA1 f.1 Ø7JF | 1 to 1000 |
| **Rab11** | Invitrogen (Thermo Fisher) | 71-5300 | 1 to 250 |
| **RAB5** | BD Transduction Laboratories Bioimaging | 610725 | 1 to 100 |
| **ANKFY1** | Sigma Aldrich | SAB1401696-50UG | 1 to 100 |
| **LC3** | MBL | M152-3 | 1 to 500 |
| **LBPA** | Echelon Biosciences/ MoBiTec | Z-SLBPA | 1 to 100 |
| **CAV1** | Cell Signaling | #3238 | 1 to 100 |

**Supplementary Table 4: Details of antibodies and their dilutions used in this study.**
